# Transcriptomics of mussel transmissible cancer MtrBTN2 suggests accumulation of multiple cancerous traits and oncogenic pathways shared among bilaterians

**DOI:** 10.1101/2023.01.03.522559

**Authors:** E.A.V. Burioli, M. Hammel, E. Vignal, J. Vidal-Dupiol, G. Mitta, F. Thomas, N. Bierne, D. Destoumieux-Garzón, G.M. Charrière

## Abstract

Transmissible cancer cell lines are rare biological entities giving rise to diseases at the crossroads of cancer and parasitic diseases. These malignant cells have acquired the amazing capacity to spread from host to host. They have been described only in dogs, Tasmanian devils and marine bivalves. The *Mytilus trossulus* Bivalve Transmissible Neoplasia 2 (MtrBTN2) lineage has even acquired the capacity to spread inter-specifically between marine mussels of the *Mytilus edulis* complex worldwide. To identify the oncogenic processes underpinning the biology of these atypical cancers we performed transcriptomics of MtrBTN2 cells. Differential expression, enrichment, protein-protein interaction network, and targeted analyses were used. Overall, our results suggest the accumulation of multiple cancerous traits that way be linked to the long-term evolution of MtrBTN2. We also highlight that vertebrate and lophotrochozoan cancers could share a large panel of common drivers, which supports the hypothesis of an ancient origin of oncogenic processes in bilaterians.

## 1. Introduction

Transmissible cancers are rare and fascinating biological entities because they have evolved the ability to overcome the physical and immunological barriers of the host to become contagious and to spread between animals by direct transfer of cancer cells, thus behaving like parasites (1,2). They have been reported in only two vertebrate species so far, in dogs and in Tasmanian devils. The transmissibility of cancer cells was first demonstrated in the Canine Transmissible Veneral Tumor (CTVT) (3,4), which is thought to have originated more than 4,000 years ago (5). CTVT has since spread across continents, probably together with humans, through coitus and oral contact between dogs. Tasmanian Devil Facial Tumor Disease (DFTD) is the other well-known transmissible cancer that affects vertebrates. Two independent cancerous lineages DFT1 and DFT2 have been described to date and have emerged less than fifty years ago (6). Infection occurs via bites, a common behaviour in Tasmanian devil social interactions (7–9). As a result, cancer transmission has devastated much of the devil population, with some local populations declining by as much as 80% in the geographical areas most affected by DFTD (10–13). However, the highest number of transmissible cancer lineages identified to date has been found in bivalve molluscs (8 different lineages) and these multiple lineages have been grouped under the term Bivalve Transmissible Neoplasia (BTN) (14–17). BTNs usually transmit between individuals of the same species, but some lineages have crossed the species barrier and are circulating in populations of related host species (15–17). Indeed, one specific lineage of BTN, called *Mytilus trossulus* Bivalve Transmissible Neoplasia lineage 2 (MtrBTN2), has emerged in a *M. trossulus* founder host and has since spread to other *Mytilus* species populations (*M. trossulus*, *M. edulis*, *M. galloprovincialis*, and *M. chilensis*) across several continents—South America, Asia, and Europe (15,16,18–21).

Transmissible cancer lineages must have emerged from a first neoplastic transformation in a founder host and then evolved specific phenotypes to become a new type of contagious etiologic agent. In fact, we have previously shown that conventional non-transmissible cancers also occur in *Mytilus* bivalves (20). MtrBTN2 cells are found circulating in the haemolymph together with haemocytes, the mussel’s immune cells. Like haemocytes, they are able to infiltrate connective tissues of various organs (19,22). As the disease progresses, MtrBTN2 cells outgrow the host cells and progressively replace almost all the circulating cells present in the haemolymph and disseminate in the tissues. They have a characteristic morphology of rounded and basophilic cells with a high nucleus-to-cell ratio and they are polyploid (19). Therefore, like other haemolymphatic cancers (20), they are easily diagnosed by histological/cytological observation or flow cytometry of haemolymph (19,23). This initial diagnosis needs to be complemented by genetic analysis to distinguish MtrBTN2 from other transmissible cancers and conventional haemolymphatic cancers (16,24). MtrBTN2 cells proliferate very quickly with a mean doubling time ofll∼ll3 days (24). In addition, BTNs must be able to survive in the extra-host environment long enough to infect a new host (24,27). We have shown that MtrBTN2 cells can survive at least for 3 days without mortality and up to 8 days in seawater (24). However, the molecular pathways related to the malignancy of BTNs have not been characterised. Based on the phenotypic traits observed in MtrBTN2 cells, such as high proliferative activity, genomic abnormalities such as polyploidy, supermetastatic ability, and extended cell survival capabilities, it is likely that multiple molecular pathways contribute to the malignancy and long-term persistence of this lineage.

Among the most common and most studied functional capabilities observed in human cancer cells are the sustaining of proliferative signalling, the evasion from growth suppressors, the resistance to cell death, the replicative immortality, the activation of invasion and metastasis, the reprogramming of cellular metabolism, and the avoidance of immune destruction (25). The acquisition of these capabilities is ensured by the activation or inactivation of specific signalling pathways. Among these, Sanchez-Vega et al. (26) highlighted ten oncogenic signalling pathways that are most frequently altered in most human cancers (HIPPO, MYC, NOTCH, NRF2, PI3K, RTK/RAS, TGFβ, P53, WNT, and cell cycle), and are mainly involved in promoting cell proliferation. These conclusions were drawn from extensive data on hundreds of human cancers obtained by transcriptomic, genomic and epigenomic analyses.

Measuring relative levels of gene expression has been a key approach to identify genes and biological pathways associated with the cancerous process and functional adaptations of cancers (28). In recent years, RNA sequencing (RNA-seq) has emerged as a fast, cost-effective, and robust technology to address various biological questions (29,30). Transcriptome analyses allow to link cellular phenotypes to their molecular underpinnings. In the context of cancers, these links provide an opportunity to dissect the complexity of the cancer biological adaptations. For non-model organisms and in the absence of a suitable reference genome, as is the case for *M. trossulus*, RNA-seq is used to reconstruct and quantify *de novo* transcriptomes (31). Differential gene expression analysis is then carried out to compare the effect of treatments or conditions on gene expression. To date, although several transmissible neoplasia have been described in marine bivalves, transcriptome-wide functional studies are still lacking. However, Bruzos et al. (pre-print 2022, 32) have performed a comparative quantitative transcriptomic study between cancer cells and various healthy tissues in cockle BTNs, which supports their status as haemocyte-derived marine leukaemias.

Here, we performed a deep sequencing of cancer cell transcriptomes to investigate the gene expression profile of MtrBTN2. We sequenced mRNA from circulating cells in both healthy and MtrBTN2-affected *M. edulis*, and since that the founder host species was *M. trossulus*, we included haemocytes from this species as a control. Analysis of differentially expressed genes unveiled some functional characteristics of this unusual cancer and highlighted potential cancer drivers common to all bilaterians.

### 2. Results

MtrBTN2 cells were collected from the haemolymph of *M. edulis* mussels reared in the English Channel (Normandy, France). Only individuals with more than 95% of circulating neoplastic cells in their haemolymph were used for MtrBTN2 cell collection. *M. edulis* haemocytes were collected from individuals from the same geographical location but tested negative for MtrBTN2 by cytology and qPCRs targeting a sequence in the EF1 gene specific for the *M. trossulus* species and a MtrBTN2 specific sequence in the mitochondrial COI gene (16,24). *M. trossulus* haemocytes were collected from the haemolymph of wild *M. trossulus* found in the Barents Sea (Kola Bay, Russia), as this species is absent from the French coasts. All *M. trossulus* mussels were positive for the qPCR targeting the *M. trossulus* specific sequence, but negative by cytology and qPCR for MtrBTN2. We did not observe any other pathogen by histology in the nine individuals used for RNA-Seq analysis. Transcripts from haemolymph samples were sequenced for each condition: MtrBTN2-positive (i.e. cancerous) *M. edulis* (CAN 1-3), MtrBTN2-negative *M. edulis* (EDU 1-3) and MtrBTN2-negative *M. trossulus* (TRO 1-3). All RNA-seq reads were used to assemble *de novo* a pantranscriptome from the 9 samples (CAN 1-3, EDU 1-3, and TRO 1-3). The graphical representation of the experimental setup and bioinformatic analyses is shown in Figure 1.

**Figure 1:**
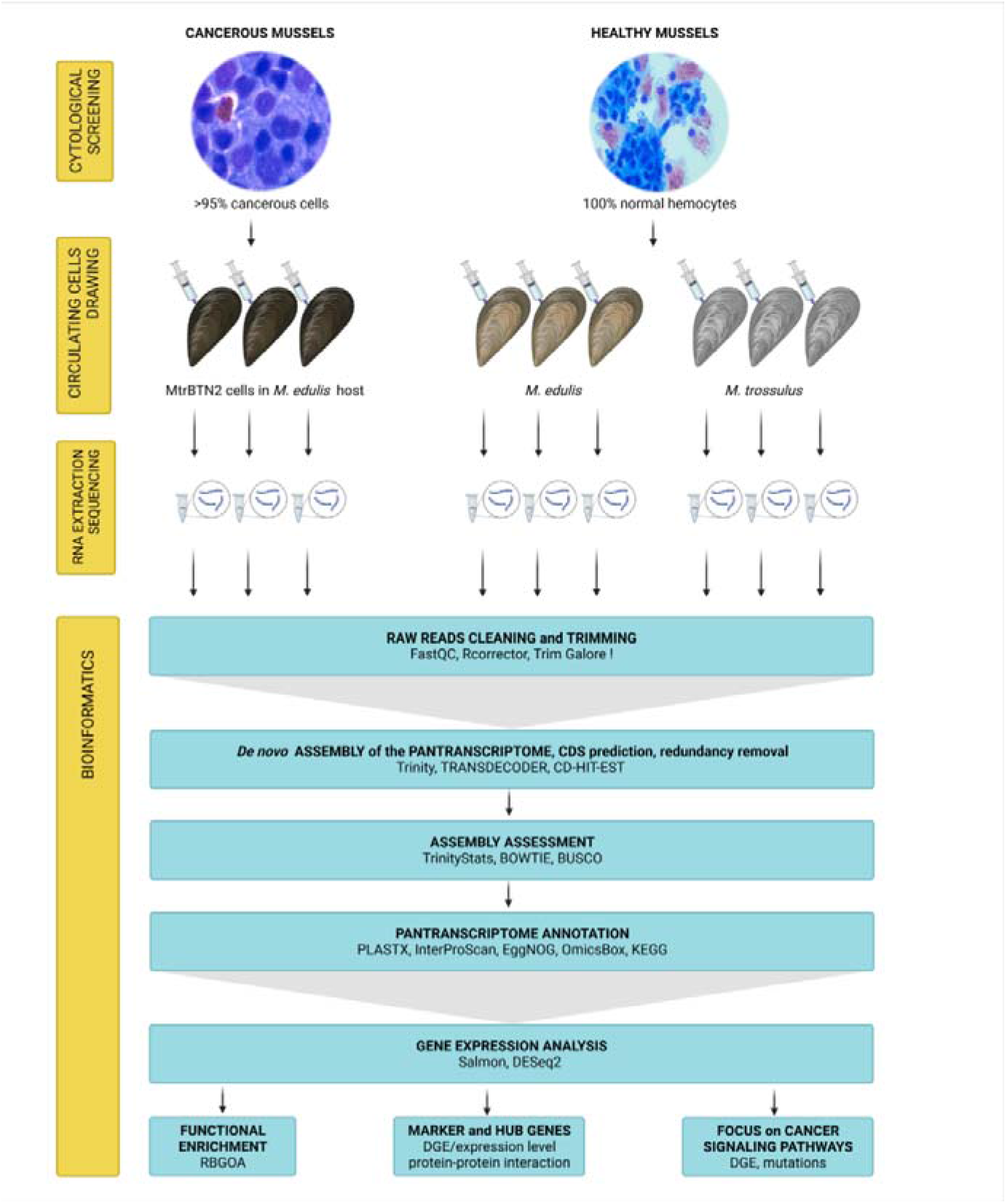
Graphical representation of the experimental design and RNA-Seq data analysis. DGE: differential gene expression analysis. Microphotographs of cancerous and healthy circulating cells are on the same scale.

### 2.1 Pantranscriptome assembly and annotation

The raw assembly contained 2,149,788 transcripts. After filtering, we retained a total of 138,071 transcripts consisting of predicted coding sequences (CDS). The pantranscriptome completeness assessment revealed that 95.6% of the highly conserved single-copy metazoan genes (n = 954) were present in full length as a single (81.2%) or duplicated (14.4%) copies. The mean remapping rate was 53.38 ± 0.58% with negligible differences between samples (Kruskal-Wallis, H(2)=1.69, p=0.43), which is acceptable for a remapping on CDS (33). Only 5826 transcripts showed no blast similarity or a conserved domain. The OmicsBox program v2.0 (34,35) assigned GO terms to 131,346 transcripts, but ultimately 41.39% (57,154) of transcript annotations were retained after computing a filtration step based on a minimum annotation score (Supplementary Table S1). The assembly and annotation metrics are shown in Table 1. Raw read sequences coming from RNA-seq were deposited in the NCBI BioProject database (https://www.ncbi.nlm.nih.gov/bioproject/) with links to BioProject accession number PRJNA749800. Individual SRA numbers are listed in Supplementary Table S2.

**Table 1:** Assembly and annotation metrics. The pantranscriptome was obtained from the mRNA sequencing of circulating cells from three Mytilus edulis, three M. trossulus, and three MtrBTN2-affected M. edulis at a late stage of the disease (>95% of cancer cells).*after applying the OmicsBox annotation rule (e-value hit filter of 1.10, annotation cutoff of 55, a HSP-Hit coverage of 60%).

| Attributes | Cancerous individuals |  |  | Healthy individuals |  |  |  |  |  |
| --- | --- | --- | --- | --- | --- | --- | --- | --- | --- |
|  | *MtrBTN2* |  |  | *Mytilus edulis* |  |  | *Mytilus trossulus* |  |  |
|  | CAN1 | CAN2 | CAN3 | EDU1 | EDU2 | EDU3 | TRO1 | TRO2 | TRO3 |
| Clean read pairs x10<sup>6</sup> | 83,69 | 82,83 | 82,05 | 83,88 | 81,93 | 98,33 | 81,57 | 76,67 | 84,82 |
| GC% | 36 | 36 | 36 | 37 | 37 | 37 | 36 | 36 | 36 |
| **Transcriptome statistics** |  |  |  |  |  |  |  |  |  |
|  | Raw number of transcripts |  |  |  |  |  | 2,149,788 |  |  |
|  | Final number of transcripts |  |  |  |  |  | 138,071 |  |  |
|  | GC% |  |  |  |  |  | 38 |  |  |
|  | Total assembled bases |  |  |  |  |  | 97,665,057 |  |  |
|  | Contigs N50 (bp) |  |  |  |  |  | 1,038 |  |  |
|  | Median contig length (bp) |  |  |  |  |  | 439 |  |  |
|  | Average contig length (bp) |  |  |  |  |  | 745 |  |  |
|  | BUSCO % (*Metazoa*, n=954) |  |  |  |  |  | 95,6 |  |  |
|  | Complete single-copy |  |  |  |  |  | 167 |  |  |
|  | Duplicated |  |  |  |  |  | 745 |  |  |
|  | Fragmented |  |  |  |  |  | 34 |  |  |
|  | Missing |  |  |  |  |  | 8 |  |  |
| **Back alignment on pantranscriptome %** | 53,53 | 53,92 | 53,48 | 53,88 | 53,83 | 52,83 | 53,70 | 52,18 | 53,14 |
| **Annotation statistics** |  |  |  |  |  |  |  |  |  |
|  | Transcripts with GO associated |  |  |  |  |  | 131,346 |  |  |
|  | Transcripts annotated* |  |  |  |  |  | 57,154 |  |  |
|  | No-blast, no Interproscan hits found |  |  |  |  |  | 5,826 |  |  |

### 2.2 The specific transcriptomic profile of MtrBTN2 cells

Principal component analysis (PCA) based on the normalised count matrix (RLE values) discriminated the three groups of samples (CAN, EDU, TRO) well (Figure 2). These first results indicate that the transcriptomic profile of MtrBTN2 differs from the transcriptomic profile of healthy *Mytilus* haemocytes, irrespective of their species or geographical origin.

**Figure 2:**
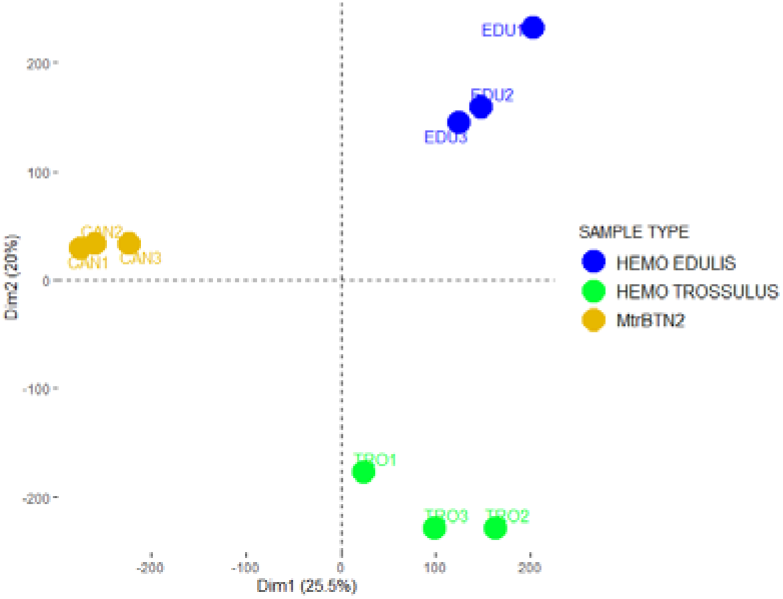
Principal component analysis (PCA) based on the normalised count matrix (RLE values). The first axis discriminates MtrBTN2 cells from hemocytes.

We investigated differentially expressed transcripts in MtrBTN2 cells (CAN 1-3) by comparing their abundance with *M. edulis* (EDU 1-3) and *M. trossulus* haemocytes (TRO 1-3) using DESeq2 v1.34 (36). These comparative results were then validated by RT-qPCRs (see sections 5.3 and 5.5) targeting 17 key-gene transcripts and confirmed in 10 additional haemolymph samples (CAN 7-11, EDU 7-11) (see Supplementary Table S3; Supplementary Figure S1 and S2). A total of 4,356/138,071 transcripts were significantly differentially expressed; 2,632 were more expressed and 1,724 were less expressed in MtrBTN2 cells than in haemocytes (see Log2 fold change (Log2FC) and adjusted p-values (FDR) in Supplementary Table S4). We further validated the significance of differential expression for 4,254 transcripts by running QuantEval (37), which builds connected components from the assembled transcripts based on sequence similarity and evaluates the quantification results for each connected component (Supplementary Tables S4 and S5, Supplementary Figure S3). Indeed, for 102 transcripts, the significance of the differential expression was not confirmed at the component level. A hierarchical clustering of DETs is shown on a log2 centred-heatmap (Figure 3). Although the results were less contrasted with RNA-seq data obtained from the whole flesh of other 6 additional individuals coming from a separate study (CAN 4-6, EDU 4-6) (see sections 5.3 and 5.4; Supplementary Table S3) than from haemolymph cells, the principal component analysis of the transcript abundances also confirmed a differential clustering between MtrBTN2-affected and MtrBTN2-free mussels (Supplementary Figure S4).

**Figure 3:**
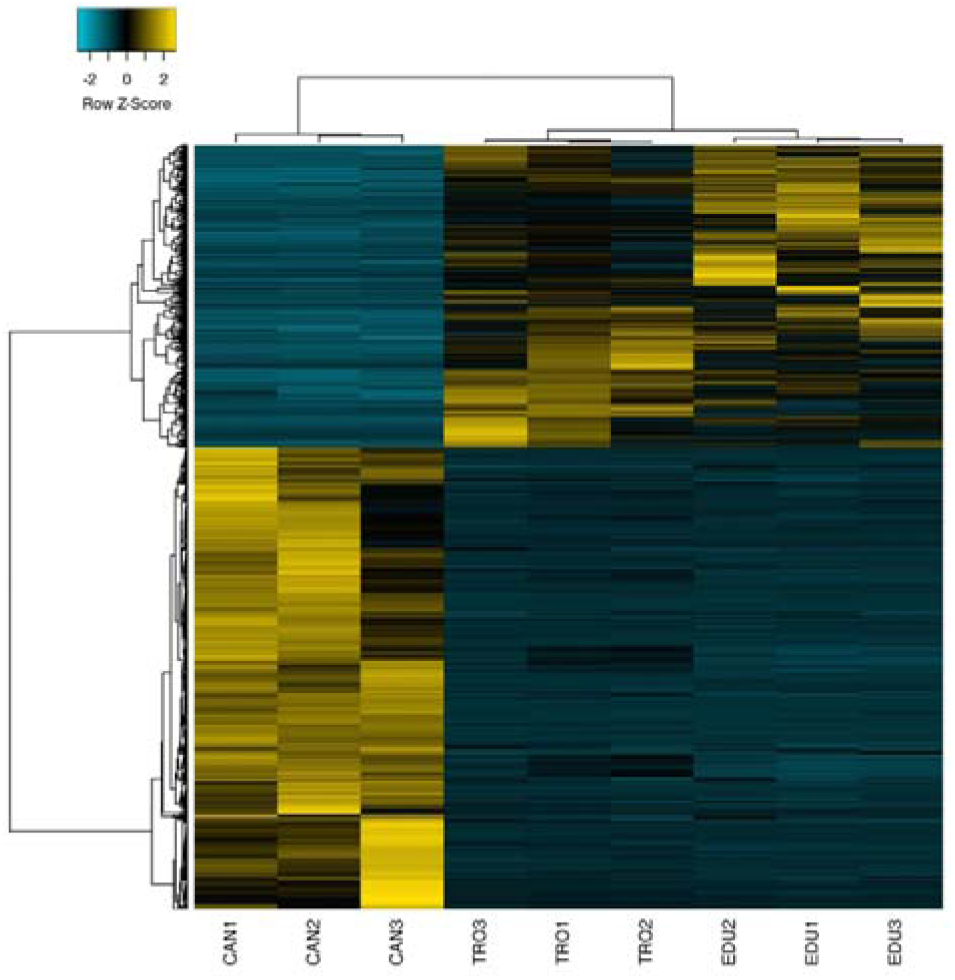
Heatmap (log2 centred) with hierarchical clustering of the 4254 differentially expressed transcripts at a FDR <0.05.

Among the most differentially expressed transcripts (DET) (with log2FC > 5), we identified six top-expressed (Relative Log Expression (RLE) > 500) transcripts of the genes *CYP11A1*, *THBS1*, *SELP*, *ALOX5*, *BMP2*, and *ADAMTS1* (Figure 4). These transcripts were also confirmed among the most differentially and top-expressed in the whole flesh of the additional 3 individuals (CAN 4-6) (see sections 5.3 and 5.5; Supplementary Table S3; Supplementary Figure S5). Indeed, these genes can be considered as specific for the malignant state and have been described to be involved in human tumorigenesis and metastasis (Table 2).

**Figure 4:**
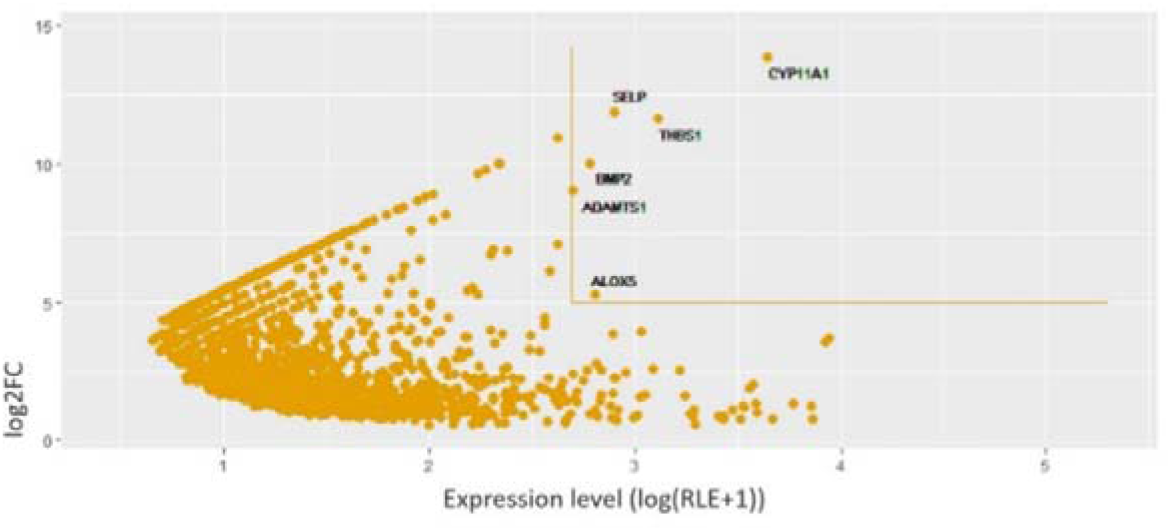
Scatter plot showing genes specific of the malignant state with y=log2FC, x=log(RLE+1). Lines delimit the areas above log2FC values > 5 and RLE > 500. Six transcripts of genes (CYP11A1, THBS1, SELP, ALOX5, BMP2, and ADAMTS1) exceeded these values and can be considered as specific of the malignant state.

**Table 2:**
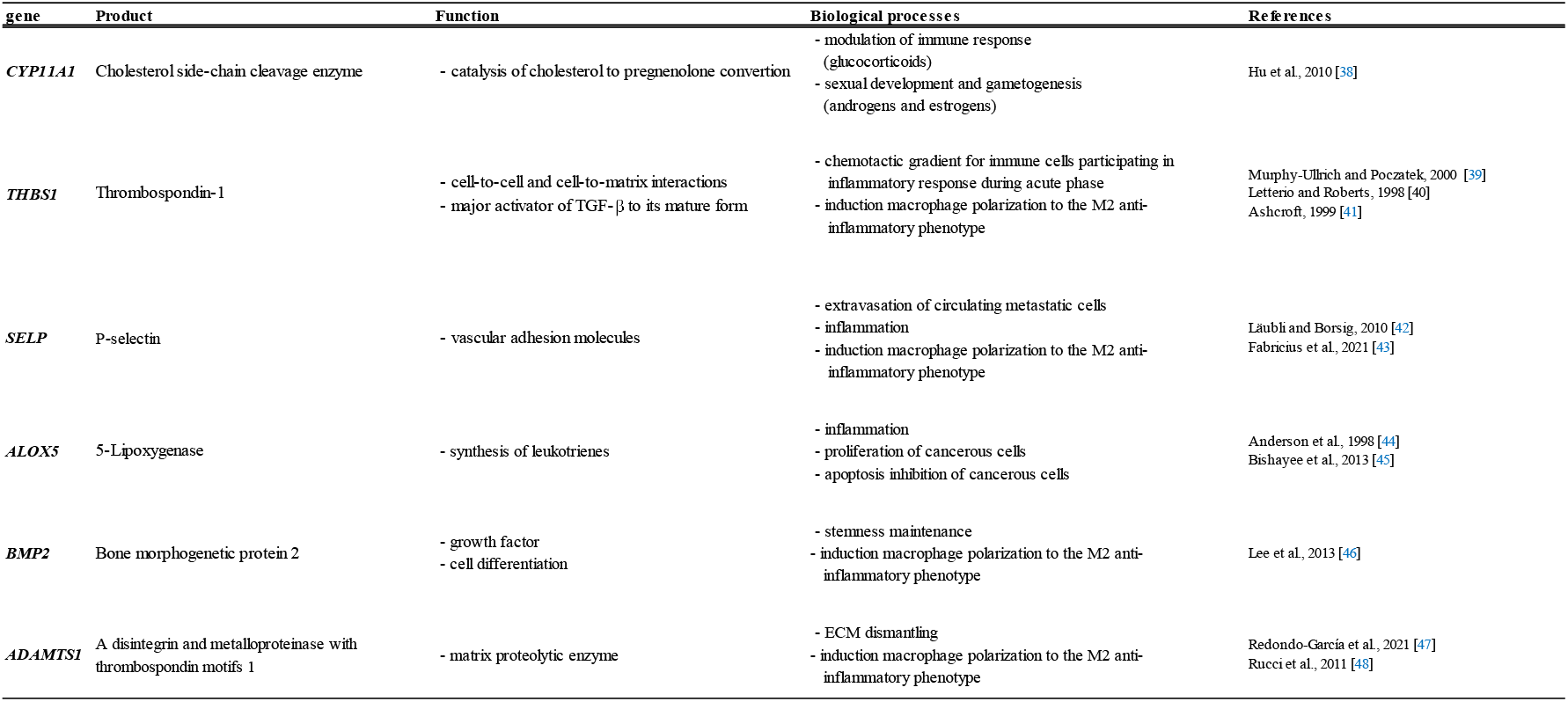
Top-expressed transcripts of genes in MtrBTN2 cells (log2FC > 5 and RLE > 500). Most are involved in inflammation and immune response.

They are associated with inflammation, immune processes, and extracellular matrix (ECM) disruption (38–48). The exact role of some of these genes (*THBS1, SELP, ALOX5,* and *BMP2*) during tumorigenesis is still under debate, mainly for their involvement in the inflammatory process (49,50). However, *THBS1* expression is increased in many cancers promoting invasion and metastasis (51–54). *SELP* and *ALOX5* are highly expressed in some tumours and cancer cell lines, inducing cancer cell proliferation (55–58) and inhibiting apoptosis (44,45). There are also conflicting data regarding the effect of *BMP2* on cancer (59). Nevertheless, most studies suggest that *BMP2* promotes metastatic progression and tumorigenesis. Although mollusc immunity against cancer has never been studied so far, most of these genes could be involved in modulating the host immune response. Some of them, such as *THBS1*, *SELP*, *BMP2*, and *ADAMTS1,* are involved in the induction of macrophage polarisation towards the anti-inflammatory M2 phenotype in mammals and their role in the interactions of MtrBTN2 with the host immune response deserves further attention to better understand the host invasion process.

### 2.3 Dysregulated biological processes in MtrBTN2

To identify the key biological processes altered in the MtrBTN2 phenotype, we performed a GO_MWU rank-based enrichment analysis on the DETs (for protein descriptions and gene IDs, see Supplementary Table S4). A biological process is considered to be altered if DETs are significantly overrepresented (enriched) within the total set of transcripts related to that process. The complete list of transcripts clustered by Gene Ontology (GO) term is provided in the Supplementary Table S6. The GO_MWU analysis revealed several biological functions that were differentially regulated in MtrBTN2 cells compared to normal haemocytes (Figure 5).

**Figure 5:**
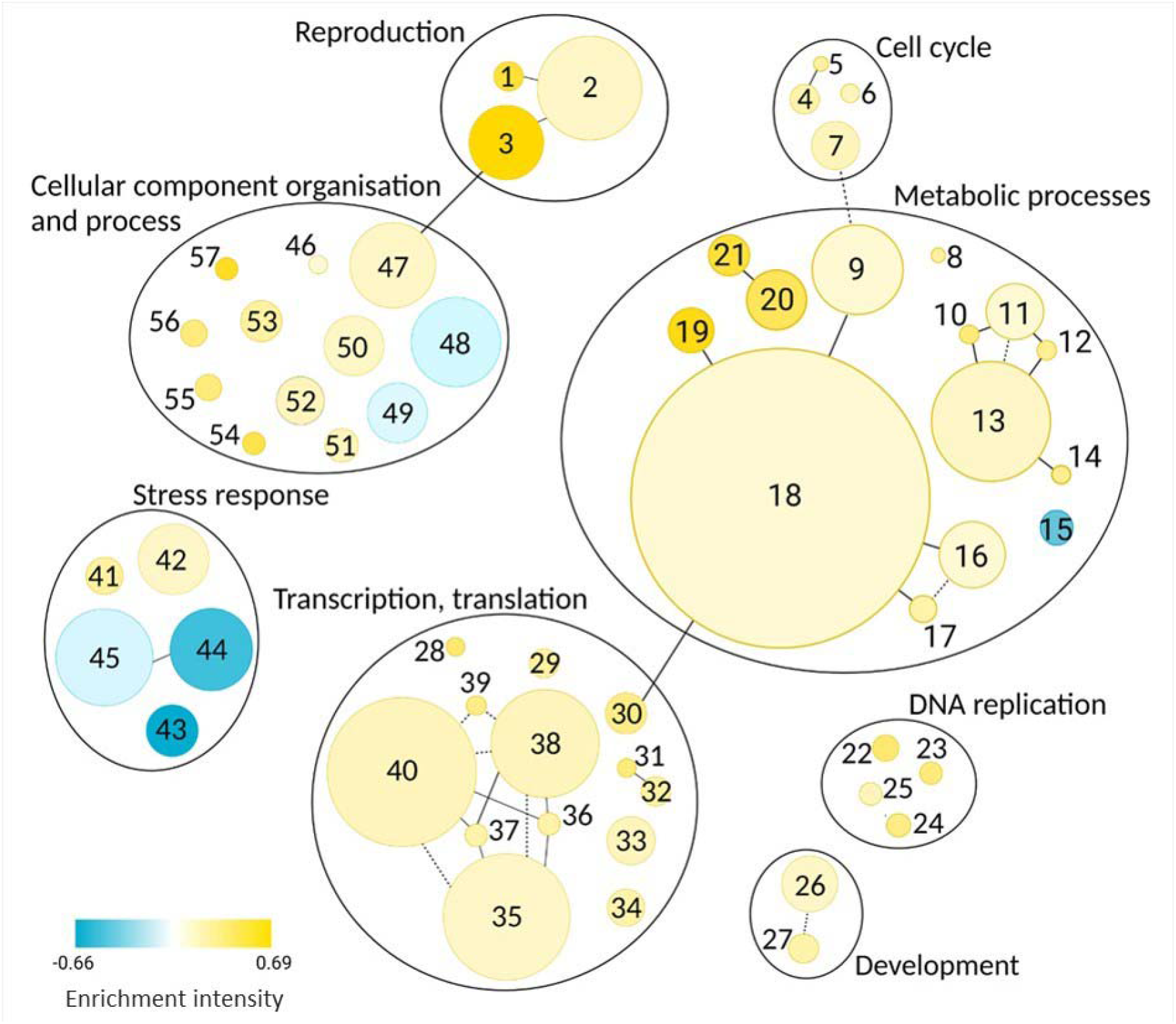
Biological processes dysregulated in transmissible neoplasia (p-value < 0.05): **1.**Meiosis I; **2.**Cellular process involved in reproduction; **3.**Sperm motility; **4.**Cytokenetic process; **5.**Contractil ring contraction; **6.**Mitotic spindle elongation; **7.**Regulation of cell cycle process; **8.**Protein deacylation; **9.**Proteolysis; **10.**Adenosine metabolic process; **11.**Nucleoside phosphate biosynthetic process; **12.**IMP metabolic process; **13.**Carbohydrate derivative metabolic process; **14.**Dolichol-linked oligosaccharide biosynthetic process; **15.**Tricarboxylic acid cycle; **16.**Peptidyl-amino acid modification; **17.**Histone modification; **18.**Protein metabolic process; **19.**Protein deubiquitination; **20.**Nucleobase metabolic process; **21.**Pyrimidine nucleobase metabolic process; **22.**DNA replication initiation; **23.**DNA strand elongation; **24.**DNA replication, synthesis of RNA primer; **25.**DNA-dependent DNA replication; **26.**Regulation of growth; **27.**Cell fate determination; **28.**Cis assembly of pre-catalytic spliceosome; **29.**RNA phosphodiester bond hydrolysis; **30.**Mitochondrial translation; **31.**Transcription initiation from RNA polymerase I promoter; **32.**DNA-templated transcription, initiation; **33.**Translational elongation; **34.**Spliceosomal snRNP assembly; **35.**mRNA processing; **36.**tRNA splicing, via endonucleolytic cleavage and ligation; **37.**tRNA wobble base modification; **38.**tRNA metabolic process; **39.**Deadenylation-dependent decapping of nuclear-transcribed mRNA; **40.**ncRNA metabolic process; **41.**Response to oxidative stress; **42.**Double-strand break repair; **43.**Cellular response to amino acid starvation; **44.**Response to bacterium; **45.**Defense response; **46.**Actin filament based process; **47.**Microtubule-based movement; **48.**Cell surface receptor signalling pathway; **49.**Anion transport; **50.**Golgi vesicle transport; **51.**Ribosomal small subunit assembly; **52.**Ribosomal large subunit assembly; **53.**rRNA-contain RNP complex; **54.**Protein secretion; **55.**Protein targeting to ER; **56.**Mitochondrial respiratory chain complex assembly; **57.**Protein insertion into ER membrane. The color intensity is proportional to the enrichment intensity, expressed as the ratio between the number of transcripts that were significantly over-(yellow) or down-expressed (blue) compared with the total number of transcripts in the process. The disk area is proportional to the number of DETs linked to the process (from 2 to 191 transcripts). Lines connecting two functions represent shared transcripts (solid line means that all transcripts in the smaller family are part of the largest family; dotted line means that only some transcripts are shared).

#### Proliferation-related functions are altered

Some functions consistent with a high proliferative activity are altered in MtrBTN2 cells. The “cell cycle” family (4.Cytokenetic process; 5.Contractil ring contraction; 6.Mitotic spindle elongation; 7.Regulation of cell cycle process, in Figure 5) is enriched in MtrBTN2 cells, as well as “DNA replication” (22.DNA replication initiation; 23.DNA strand elongation; 24.DNA replication, synthesis of RNA primer; 25.DNA-dependent DNA replication, in Figure 5). More specifically, negative regulation of the cell cycle progression appeared to be reduced: *ASAH2,* which inhibits the ceramide mediation of the cell cycle arrest (60), was more highly expressed, whereas *PPP2R5B,* which inhibits cell cycle progression (61), was less expressed in MtrBTN2. Moreover, transcripts of genes promoting the cell cycle progression were more abundant in MtrBTN2 cells, in particular five genes that form the Anaphase Promoting Complex (*ANAPC1, ANAPC4, ANAPC5, CDC26, and CDC23*) and *CCND1* encoding cyclin D1, which promotes progression through the G1-S phase of the cell cycle (62). *TERT,* which is essential for the chromosome telomere replication in most eukaryotes and is particularly active in progenitor and cancer cells (63) was also more expressed in MtrBTN2 cells. By maintaining telomere length, telomerase removes a major barrier to cellular longevity, allowing unlimited proliferation driven by oncogenes (64). In addition, in the “Development” family, we found a higher abundance of transcripts of genes involved in promoting cell growth, such as *TTK* (65), *METAP2* (66), *THBS1* (activator of TGF-β) (39), *MEAK7* (67) and *LAMTOR1* (68). Transcripts of several genes that act as growth inhibitors were less abundant in MtrBTN2, as often seen in cancer contexts, such as *FAM107A* (69) and *LTBP3* (70), which keeps TGF-β in a latent state.

#### Metabolic pathways are modified

Many metabolic processes were shown to be altered (Figure 5 and Supplementary Table S6). Nucleotide metabolism in MtrBTN2 cells was characterised by a higher expression of more than 25 transcripts of genes involved in the *de novo* and salvage biosynthetic pathways of purines and pyrimidines and in the homeostasis of cellular nucleotides. The metabolism of protein and carbohydrate derivatives was also modified. The enrichment concerned transcripts of genes encoding ribosomal proteins, protein turnover and localisation (especially deubiquitination), proteolysis, peptidyl amino-acid modification, histone acetylation, and lipid and protein glycosylation. In addition to *MEAK7* and *LAMTOR1*, transcripts of eight genes (*BMP2*, *KICS2*, *ITFG2*, *MIOS*, *NPRL2*, *RagA, RagD, and SEC13*) involved in the mTORC1 amino-acid sensing pathway (43.Cellular response to amino acid starvation, in Figure 5) (71) were differentially expressed. Interestingly, in addition to the role of mTORC1 in the regulation of autophagy, mTORC1 plays a pivotal role as a master regulator of protein, lipid, and nucleic acid metabolism changes that have been reported in many different cancer cells (72–75). Several transcripts of genes involved in organelle activities and related to protein metabolism (anabolism, catabolism, post-translational modifications, folding, and secretion) were also more highly expressed in MtrBTN2.

The tricarboxylic acid cycle (TCA) also appeared to be differentially regulated in MtrBTN2, with a lower expression of *DLST* (a component of the 2-oxoglutarate dehydrogenase complex), *SDHD* and *NNT*, and a higher expression of *SUCLA2*. The TCA cycle is critical for the generation of cellular energy and precursors for biosynthetic pathways (76). By screening the expression of 16 glycolysis-related genes, we found that *PFK*, *PKM*, *SLC2A1, and SLC2A3* were more expressed; however, *PGK1*, *TPI1*, and *ALDOA* were less expressed than in control haemocytes. These conflicting results did not allow us to conclude on how the regulation of glycolytic activities is altered in transmissible neoplasia cells. However, these results suggest major changes in nucleotide, amino-acid and energy metabolism in MtrBTN cells as often seen in other cancers (77).

#### DNA repair systems are dysregulated

MtrBTN2 cancer was found to express at higher levels a significant panel of transcripts of genes involved in the DNA Homologous Recombination (HR) (42.Double-strand break repair, in Figure 5), such as *BABAM1*, *ZSWIM7*, *RAD51L*, *RAD54L*, *MRE11*, *MCM9*, and *TONSL*. The DNA helicase MCM9 is also involved in the repair of inter-strand crosslinks (78), as is MUS81 (79), which is also more expressed in MtrBTN2 cells. This type of damage is a double-edged sword, as double-strand breaks can induce cell death if not efficiently repaired, but inefficient or inappropriate repair can lead to mutation, gene translocation and cancer (80). Due to their high proliferation rate and increased metabolic activities, cancer cells are under a huge replication stress (81), which may emphasise the need for these long-lived cancer lineages to activate DNA repair to persist on time. Cells are able to repair double-strand breaks through two major pathways: non-homologous end joining and HR (82). HR is critical for re-establishing replication forks at the sites of damage during S and G2 phases of the cell cycle. However, when combined with the depletion of cell cycle checkpoints and apoptosis, this mechanism can lead to genomic instability. As the ploidy of MtrBTN2 appears to be highly abnormal (varying from 8N to 18N, (19)), further investigations will be required to determine whether the dysregulation of these DNA repair pathways favours recombination between homologous chromosomes and the accumulation of chromosomal abnormalities, or whether these pathways are activated in order to limit DNA damage. Interestingly, upregulation of double-stranded DNA break repair via the HR pathway has also been found in DFTD, in addition to repair *via* non-homologous end joining (83).

#### Interaction with ECM is affected

The expression levels of transcripts of seven genes encoding integrin subunits (48.Cell surface receptor signalling pathway, in Figure 5), a major class of transmembrane glycoproteins that mediate cell-matrix and cell-cell adhesion, were altered (Supplementary Table S6). As integrins are mostly less expressed in MtrBTN2 cells, this suggests that the ability of these cells to interact with ECM components is profoundly altered, which is often involved at multiple stages of the cancer processes (84).

#### Cell fate determination pathways are modified and soma-to-germline transition may have occurred

Major cell fate pathways appear to be modified (Figure 5 and Supplementary Table S6). The expression of transcripts of six genes involved in the Notch and Wnt pathways was altered. These changes may drive dedifferentiation processes often observed in aggressive cancer cells or cancer stem cells (85). We report the results of the focused analysis performed on these two signalling pathways in section 2.5. In addition, three biological processes clustered under GO terms related to “reproduction” were enriched (1.Meiosis I; 2.Cellular process involved in reproduction; 3.Sperm motility, in Figure 5). Nineteen transcripts more expressed in MtrBTN2 cells gathered under the “sperm motility” process and encode dynein assembly factors, dynein chains, and cilia and flagella associated proteins. Moreover, transcripts of genes involved in meiosis such as *MSH5* (86)*, SYCP3* (87) and *TEX11* (88), were more abundant in MtrBTN2. These unexpected results may reflect a soma-to-germline transition during the oncogenic process, which has been reported in a variety of cancers, and support the hypothesis of profound changes in the differentiation state of MtrBTN2 cells (89,90). Interestingly, when abnormally produced in mitotically dividing cells, the *SYCP3* can impair recombination and drive ploidy changes that affect chromosome segregation in cancer cells, which may be one of the mechanisms that led to the hyperploidy of MtrBTN2 cells (90).

#### Immune response is altered

One of the most striking differences between MtrBTN2 cells and haemocytes is that biological processes belonging to innate immunity (44.Response to bacterium; 45.Defense response; in Figure 5) are significantly underrepresented in MtrBTN2 cells. Transcripts of genes encoding important mussel antimicrobial peptides and bactericidal proteins such as *Mytilin B*, *Myticin A*, *Myticin B*, *MGD1*, and *BPI* (91–93) were significantly less abundant in MtrBTN2 cells. A similar pattern was also observed for genes encoding several pattern recognition receptors, such as *TLR2* and *TLR4,* which belong to the Toll-Like-Receptor family and are essential for innate immunity against pathogens (94), and the lectin-related pattern recognition molecules *FCN2*, *FIBCD1*, *CLEC4E*, and *CLEC4M*, which are involved in the recognition of pathogens and apoptotic or necrotic cells (95–97). This underrepresentation of biological functions related to host defences suggests that MtrBTN2 cells are immunologically incompetent. During MtrBTN2 disease progression, we also observe a drastic decrease in the number of host haemocytes, which are progressively replaced by circulating MtrBTN2 cells, which can represent over 95% of the circulating cells in cancerous mussels (19,22). It remains to be determined whether MtrBTN2 cells outcompete normal haemocytes in the haemolymph or whether they interfere with haematopoiesis, as seen in some human leukaemias (98). Still, the combination of fewer circulating haemocytes and the immune incompetence of MtrBTN2 cells could have dramatic consequences for the host health, leading to lethal opportunistic systemic infections (99).

### 2.4 CASP3, FN1, and CDC42 are hub genes in the interaction network of MtrBTN2

Protein-protein interaction (PPI) network of the DETs was constructed to evaluate their connectivity and to identify hub genes that may play a critical role in the MtrBTN2 phenotype. The complete PPI network is shown in Figure 6. The top 20 genes in the connectivity ranking were found in functions already identified as enriched with GO_MWU, such as translation (most involved in ribosome biogenesis) and DNA replication. Outputs from NetworkAnalyzer v4.4.8 (100) are reported in Supplementary Table S7. Interestingly, this analysis highlighted three genes as major hubs in the PPI network that did not stand out in the GO_MWU analysis, namely *FN1, CDC42,* and *CASP3*. Fibronectin 1, encoded by *FN1*, is a major component of the ECM and plays an important role in cell adhesion, migration, growth and differentiation (101). The abundance of *FN1* transcripts was significantly higher in MtrBTN2 cells (log2FC > 5). CDC42, a member of the Rho GTPase family, plays an important role in cell-cell and cell-matrix adhesion, actin cytoskeleon architecture, cell migration, and cell cycle regulation (102); corresponding transcripts were significantly less abundant in MtrBTN2 cells. As Fibronectin 1 and CDC42 are involved in complementary functions and signalling pathways (such as cell-ECM interactions and cell migration), both hub genes were found to be interconnected in the PPI network. In addition, a large number of integrin subunits were found to be dysregulated. Taken together, these results strongly suggest that cell-ECM interactions and cell migration functions are profoundly modified in MtrBTN2 cells, consistent with their observed rounded morphology, poor adhesion properties and f-actin modifications (19,21). The master effector of apoptosis, Caspase 3, encoded by *CASP3*, is also less expressed in MtrBTN2 cells. It plays a central role in the execution phase of cell apoptosis (103), which is often inhibited in cancer cells, allowing them to escape cell death, despite the accumulation of DNA damage and cell cycle dysfunction.

**Figure 6:**
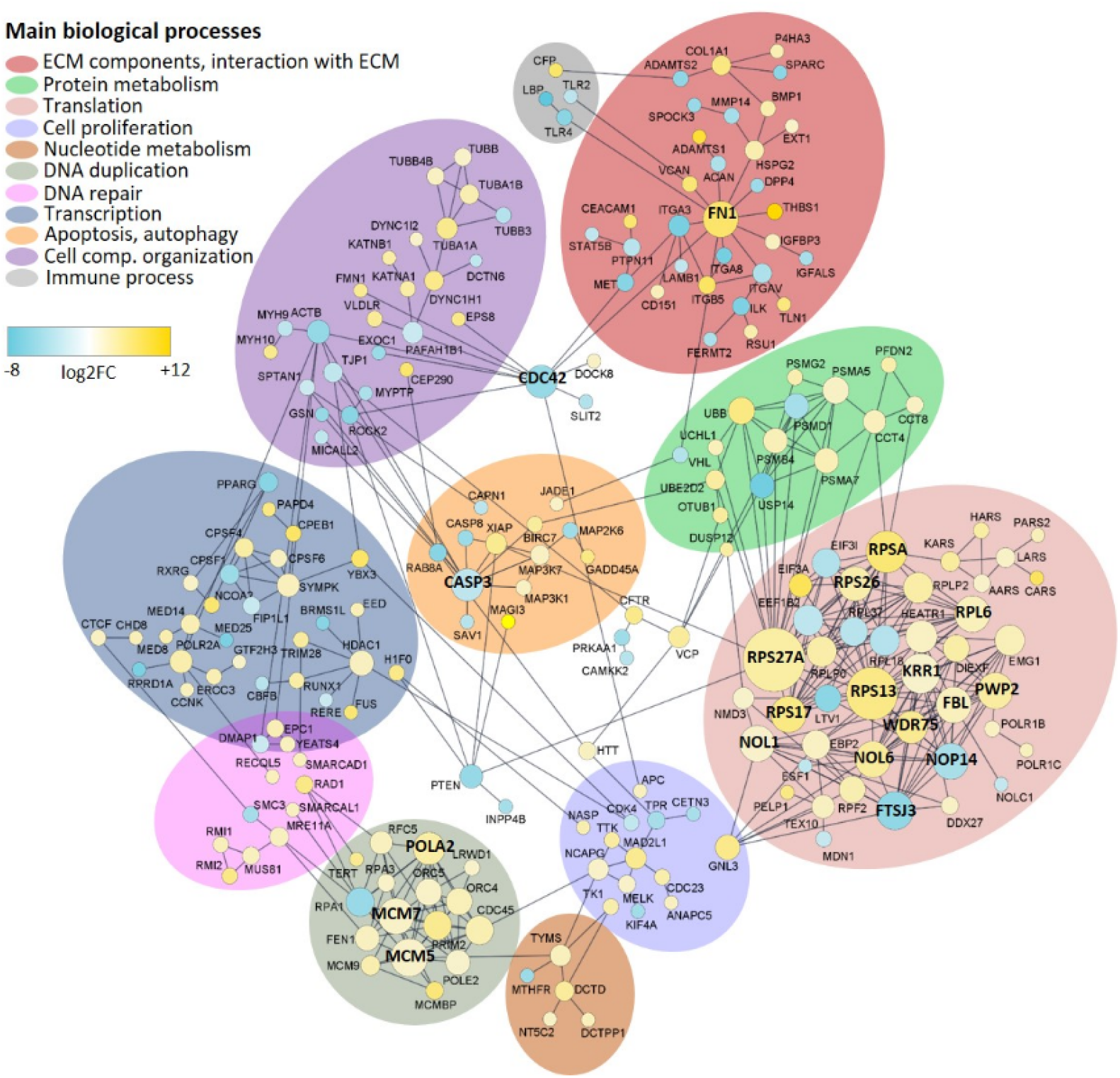
Protein-protein interactions (PPI) network. The size of the disc is proportional to the number of connections. The top 20 Hub proteins are shown in bold. Up-regulated genes are shown in yellow; Down-regulated genes are shown in blue. The color intensity is proportional to the Log2FC value.

### 2.5 Most key oncogenic signalling pathways are altered in MtrBTN2

Based on the Cancer Genome Atlas project, Sanchez-Vega et al. (26) highlighted that 10 signalling pathways are very frequently altered in most human cancers. We found that six of these 10 oncogenic signalling pathways may be altered in MtrBTN2 cancer. These pathways are involved in cell proliferation and differentiation.

DETs were found for key genes related to the “Hippo”, “Notch”, “Wnt”, “Myc”, “PI3K”, and “Cell Cycle” signalling pathways (Table 3).

**Table 3:** Transcripts of genes differentially expressed in MtrBTN2 among the most frequently altered genes in the human oncogenic signalling pathways. We report their action on the corresponding pathway, their expression status, and the Log2(FC). Two genes encoding JAG1 (confirmed as distinct by QuantEval analysis) were more expressed in MtrBTN2 cells.

| Pathway | Main functions | Gene | Action on pathway | MtrBTN2/hemocytes expression | Log2FC |
| --- | --- | --- | --- | --- | --- |
| Hippo | anti-proliferative<br>anti-stem cell self renewal<br>pro-apoptotic | SAV1 | activation | Lower | -3,65 |
| Notch | cell proliferation<br>cell fate | JAG1 (2 genes) | activation | Higher | 3.75 and 4.32 |
|  |  | FBXW7 | repression | Higher | 1,24 |
| Wnt | cell proliferation<br>stemness maintenance | SFRP1 | repression | Lower | -3,54 |
|  |  | SFRP3 | repression | Higher | 7,38 |
|  |  | APC | repression | Higher | 2,63 |
|  |  | CTNNB1 | / | Higher | 1,09 |
| Myc | cell proliferation<br>apoptosis | MAD3 | repression | Lower | -2,19 |
| PI3K | cell proliferation<br>cell survival | PTEN | repression | Lower | -4,51 |
|  |  | PIK3R1 | repression | Lower | -1,68 |
| Cell Cycle |  | CCND1 | activation | Higher | 1,89 |

The Hippo signalling pathway plays a role in inhibiting cell proliferation, controlling cell fate, and promoting apoptosis through the phosphorylation of YAP/TAZ transcription coactivators, their cytoplasmic retention and degradation. *SAV1* is a key activator of the Hippo pathway involved in LATS1/2 phosphorylation, a necessary step for subsequent YAP/TAZ phosphorylation (104). As *SAV1* is less expressed in MtrBTN2 cells, the Hippo pathway may be downregulated or inactivated.

The PI3K pathway also showed dysregulation. It is involved in cell metabolism, growth, proliferation, cell-ECM interactions and survival. We observed a lower expression of two genes, *PTEN* and *PIK3R1*, which act as PIK3 antagonists (105). When the pathway is activated, AKT1 is phosphorylated and inhibits the activity of TSC1/2. AKT-mediated phosphorylation of TSC1/2 removes its inhibition of RHEB activity, leading to activation of the mTORC1 complex. Our enrichment analysis (section 2.3) has also highlighted the activation of mTORC1 by the alternative amino-acid sensing pathway. In addition, AKT1 plays an indirect role in the Wnt and Myc pathways by negatively regulating GSK3β through its phosphorylation (106). Thus, the inhibitory effect of unphosphorylated GSK3β on MYC and its contribution in the destruction complex of β-cathenin in the Wnt pathway may be repressed in MtrBTN2.

The gene encoding MAD3, an antagonist of MYC for MAX binding (107), was less expressed in MtrBTN2 cells, suggesting that the MYC-MAX complex is favoured and the pathway is activated.

Among SFRPs, which act as Wnt pathway inhibitors by binding extracellular WNT ligands, *SFRP1* was less expressed and *SRFP3* more. Moreover, *APC,* which encodes a constitutive protein of the Destruction Complex, was more expressed. APC is a multifunctional protein and is involved, for instance, in the normal compaction of mitotic chromatin (108). We looked for differentially expressed genes among the downstream genes of the Wnt pathway, which transcription is regulated by unphosphorylated CTNNB1 mediation. We found that *CCND1*, *JAG1*, and *FN1* were more expressed suggesting that the Wnt pathway was activated (109). CCND1 promotes cell proliferation and JAG1 is a transmembrane ligand that activates the Notch pathway. However, FBXW7, which inhibits the NOTCH cleavage and its subsequent activation, was more expressed in MtrBTN2 cells. In addition, the *HES1* expression, which is promoted by activated NOTCH, was lower in MtrBTN2 cells. HES1 negatively regulates the expression of downstream target genes such as tissue-specific transcription factors (110).

To identify potentially relevant mutations related to these pathways, we screened for the presence of MtrBTN2-specific SNVs in the transcript sequences of key genes. Although the recovery of complete transcript haplotypes was hampered by the polyploidy of cancer cells and the short-read sequencing data, we were able to identify MtrBTN2-specific SNVs, variants that were only observed in the three MtrBTN2 samples (CAN 1, 2, 3) but not in the 6 MtrBTN2-free samples (TRO1-3, EDU1-3). MtrBTN2-specific SNVs were identified in 8 key genes (Table 4; for alignments see Supplementary Files S1 and S2) and were confirmed to be present in the additional cancerous individuals (CAN 4-6).

**Table 4:** Genes with transcript sequences carrying MtrBTN2-specific SNVs among the most frequently altered genes in the oncogenic signalling pathways. AKT and MYC carried missense substitutions in a protein domain.

| Gene | Transcript length | Pathway | Substitution type |  | Total number | Missense substitution<br>in a domain |
| --- | --- | --- | --- | --- | --- | --- |
|  |  |  | Synonymous | Missense |  |  |
| *PIK3CA* | 3144 bp | PI3K | 33 | 0 | 33 | - |
| *PIK3CB* | 3159 bp | PI3K | 3 | 0 | 3 | - |
| *PTEN* | 1395 bp | PI3K | 2 | 0 | 2 | - |
| *AKT* | 1458 bp | PI3K | 1 | 1 | 2 | 1 |
| *p53* | 1797 bp | p53 | 4 | 1 | 5 | 0 |
| *MDM2* | 1647 bp | p53 | 21 | 10 | 31 | 0 |
| *MYC* | 1269 bp | Myc | 0 | 2 | 2 | 2 |
| *FBXW7* | 2100 bp | Notch | 1 | - | 1 | - |

The highest number of MtrBTN2-specific SNVs were found in *PIK3CA* (PI3K pathway), with 33 MtrBTN2-specific SNVs/3144 nucleotides. All these specific SNVs were synonymous substitutions but missense substitutions were present in four genes: *AKT*, *p53*, *MDM2*, and *MYC*. In two of these genes, the missense substitutions were located within protein domains. The missense substitution in the *AKT* sequence was located in the kinase domain, and the two missense substitutions in the *MYC* sequence in the transcriptional activation domain. Interestingly, in the *p53* gene which encodes both p53 and p63/73 proteins in molluscs (111) we found five MtrBTN2-specific SNVs. The variants T177C and C816T are synonymous substitutions previously described by Vassilenko et al. (2010) (112) in *M. trossulus* affected by haemic neoplasia. The variant A1264T is a missense substitution, newly described here, but is not located in a domain. *M. trossulus p53* is characterized by a deletion of 6 nucleotides if compared to *M. edulis p53* (113). We found this deletion in 100% of reads coming from MtrBTN2 samples. The frequencies of these MtrBTN2-specific SNVs are reported in Supplementary Table S8, and in most cases, heterozygosity was observed, with maintenance of expression of the *M. trossulus* alleles whereas host-specific (*M. edulis*) alleles were predominantly absent (Supplementary Files S1). Indeed, the real impact of these variants on the cellular functions is difficult to assess and this MtrBTN2-specific variant set could include both somatic variants and ancestral germline polymorphisms.

## 3. Discussion

In their survey of bilaterian animals, Aktipis et al. (114) have revealed that all or almost all bilaterians are susceptible to cancer. Molluscs (Lophotrochozoa) are no exception (20,22). Although a high degree of genotypic and phenotypic diversity has been described among mammalian cancers (115), molecular studies, mainly in humans and mice, have identified common drivers of the cancer process (25,26). Still, oncogenic processes in invertebrate phyla have been poorly studied, especially at the molecular level. Even if expression levels are not the only factor determining the effect of genes on phenotypes, our present transcriptomic data suggest that the most common alterations associated with oncogenesis found in mammals are present in MtrBTN2 cells. Six of the major oncogenic signalling pathways are likely to be altered. One of the most fundamental traits of cancer cells is their high proliferation rate. We have previously found a mean doubling time of ∼ 3 days for MtrBTN2 (24). The present study provides information on the molecular bases of the high proliferation of MtrBTN2. The overexpression of these proliferation-related pathways is probably even more evident because we have used specialised cells (haemocytes) as a reference. At present, the issue of haematopoiesis in bivalves is poorly understood; however, some circulating haemocytes are capable of dividing and expressing genes involved in cell proliferation, particularly during the immune response (116). The proliferative activity of MtrBTN2 appears to be supported by metabolic rewiring, as is often seen in cancer cells (77). Indeed, we highlight a possible central role for the Pi3K-AKT-mTORC1 pathway, which is a master regulator in coupling cell growth and amino-acid, lipid, and nucleotide metabolisms that are frequently modified in many mammalian cancers of diverse origin (117–119). Our data also suggest inhibition of apoptosis. Dysregulation of receptors and pathways involved in cell-ECM interactions (120) was also highlighted here, which is consistent with our knowledge of MtrBTN2 biology, pathogenesis (intra-host disease progression) and its inter-host transmission. Indeed, MtrBTN2 cells are non-adherent circulating cells that infiltrate tissues and breach physiological barriers such as epithelium during the transmission process. MtrBTN2 cells are also characterised by a dysregulated differentiation state often seen in aggressive cancers (85), as reflected by the aberrant activation of some meiosis-related genes suggesting a soma-to-germline transition. Such reactivation of meiosis-related gene expression has previously been observed in a variety of non-germ cell cancers in humans (89). Finally, our transcriptomic data suggest a dampening of host defenses in cancerous mussels, which is likely to facilitate disease progression and pathological outcome. Indeed, evasion from immune destruction by silencing components of the immune system is another hallmark of human cancers (121).

This transcriptomic analysis generated a large amount of information about gene expression in MtrBTN2 cells and we identified candidate genes and pathways associated with their cancerous state. These results represent a significant step forward in understanding this disease at the cellular and molecular level, and lay the groundwork for future research. To date, most of the molecular dysfunctions have not been studied in molluscs and may differ to some extent from vertebrates. Priorities are to i) obtain well curated and annotated reference genomes, ii) accurately characterise gene products, and iii) specifically describe protein functions and relationships within and between pathways. To achieve these goals, gene manipulation tools are strongly required, which is a major challenge in such non-model organisms. However, a comparative oncology approach offers a unique and powerful opportunity to learn more about the evolutionary mechanisms of cancers and metastatic processes. Our model has several advantages. It is a naturally occurring cancer, it is transmissible, ensuring the repeatability of analyses and long-term studies, and molluscs are easy to handle under laboratory conditions with few ethical concerns.

## 5. Materials and Methods

### 5.1 Mussel samplings and MtrBTN2 diagnostic

We collected mussels in the English Channel and the Barents Sea in late 2019. Two hundred *Mytilus edulis* were sampled in a farm located in Agon-Coutainville (49°0’44.784’’N 1°35’55.643’’O, Normandy, France) and immediately screened for cancer at the LABEO laboratory (Caen, France). The presence of circulating cancer cells and the stage of the disease were initially diagnosed by cytological examination of haemolymph samples (19). We found 14 positive individuals among which three were at an advanced stage of the disease (>95% of circulating cells were cancer cells). After anaesthesia (122), we drew a maximum volume of haemolymph from the adductor muscle of these three cancerous mussels, as well as from three mussels diagnosed as MtrBTN2-free. The haemolymph was placed individually in RNase-free microtubes kept on ice and centrifuged at 800 x g for 10 min at 4°C. The pelleted cells were immediately resuspended in TRIzol® (invitrogen) and stored at −80°C until RNA extraction. As non-transmissible circulating cancers also exist in mussels (20), we confirmed the MtrBTN2 diagnosis by two qPCRs, one specific for *M. trossulus* (and MtrBTN) and targeting the nuclear marker EF1α (24) and one specific for the MtrBTN2 lineage targeting mtCR (16). Briefly, a piece of mantle and gills was fixed in 96% ethanol and used for DNA extraction using the Nucleomag® 96 Tissue kit (MachereyNagel). We carried out both qPCRs on cancerous and cancer-free *M. edulis* using the sensiFAST^TM^SyBR® No-ROX Kit (Bioline) and the LightCycler® 480 Real-Time PCR (Roche Life Science) system. A transversal section (including gills, digestive gland, mantle, and gonad) of each individual was fixed in Davidson for 48h and embedded it paraffin (RHEM facility). Sections were cut at 3 µm-thin and stained with haematoxylin and eosin to confirm the health status of each mussel and to exclude the presence of other pathologies.

Three *M. trossulus* were collected in the wild in Mishukovo (69°02’39;34.3’’N 33°01’39;45.9’’E, Kola Bay, Russia, Barents Sea). The haemolymph was drawn as for *M. edulis* individuals, the cells were pelleted and resuspended in 100 µL of RNA*later^TM^* (invitrogen) and sent to our laboratory where they were frozen at −80°C until RNA extraction. We also received a piece of mantle and gills from each individual fixed in 96% ethanol for genetic screening and to exclude the presence of MtrBTN, and tissue sections in 70% ethanol after 48h of fixation in Davidson’s solution. *M. trossulus* individuals were subjected to both qPCRs and histological examination.

### 5.2 Total RNA extraction, library preparation, and sequencing

*M. trossulus* samples preserved in RNA*later* were centrifuged at 5000 x g for 10 min at 4°C, the supernatant removed, and the cells were resuspended in 500 µL of TRIzol®. *M. edulis* preserved directly in TRIzol® were thawed. Both sample types were incubated at room temperature for 20 min under agitation to lyse the cells. RNA extraction was performed using Direct-zol^TM^ RNA MiniPrep according to the manufacturer’s instructions (Zymo Research). Quantification and integrity of total RNA were checked using a NanoDrop® ND-1000 spectrophotometer (Thermo Scientific) and by capillary electrophoresis on a 2100 Bioanalyzer (Agilent).

Polyadenylated RNA-seq library construction and sequencing using Illumina® technology were performed by the GENEWIZ® Company (Germany). Four hundred ng of total RNA at a concentration of 10 ng/μL, and OD260/280 comprised between 1.85 and 2.21 were used for library preparation. The NEBnext® Ultra^TM^ II Directional RNA kit was used for the cDNA library preparation and 9 cycles of enrichment PCR were run. Sequencing was performed on Illumina® NovaSeq^TM^, with a 150bp paired-end configuration, and a sequencing depth of 100M raw paired-end reads per sample.

### 5.3 Additional samples

Sixteen additional mussel individuals were used to validate our results. Ten mussels (5 MtrBTN2-affected and 5 MtrBTN2-free) were collected in a distinct location, at Le Croisic (47°17′58.5′′N 2°31′00.4′′W, Loire-Atlantique, France) in February 2023 and used to validate RNAseq-based quantification and differential expression between MtrBTN2 cells and normal haemocytes by RT-qPCRs (see Section 5.5). Total RNA extraction was performed as previously described (see Section 5.2). RNAseq reads from 6 mussels (3 MtrBTN2-affected and 3 MtrBTN2-free) collected in Agon-Coutainville (49°0′44.784′′N 1°35′55.643′′O, Normandy, France) in February 2021 for a separate study were used to confirm the differential transcriptomic profile of MtrBTN2 (see Section 5.5) and to validate the MtrBTN2-specific SNPs (see Section 5.6). Whole flesh from these 6 individuals was individually frozen in liquid nitrogen and powdered by cryogenic grinding (Mixer Mill MM 400, Retsch). For RNA extraction, mussel powder (10⍰mg) was homogenised in 500 µL of TRIzol by vortexing for 1⍰h at 4⍰°C. Prior to extraction, performed as described above, insoluble material was removed by centrifugation at 12,000 × g for 10⍰min at 4⍰°C.

### 5.4 De novo transcriptome assembly and functional annotation

Raw reads were processed using RCorrector v1.04 (https://anaconda.org/bioconda/rcorrector/files?version=1.0.4) with default settings and -rf configuration to correct sequencing errors (123). Uncorrectable reads were then removed using the FilterUncorrectabledPEfastq tool (https://github.com/harvardinformatics/TranscriptomeAssemblyTools/blob/master/FilterUn correctabledPEfastq.py). The output reads were further processed for adapter removal and trimming with TrimGalore! v0.6.4 (https://github.com/FelixKrueger/TrimGalore) with default parameters and -q 28, --length 100. Ribosomal RNAs potentially still present after polyA capture were removed by alignment against the SILVA Ribosomal database with Bowtie2 v2.4.1 (124). Read quality was assessed before and after read trimming using FastQC v0.11.9 (https://www.bioinformatics.babraham.ac.uk/projects/fastqc/).

In France, MtrBTN2 infects *M. edulis* hosts, but the lineage originated in a *M. trossulus* founder host. These two species are closely related, hybridise when they come into contact, either naturally or via human-induced introductions (125–127), and introgression between them is observed. However, *M. trossulus* is not present in France and individuals of this species have been sampled in a different environment (Barent Sea *vs* Channel Sea for MtrBTN2). Therefore, to allow a differential gene expression analysis of the cancer cells, all retained reads (from MtrBTN2 cells, *M. edulis* haemocytes and *M. trossulus* haemocytes) were assembled into a pantranscriptome using Trinity v2.8.5 (128) with the default options. TransDecoder v3.0.1 (https://github.com/TransDecoder/TransDecoder/wiki) was run on these contigs to identify CDSs with a minimum length of 90 amino acids. Finally, CDS sequence redundancy was reduced using CD-HIT-EST v4.8.1 (https://github.com/weizhongli/cdhit/wiki) with the following options: -n 6, -c 0.86, -G 0, -aL 1, -aS 1. We assessed the quality of the assembly using several tests. The assembly was first analysed using TrinityStats, and the final pantranscriptome completeness was estimated using BUSCO v5.1.1 (129) against the conserved single-copy metazoan gene database (n = 978). Finally, filtered reads were mapped back to the filtered pantranscriptome to assess the individual mapping rate with Bowtie2 v2.4.1 (124). To conclude that our pantranscriptome strategy was reliable, we performed a Redundancy Analysis (RDA) on the normalised read counts to analyse the impact of explanatory variables (“cell type”: haemocyte/MtrBTN2 cells, “species”: trossulus/edulis, “environment”: Barents Sea/Channel Sea) on response variables (gene expression), followed by an ANOVA-like permutation test (nperm=999, model=”full”) (https://cran.r-project.org/web/packages/vegan/vegan.pdf) (supplementary Figure S1). Both “cell type” and “environment” were retained as significant explanatory variables (p<0.05). This confirms that the inclusion of healthy *M. edulis* mussels in the analysis is necessary to subtract the environment effect (Channel Sea *vs* Barent Sea).

For functional annotation, the transcripts were searched against the Uniprot (Swissprot and TrEMBL) protein reference database (130) using PLASTX v2.3.3 algorithm with an e-value cutoff of 1.0E-3 (131). Domain prediction against the InterPro database (132) was carried out with InterProScan v5.48-83.0. Both results were combined and we used the OmicsBox program v2.0 (34,35) to assign GO terms to the annotated sequences with an e-value hit filter of 1.10−3, an annotation cutoff of 55, a HSP-Hit coverage of 60%, and an evidence code of 0.8. We searched for additional correspondences using EggNog v5.0 (133) by an orthology analysis.

### 5.5 Quantification of transcript expression

To investigate differential expression between MtrBTN2 cells and *M. edulis-M. trossulus* haemocytes, reads were first aligned to the pantranscriptome using SALMON v1.6 (134). DETs were identified using DESeq2 v1.34 (36) with a FDR<0.05. Count normalisation was performed using the RLE method implemented in the DESeq2 package (135). To confirm the identified DETs, we ran the QuantEval pipeline (https://github.com/dn070017/QuantEval, 37) with the “contig mode” and DESeq2 on the quantification outputs. We compared the quantification results by Spearman’s correlation analysis and we retained the shared DETs for the subsequent analyses.

To validate the quantifications and differential expression profiles obtained from our RNA-seq data, we carried out two alternative analyses on additional samples (see Section 5.3). First, we performed RT-qPCRs targeting 17 genes (*ADAMTS1, ALOX5, APC, BMP2, CCND1, CTNNB1, CYP11A1, FBXW7, JAG1, MAD3, PIK3R1, PTEN, SAV1, SELP, SFRP1, SFRP3,* and *THBS1*) selected for their contrasting expression levels according to the DESeq2 analysis and their oncogenic relevance. Reverse transcription into cDNA was performed using M-MLV Reverse Transcriptase (MMLV-RT, invitrogen) following the manufacturer’s instructions. The total RT-qPCR reaction volume was 1.5⍰μL and consisted of 1⍰μL of LightCycler 480 SYBR Green I Master Mix (Roche) containing 0.5⍰μM of PCR primer (Eurogenetec) and 0.5⍰μl of cDNA (1/10 dilution). The amplification efficiency of each primer pair (supplementary Table S7) was validated by serial dilution of a pool of all cDNAs. Relative expression was calculated as the threshold cycle (Cq) values of selected genes minus the mean of the measured threshold cycle (Cq) values of three constitutively expressed genes (*GAPDH, RPS28,* and *UBE2D3*) (supplementary Table S8). We compared the Log2FC values obtained from the two quantification methods. Log2FCs for RT-qPCRs were calculated as follow:

FC= efficiency of the qPCR ^∣ mean of Ct of the 5 MtrBTN2-affected individuals - mean of Ct of the 5 MtrBTN2-free individuals∣

We also performed Spearman’s correlation analysis between normalised expression obtained from RNA-seq data (RLE values) and normalised Ct obtained from RT-qPCR. Second, we performed DESeq2 analysis on complementary RNA-seq data obtained from the whole flesh of 6 additional samples from a separate study (see Section 5.3) to confirm transcripts specific to the malignant state.

### 5.6 Biological interpretation of gene expression profiles

In the context of a non-model species, we used three different approaches to interpret the biological relevance of DETS.

#### Enrichment analysis

We performed a GO term enrichment analysis focusing on biological processes with the GO_MWU tool (https://github.com/z0on/GO_MWU) using adaptive clustering and a rank-based statistical test (Mann–Whitney U-test). We used the following parameters for adaptive clustering: largest⍰=⍰0.5; smallest⍰=⍰10; clusterCutHeight⍰=⍰0.5. We considered both the level of expression and the significance of the differential expression: we assigned the log2 fold change value to transcripts that were significantly differentially expressed (adjusted p⍰<⍰0.05), while assigning zero to the others. We considered as enriched a biological process with a FDR⍰<⍰1%. To display the results synthetically, we used the Enrichment Map v3.3.3 tool (136) in Cytoscape v3.9.1 (137). The intensity of the enrichment was evidenced in the network and was calculated as follows: i) for the processes enriched with over-expressed transcripts, “number of transcripts significantly over-expressed in the process/total number of transcripts in the process”; (ii) for the processes enriched with under-expressed transcripts, “−1⍰×⍰(number of transcripts significantly downregulated in the process/total number of transcripts in the process)”.

#### Hub and top expressed gene identification

The top 20 genes in terms of connectivity ranking in the PPI network were selected as Hub genes. We used the Search Tool for the Retrieval of Interacting Genes (STRING), a biological database designed to predict PPI networks (138). The results were visualised in Cytoscape v3.9.1 using NetworkAnalyzer v4.4.8 visualisation software (139), which can construct comprehensive models of biological interactions. Isolated and partially connected nodes were not included.

To identify marker genes of the cancerous condition, we considered both differential expression (Log2FC) and expression level (RLE) values to build a plot graph. We defined arbitrarily thresholds (log2FC values > |5|and RLE > 500) to highlight the most discriminating genes between the two conditions (cancerous/healthy circulating cells).

#### Targeted analyses

We performed a targeted analysis focusing on genes and pathways that have been identified by Sanchez-Vega et al. (26) as being altered at high frequency across many different human cancer types. We searched for the presence of a differential expression of these genes in MtrBTN2 cells or of MtrBTN2-specific alleles (i.e. present only in MtrBTN2-affected mussels, and present in all MtrBTN2-affected mussels). We also identified *M. edulis* and *M. trossulus*-specific alleles. Allele visualisation and calculation of variant frequencies were carried out using the IGV v2.12.3 (140) and iVar (141) tools on BAM files obtained after read alignment to the transcript sequences with Bowtie2. v2.4.1 (124). The minimum quality score threshold to count a base and the minimum frequency threshold were set to 20 and 0.02, respectively, with a minimum base coverage of 50 reads. MtrBTN2-specific alleles were searched in CAN1-3 individuals and then confirmed in additional CAN4-6 individuals (see Section 5.3).

## Acknowledgments

We acknowledge the Animal Health-Clinical Biology unit of the LABÉO laboratory (Caen, France), especially Maryline Houssin and Ludovic Petit, for their help during cytological screening and Professor Petr Strelkov from the Saint Petersburg State University for the sampling of *M. trossulus* individuals. We thank the “Réseau d’Histologie Expérimentale de Montpellier” - RHEM facility - supported by SIRIC Montpellier Cancer (Grant INCa_Inserm_DGOS_12553), the European regional development foundation and the Occitanian region (FEDER-FSE 2014-2020 Languedoc Roussillon), for processing our animal tissues, histology technics and expertise. This study is set within the framework of the ‘Laboratoires d’Excellence (LABEX)’ Tulip (ANR-10-LABX-41).

## Author contributions

Conceptualization: EAVB, DDG, GM, GMC, EV, JVD; Investigation, Formal analysis: EAVB, MH; Methodology: EAVB, EV, JVD; Funding Acquisition: DDG, GMC, NB, FT; Original Draft preparation: EAVB; Reviewing&Editing: DDG, GMC, MH, EV, JVD, NB, FT, GM.

## Data and materials availability

Raw read sequences were deposited with links to BioProject accession number PRJNA749800 in the NCBI BioProject database (https://www.ncbi.nlm.nih.gov/bioproject/).

