## Supplementary figures and images for "Transcriptomics of mussel transmissible cancer MtrBTN2 suggests accumulation of multiple cancerous traits and oncogenic pathways shared among bilaterians"

### Supplementary Figure S1

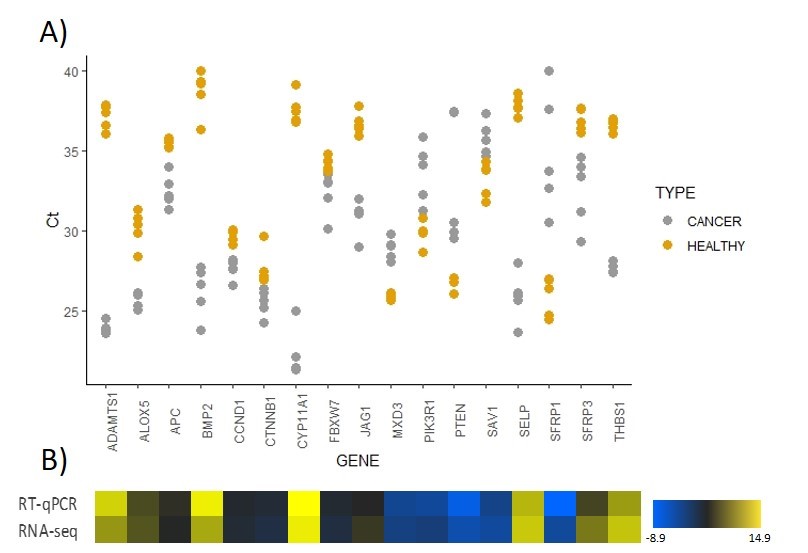

### Supplementary Figure S2

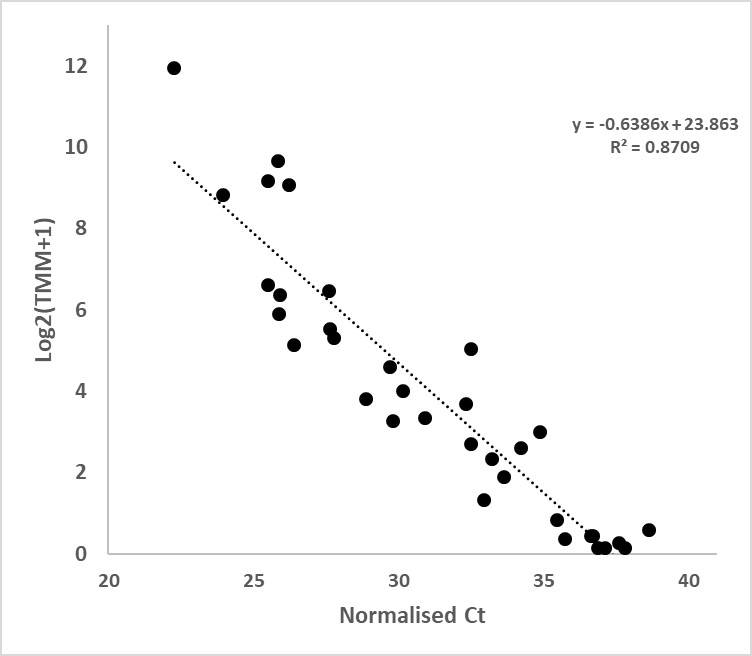

### Supplementary Figure S3

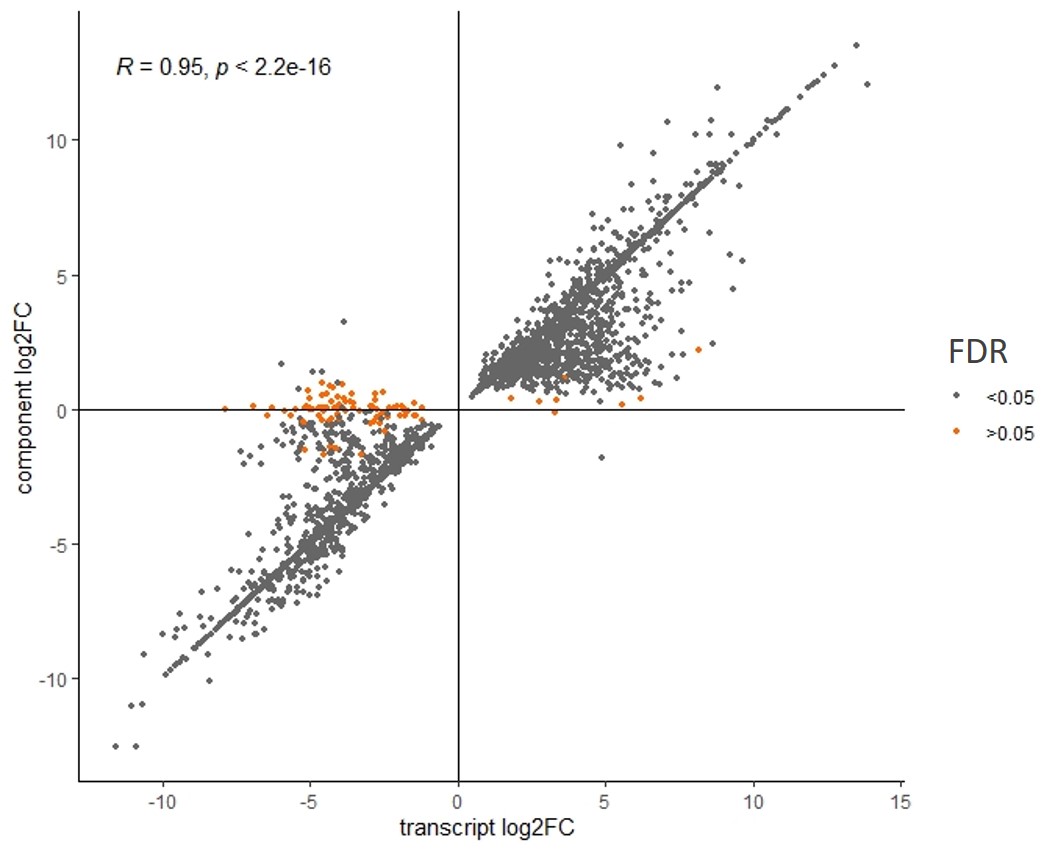

### Supplementary Figure S4

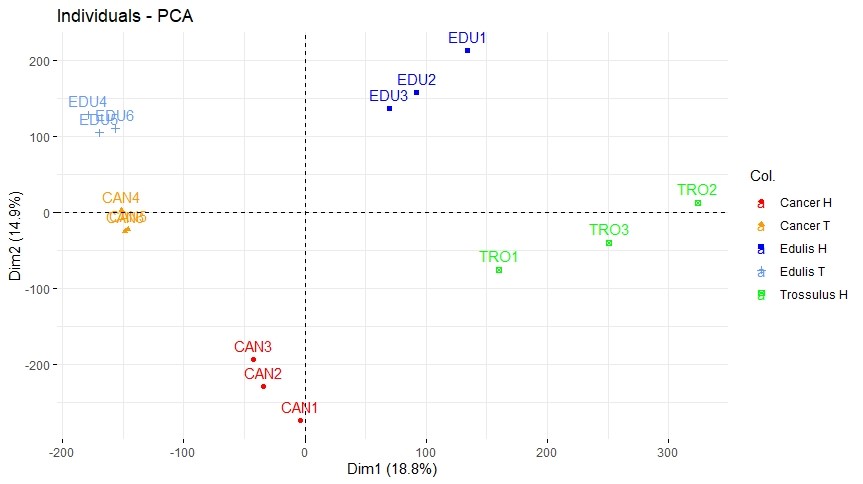

### Supplementary Figure S5

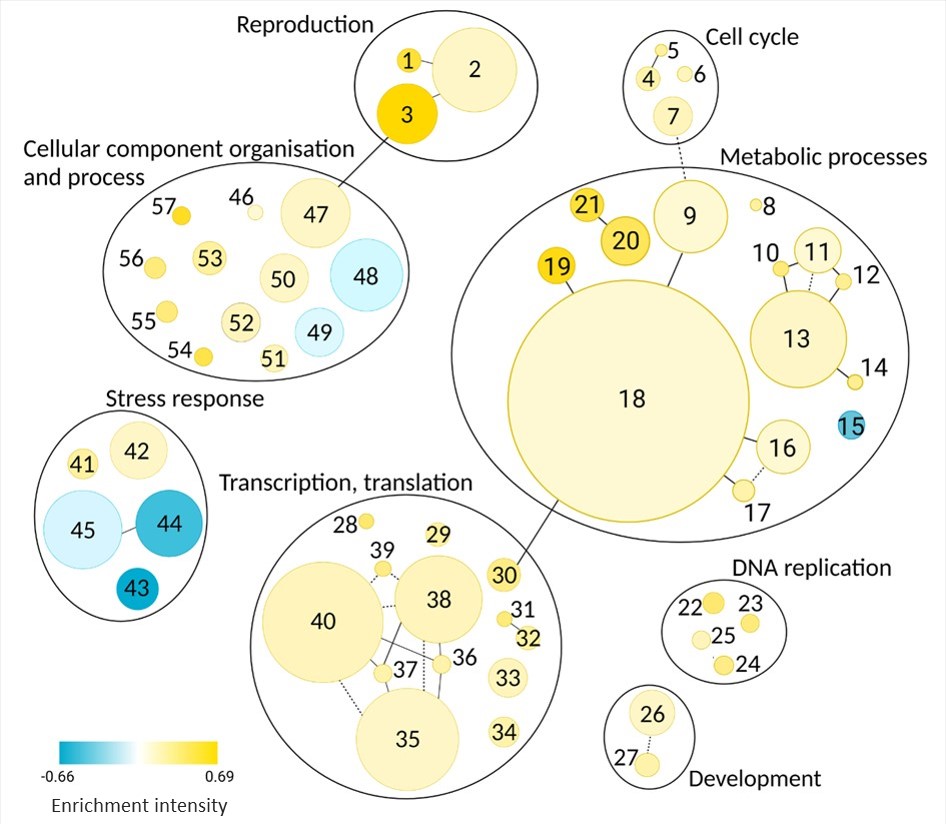

### Supplementary Figure S6

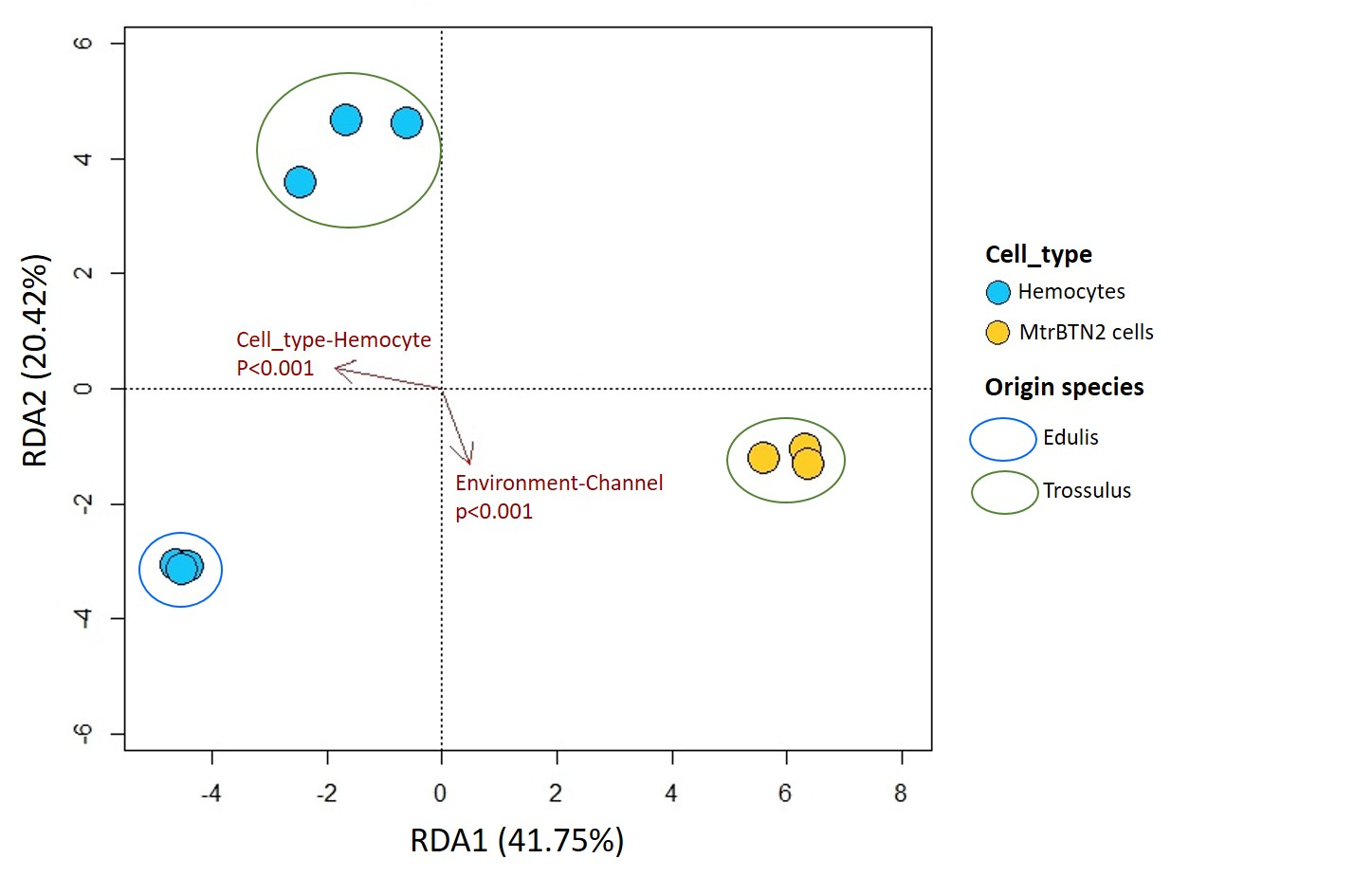
