## Supplementary File S1 for "Transcriptomics of mussel transmissible cancer MtrBTN2 suggests accumulation of multiple cancerous traits and oncogenic pathways shared among bilaterians"

**ALIGNMENTS PIK3CA-GENE_NUCLEOTIDE SEQUENCES**

```
PIK3CA_EDULIS                       ATGCCTCCTAGTTCAGGAGAACTATGGGGTCATCATTTGATGCCCTCAAC
PIK3CA_TROSSULUS                    ATGCCACCAAGTTCGGGGGAATTATGGGGTCATCATTTGATGCCCTCAAC
PIK3CA_CANCER                       ATGCCACCAAGTTCGGGGGAATTATGGGGTCATCATTTGATGCCCTCAAC
                                    ***** ** ***** ** *** **************************** 50

PIK3CA_EDULIS                       TATTCAGGTGGACTGTTTATTACCCACAGGGATCATTATTCAGGTGGCTG
PIK3CA_TROSSULUS                    TATTCAGGTGGACTGTTTATTGCCCACAGGGATCATTATTCAGGTGGCTG
PIK3CA_ CANCER                      TATTCAGGTGGACTGTTTATTGCCTACAGGGATCATTATTCAGGTGGCTG
                                    ********************* ** ************************* 100

PIK3CA_EDULIS                       TGAGTCGTGACGAAACACTTGAACGGATTAAAGCAGATCTTTGGGTAAAG
PIK3CA_TROSSULUS                    TGAGCCGTGACGAAACACTGGAACGGATTAAAGCCGATCTTTGGGTAAAG
PIK3CA_ CANCER                      TGAGCCGTGATGAAACACTGGAAAGGATTAAAGCCGATCTTTGGGTAAAG
                                    **** ***** ******** *** ********** *************** 150

PIK3CA_EDULIS                       GCAAAGCTTTATCCTCTTTATGAAAGACTATTAGAACCTGCATCATACAT
PIK3CA_TROSSULUS                    GCAAAGCTTTATCCTCTTTATGAAAGACTATTAGAACCTGCATCATACAT
PIK3CA_CANCER                       GCAAAGCTTTATCCTCTTTATGAAAGACTATTAGAACCTGCATCATACAT
                                    ************************************************** 200

PIK3CA_EDULIS                       ATTTGTGAGCATTACCCAAGATGCGCGGAAAGAGGAGTTTTATGATGAAA
PIK3CA_TROSSULUS                    ATTTGTGAGCATTACCCAGGATGCACGGAAAGAGGAGTTTTATGATGAAA
PIK3CA_ CANCER                      ATTTGTAAGCATTACCCAGGATGCACGGAAAGAGGAGTTTTATGATGAAA
                                    ****** *********** ***** ************************* 250

PIK3CA_EDULIS                       CAAGACGATTTTGTGATTTACGCTTGTTTCAACCTATTCTGAAAATTGTT
PIK3CA_TROSSULUS                    CAAGACGATTTTGTGATTTACGATTATTTCAACCTATTCTGAAAATTGTT
PIK3CA_ CANCER                      CAAGACGATTTTGTGATTTACGATTATTTCAACCTATTCTGAAAATTGTT
                                    ********************** ** ************************ 300

PIK3CA_EDULIS                       GAACCAGTTGGAAACAGAGAGGAGAAAATGTTGAATTTTGAAATAGGTGT
PIK3CA_TROSSULUS                    GAACCAGTTGGAAACAGGGAGGAGAAAATGCTGAATTTTGAAATAGGTGT
PIK3CA_ CANCER                      GAACCAGTTGGAAACAGGGAGGAGAAAATGCTGAATTTTGAAATAGGTGT
                                    ***************** ************ ******************* 350

PIK3CA_EDULIS                       AACGATAGGTGTTTCTATAAATGAATTTAATGAAATGAAAGACTTAGAAG
PIK3CA_TROSSULUS                    AACGATAGGTGTATCTATAAATGAATTTAATGAAATGAAAGACTTAGAAG
PIK3CA_ CANCER                      AACGATAGGTGTATCTATAAATGAATTTAATGAAATGAAAGACTTAGAAG
                                    ************ ************************************* 400

PIK3CA_EDULIS                       TGATGACATTCCGGAGGAATATTTTAGACATTTGTAAGAATGCAATAGAT
PIK3CA_TROSSULUS                    TGATGACATTCCGGAGGAACATTTTAGACATTTGTAAAAATGCAATAGAT
PIK3CA_ CANCER                      TGATGACATTCCGGAGGAACATTTTAGACATTTGTAAAAATGCAATAGAT
                                    ******************* ***************** ************ 450

PIK3CA_EDULIS                       GAGCGACAGCGACGGGGTAAGGCAAGCGAGGCCTTATATGCCTACCCACC
PIK3CA_TROSSULUS                    GAGCGTCAGCGACGGGGTAAGGCAAGCGAGGCCTTATATGCCTACCCACC
PIK3CA_ CANCER                      GAGCGTCAGCGACGTGGTAAGGCAAGCGAGGCCTTATATGCCTACCCACC
                                    ***** *******  *********************************** 500

PIK3CA_EDULIS                       AGATATAGAATCATCACCTATTTTACCGTCACATTTACAGACAAAAATTA
PIK3CA_TROSSULUS                    AGATATAGAATCATCACCTATTTTACCCTCACATTTACAGACAAAAATTA
PIK3CA_ CANCER                      AGATATAGAATCGTCACCTATTTTACCCTCACATTTACAGACAAAAATTA
                                    ************ ************** ********************** 550

PIK3CA_EDULIS                       AAGAAATGAAAAATGATGTAGTGGCATGTGTTTGGGTTGTATCAGAAGAC
PIK3CA_TROSSULUS                    AAGAAATGAAAAATGATGTAGTGGCATGTGTTTGGGTTGTATCAGAAGAC
PIK3CA_ CANCER                      AAGAAATGAAAAATGATGTAGTGGCATGTGTTTGGGTTGTATCAGAAGAC
                                    ************************************************** 600

PIK3CA_EDULIS                       AATAGTCGTAGTAAATTTAGTGTTAAAGTTTCACATGATGCTTATCCAAT
PIK3CA_TROSSULUS                    AATAGTCGTAGTAAATTTAGTGTAAAAGTTTCACATGACGCTTATCCAAT
PIK3CA_ CANCER                      AATAGTCGTAGTAAATTTAGTGTAAAAGTTTCACATGACGCTTATCCAAT
                                    *********************** ************** *********** 650

PIK3CA_EDULIS                       TGATGTTATTGCTGGAACAATACGACGGAGGAGCAGAATGATGGGAATTT
PIK3CA_TROSSULUS                    TGACGTAATTGCTGGAACAATACGACGGAGGAGCAGAATGATGGGAATTT
PIK3CA_ CANCER                      TGACGTAATTGCTGGAACAATACGACGGAGGAGCAGAATGATGGGAATTT
                                    *** ** ******************************************* 700

PIK3CA_EDULIS                       CTAAGGAACACGCTGAACGTTGTATAGAAGAATATAGTGACACTTATGCA
PIK3CA_TROSSULUS                    CTAAGGAACACGCTGAACGTTGTATAGAAGAATATAGTGACACTTATGCA
PIK3CA_ CANCER                      CTAAGGAACACGCTGAACGTTGTATAGAAGAATATAGTGACACTTATGCA
                                    ************************************************** 750

PIK3CA_EDULIS                       CTTAAAGTGTGTGGTTGTGATCAATTCTTATTAGAGGAGCATCCGTTAAG
PIK3CA_TROSSULUS                    CTTAAAGTGTGTGGTTGTGATCAATTCTTATTAGAGGAGCATCCGTTAAG
PIK3CA_ CANCER                      CTTAAAGTCTGTGGTTGTGATCAATTCTTATTAGAGGAGCATCCGTTAAG
                                    ******** ***************************************** 800

PIK3CA_EDULIS                       TCAGTATAAGTATGTGAGAGAATGTATAGCGAGAGATAATATACCACAGT
PIK3CA_TROSSULUS                    TCAGTATAAGTATGTGAGAGAATGTATAGCAAGAGATAATATTCCACAGT
PIK3CA_ CANCER                      TCAGTATAAGTATGTTAGAGAATGTATAGCAAGAGATAATATTCCACAGT
                                    *************** ************** *********** ******* 850

PIK3CA_EDULIS                       TTATGTTATTAACAAAGGAAAGTTTATATGCAGCGATATCACCTAATATA
PIK3CA_TROSSULUS                    TTATGTTATTAACAAAGGAAAGTTTATATGCAGCGATATCACCTAATATA
PIK3CA_ CANCER                      TTATGTTATTAACAAAGGAAAGTTTATATGCAGCGATATCACCTAATATA
                                    ************************************************** 900

PIK3CA_EDULIS                       TTTATAACACCATCATATACACAGAAAGGGATGCAGTTACTAACAGAAAT
PIK3CA_TROSSULUS                    TTTATAACACCATCATATACACAGAAAGGTATGCAGTTATTAACAGAAAT
PIK3CA_ CANCER                      TTTATAACACCATCATATACACAGAAAGGTATGCAGTTATTAACAGAAAT
                                    ***************************** ********* ********** 950

PIK3CA_EDULIS                       TAACCAACAAAAGACATTATCTCTGTGGGAAATTCATGCCAAACTTAGAA
PIK3CA_TROSSULUS                    TAACCAACAAAAGACATTATCTCTGTGGGAAATTCATGCCAAACTGAGAA
PIK3CA_ CANCER                      TAACCAACAAAAGACATTATCTCTGTGGGAAATTCATGCCAAACTGAGAA
                                    ********************************************* **** 1000

PIK3CA_EDULIS                       TTAAAATATTATGTGCCACTTATGTCAATGTAAAAGAACTTGGAAAGATA
PIK3CA_TROSSULUS                    TTAAAATATTATGTGCCACTTATGTCAATGTAAAAGAACTTGGAAAGATA
PIK3CA_ CANCER                      TTAAAATATTATGTGCCACTTATGTCAATGTAAAGGAACTTGGAAAGATA
                                    ********************************** *************** 1050

PIK3CA_EDULIS                       TATGTTAAAGCAGGAATTTACCATGGAACAGAAGCTCTGTGTGAATTCCA
PIK3CA_TROSSULUS                    TATGTTAAAGCAGGAATTTACCATGGAACAGAAGCTCTGTGTGAATTCCA
PIK3CA_ CANCER                      TATGTTAAAGCAGGAATTTACCATGGAACAGAAGCTCTGTGTGAATTCCA
                                    ************************************************** 1100

PIK3CA_EDULIS                       AGATACAAAAATGGTTGACTCAAACAATCCCCAGTGGCATGAATGGTTAG
PIK3CA_TROSSULUS                    AGATACAAAGATGGTTGACTCAAACAACCCCCAGTGGCATGAGTGGTTAG
PIK3CA_ CANCER                      AGATACAAAGATGGTTGACTCAAACAACCCCCAGTGGCATGAGTGGTTAG
                                    ********* ***************** ************** ******* 1150

PIK3CA_EDULIS                       AGTTTTTGTACATACAGGACTTACCTAGAAGTGCTAAACTATGTTTATCT
PIK3CA_TROSSULUS                    AGTTCTTGTACATACAGGATTTACCTAGAAGTGCTAAACTATGTTTATCT
PIK3CA_ CANCER                      AGTTCTTGTACATACAGGATTTACCTAGAAGTGCTAAACTATGTTTATCT
                                    **** ************** ****************************** 1200

PIK3CA_EDULIS                       ATTTGCTATACATCCAAGAAAAAGAGGGAACCCTTATCTTTAGGATGGGC
PIK3CA_TROSSULUS                    ATATGTTATACATCTAAGAAAAAGAGGGAACCCTTATCTTTAGGATGGGC
PIK3CA_ CANCER                      ATATGTTATACATCTAAGAAAAAGAGGGAACCCTTATCTTTAGGATGGGC
                                    ** ** ******** *********************************** 1250

PIK3CA_EDULIS                       CAATTTGCAGCTATTTGATTTCAACGATAGACTTATGAACGAAAAGGTCA
PIK3CA_TROSSULUS                    TAATCTGCAGCTATTTGATTTCAACGACAGACTTATGAATGAAAAAGTCA
PIK3CA_ CANCER                      TAATCTGCAGCTATTTGATTTCAACGACAGACTAATGAATGAAAAAGTCA
                                     *** ********************** ***** ***** ***** **** 1300

PIK3CA_EDULIS                       GCTTGAATCTGTGGCCGTTACCTCAGGGAATGGATGACCTCTTGAATTAT
PIK3CA_TROSSULUS                    GCTTGAATCTATGGCCATTACCTCAGGGAATGGATGACCTATTGAACTAT
PIK3CA_ CANCER                      GCTTGAATCTATGGCCATTACCTCAGGGAATGGATGACCTATTGAACTAT
                                    ********** ***** *********************** ***** *** 1350

PIK3CA_EDULIS                       GTTGGTCTCCCAGGTTCCAATCCAGACCAAGAAACCCCATGTTTAGAGAT
PIK3CA_TROSSULUS                    GTTGGTCTTCCAGGTTCCAATCCTGACCAAGAAACTCCATGTTTAGAGAT
PIK3CA_ CANCER                      GTTGGTCTTCCAGGGTCCAATCCTGACCAAGAAACTCCATGTTTAGAGAT
                                    ******** ***** ******** *********** ************** 1400

PIK3CA_EDULIS                       AGAGTTTGACAGACTAAGTCATCCTGTGTCATATCCTCCAGAGAAACAGA
PIK3CA_TROSSULUS                    AGAGTTTGACAGACTAAGTCACCCTGTGTCATATCCTCCAGAGAAACAGA
PIK3CA_ CANCER                      AGAGTTTGACAGACTAAGTCACCCTGTATCATATCCTCCAGAGAAACAGA
                                    ********************* ***** ********************** 1450

PIK3CA_EDULIS                       TTGAAGATTTAGCTCAATACGCCATCAGCAAAGAACAACCACTCATTTAT
PIK3CA_TROSSULUS                    TTGAAGATTTAGCTCAATACGCCATCAGCAAAGAATCCCCCCTTATTTAT
PIK3CA_ CANCER                      TTGAAGATTTAGCTCAATACGCCATCAGCAAAGAATCCCCCCTTATTTAT
                                    ***********************************   ** ** ****** 1500


PIK3CA_EDULIS                       CTGGACACCCCCCAAATACAAGCAAAAGAGCAAGCTATGGTGGATGATAT
PIK3CA_TROSSULUS                    CTGGACACCCCACAAATACAAGCAAAAGAGCAAGCTATGGTGGATGATAT
PIK3CA_ CANCER                      CTTGACACCCCACAAATACAAGCAAAAGAGCAAGCTATGGTGGATGATAT
                                    ** ******** ************************************** 1550

PIK3CA_EDULIS                       AACAAGTAGAGATCCACTATCAGAAATATCAGAACAAGAAAAAGATATTC
PIK3CA_TROSSULUS                    AACCAGTAGAGATCCACTATCAGAAATATCAGAACAAGAAAAAGATATTC
PIK3CA_ CANCER                      AACCAGTAGAGATCCACTATCTGAAATATCAGAACAAGAAAAAGATATTC
                                    *** ***************** **************************** 1600

PIK3CA_EDULIS                       TATGGAAATTAAGAGAATACTGTATCCAAGTTCCTCAATCCTTACCTAAA
PIK3CA_TROSSULUS                    TATGGAAATTAAGAGAATACTGTATCCGAGTTCCTCAATCCTTACCTAAA
PIK3CA_ CANCER                      TATGGAAATTAAGAGAATACTGTATCCGAGTTCCTCAATCCTTACCTAAA
                                    *************************** ********************** 1650

PIK3CA_EDULIS                       CTCCTACAGTCAGTGAAATGGAATGAGCGGGAGTATGTAGCTCAGCTGTA
PIK3CA_TROSSULUS                    CTCCTACAGTCAGTAAAATGGAATGAGCGGGAGTACGTAGCTCAGTTGTA
PIK3CA_ CANCER                      CTCCTACAGTCAGTAAAATGGAATGAGCGGGAGTACGTAGCTCAGTTGTA
                                    ************** ******************** ********* **** 1700

PIK3CA_EDULIS                       CATGTTGTTACGACGATGGCCCCGTCTGATTCCAGAATTTGCCATGGAGC
PIK3CA_TROSSULUS                    CATGCTGTTACGACGATGGCCCAGATTGATGCCAGAGTTTGCCATGGAAT
PIK3CA_ CANCER                      CATGCTGTTACGACGATGGCCCAGATTGATGCCAGAGTTTGCCATGGAAT
                                    **** ***************** *  **** ***** ***********   1750

PIK3CA_EDULIS                       TGCTTGATTGTTCTTACCCAGATCTGTGTGTGCGTCAGTATGCTGTTACC
PIK3CA_TROSSULUS                    TGCTTGATTGTTCCTACCCAGATCTGTGTGTGCGTCAGTATGCTGTTACC
PIK3CA_ CANCER                      TGCTTGATTGTTCCTACCCAGATCTGTGTGTGCGTCAGTATGCAGTTACC
                                    ************* ***************************** ****** 1800

PIK3CA_EDULIS                       TGTCTGGATCATGGATTCTCTGATGATAAATTACAACAGTATATGTTACA
PIK3CA_TROSSULUS                    TGTCTGGATCATGGATTTTCTGATGATAAATTACAGCAGTATATGTTACA
PIK3CA_ CANCER                      TGTCTGGATCATGGATTTTCTGATGATAAATTACAGCAGTATATGTTACA
                                    ***************** ***************** ************** 1850

PIK3CA_EDULIS                       GTTAGTGCAGGCGCTGAAATTTGAGCCGTATTTAGACAGTCCCATCACAA
PIK3CA_TROSSULUS                    ATTAGTGCAGGCGCTGAAATTTGAGCCGTATTTAGACAGTCCCATCACAA
PIK3CA_ CANCER                      ATTAGTGCAGGCGCTGAAATTTGAGCCGTATTTAGACAGTCCCATCACAA
                                     ************************************************* 1900

PIK3CA_EDULIS                       GATTTCTTCTAAAGAGAGCTCTTCAAAATCAGAAAACAGGACAGCTGTTC
PIK3CA_TROSSULUS                    GATTTCTTCTAAAGAGAGCTCTACAAAATCAGAAAACAGGGCAGCTGTTC
PIK3CA_ CANCER                      GATTTCTTCTAAAGAGAGCTCTACAAAATCAGAAAACCGGGCAGCTGTTC
                                    ********************** ************** ** ********* 1950

PIK3CA_EDULIS                       TTCTGGCACCTTAAGTCAGAAATCCATAATACATCAATCCAGTTAAGATT
PIK3CA_TROSSULUS                    TTCTGGCACCTTAAGTCAGAAATCCATAATACATCAATCCAGTTAAGATT
PIK3CA_ CANCER                      TTCTGGCACCTTAAGTCTGAAATCCATAATACATCAATCCAGTTAAGATT
                                    ***************** ******************************** 2000

PIK3CA_EDULIS                       TGGTTTGGTGTTAGAAGCATTCTGTCGAGGGTGTGGATCCAATCTCAAGA
PIK3CA_TROSSULUS                    TGGTTTGGTGTTAGAAGCATTCTGTCGAGGGTGTGGATCCAATCTCAAGA
PIK3CA_ CANCER                      TGGTTTGGTGTTAGAAGCATTTTGTCGAGGGTGTGGATCCAATCTCAAGA
                                    ********************* **************************** 2050

PIK3CA_EDULIS                       TGTTACTGAGACAAGTGGAGGCTCTAGATAAACTCACCAAACTTACTAAT
PIK3CA_TROSSULUS                    TGTTACTGAGACAAGTGGAGGCTCTAGATAAACTCACCAAACTTACAAAT
PIK3CA_ CANCER                      TGTTACTGAGACAAGTCGAGGCTCTAGATAAACTCACCAAACTTACAAAT
                                    **************** ***************************** *** 2100

PIK3CA_EDULIS                       ATGATCAAAAATGAAGTCAAGGATGACATTAATGAAATTATGAAGTTTCT
PIK3CA_TROSSULUS                    ATGATCAAAAATGAAGTCAAAGATGACATTAATGAAATTATGAAGTTTCT
PIK3CA_ CANCER                      ATGATCAAAAATGAAGTCAAAGATGACATTAATGAAATTATGAAGTTTCT
                                    ******************** ***************************** 2150

PIK3CA_EDULIS                       TGCTAACCAGTTACAGCAACCAGACTACCAAGATGGACTGAAAAACTTCT
PIK3CA_TROSSULUS                    TGCTAACCAGTTACAGCAACCAGACTACCAAGATGGACTGAAAAACTTCT
PIK3CA_ CANCER                      TGCTAACCAGTTACAGCAACCAGACTACCAGGATGGGCTGAAAAACTTCT
                                    ****************************** ***** ************* 2200

PIK3CA_EDULIS                       TATCTCCCCTTGATAACAGTCATATACTAGGAGATCTAGATATTTCGCAT
PIK3CA_TROSSULUS                    TATCTCCCCTTGATAACAGTCATATACTAGGAGATCTAGATATTTCGCAT
PIK3CA_ CANCER                      TATCTCCCCTTGATAACAGTCATATACTAGGAGACCTAGATATTTCGCAT
                                    ********************************** *************** 2250


PIK3CA_EDULIS                       TGTTCAGTAATGACATCTAAAAAGAAACCATTGTGGCTAGTGTTTAGTAA
PIK3CA_TROSSULUS                    TGTTCAGTAATGACATCTAAAAAGAAACCATTGTGGTTAGTGTTTAGTAA
PIK3CA_ CANCER                      TGTTCAGTAATGACATCTAAAAAGAAACCATTGTGGTTAGTGTTTAGTAA
                                    ************************************ ************* 2300

PIK3CA_EDULIS                       CCCAGATGTCATGGCTGATATATGGTTTACAGACTACAAACTTATATTTA
PIK3CA_TROSSULUS                    CCCAGATGTCATGGCTGATATATGGTTTACAGACTACAAACTTATATTTA
PIK3CA_ CANCER                      CCCAGATGTCATGGCTGATATATGGTTTACAGACTACAAACTTATATTTA
                                    ************************************************** 2350

PIK3CA_EDULIS                       AAAATGGAGATGATTTACGTCAGGACATGCTTACTGTACAGTTATTTAAA
PIK3CA_TROSSULUS                    AAAATGGAGATGATTTGCGTCAGGACATGCTTACTGTACAGTTATTTAAA
PIK3CA_ CANCER                      AAAATGGAGATGATCTGCGTCAGGACATGCTTACTGTACAGTTATTTAAA
                                    ************** * ********************************* 2400

PIK3CA_EDULIS                       ATTATGGACACTTTATGGAAAAATGAAGGTTTAGATCTGAGGTTGATACC
PIK3CA_TROSSULUS                    ATTATGGACACTTTATGGAAAAATGAAGGTTTAGATCTGAGGTTGATACC
PIK3CA_ CANCER                      ATTATGGACACTCTATGGAAAAATGAAGGTTTAGATCTGAGGTTGATACC
                                    ************ ************************************* 2450

PIK3CA_EDULIS                       GTATAGTGTGGTATCTACTGGCAAAGATGTAGGTGTGATAGAAATAGTAC
PIK3CA_TROSSULUS                    GTATAGTGTGGTATCTACTGGCAAAGATGTAGGTGTAATAGAAATAGTAC
PIK3CA_ CANCER                      GTATAGTGTGGTTTCTACTGGCAAAGATGTTGGTGTAATAGAAATAGTAC
                                    ************ ***************** ***** ************* 2500

PIK3CA_EDULIS                       GGGATTCTTCGACTATCATGAGTATACAACAGAAGAATGGTATCCGAGCA
PIK3CA_TROSSULUS                    GGGATTCCTCGACAATTATGAGTATACAACAGAAGAATGGTATCCGTGCA
PIK3CA_ CANCER                      GGGATTCCTCGACAATTATGAGTATACAACAGAAGAATGGTATCCGTGCC
                                    ******* ***** ** ***************************** **  2550

PIK3CA_EDULIS                       GCAGTACAGATGGATTCTCTAGGGTTGTATAACTGGATAATGTACCATAA
PIK3CA_TROSSULUS                    GCAGTACAGATGGATTCTCTAGGGTTGTATAACTGGATAATGTACCATAA
PIK3CA_ CANCER                      GCAGTTCAGATGGATTCTCTAGGGTTGTATAACTGGATAATGTACCATAA
                                    ***** ******************************************** 2600

PIK3CA_EDULIS                       TAAAGATAGAGAAGAACAAGCTATATCTAACTTCACAAGATCCTGTGCTG
PIK3CA_TROSSULUS                    TAAAGATAGAGAAGAACAAGCTATTTCTAACTTCACAAGATCCTGTGCTG
PIK3CA_ CANCER                      TAAAGATAGAGAAGAACAAGCTATTTCTAACTTCACAAGATCCTGTGCTG
                                    ************************ ************************* 2650

PIK3CA_EDULIS                       GTTACTGTGTAGCTACATTTATACTGGGTATCAAAGACAGACATAGTGGA
PIK3CA_TROSSULUS                    GTTACTGTGTAGCTACATTTATACTGGGTATTAAAGACAGACATAGTGGA
PIK3CA_ CANCER                      GTTACTGTGTAGCTACATTTATACTGGGTATTAAAGACAGACATAGTGGA
                                    ******************************* ****************** 2700

PIK3CA_EDULIS                       AATATAATGGTCAGGAAAAATGGACAGGTATTCCATATAGATTTTGGACA
PIK3CA_TROSSULUS                    AATATAATGGTCAGGAAAAATGGACAGGTATTCCATATAGACTTTGGACA
PIK3CA_ CANCER                      AATATAATGGTCAGGAAAAATGGACAGGTATTCCATATAGACTTTGGACA
                                    ***************************************** ******** 2750

PIK3CA_EDULIS                       TTTTTTGGATCATAGGAAAAAGAAATTTGGTATAACTCGAGAAAGAGTAC
PIK3CA_TROSSULUS                    TTTTTTGGATCATAGGAAAAAGAAATTTGGTATAACTCGAGAAAGAGTAC
PIK3CA_ CANCER                      TTTTTTGGATCATAGGAAAAAGAAATTTGGTATAACTCGAGAGAGAGTAC
                                    ****************************************** ******* 2800

PIK3CA_EDULIS                       CATTTGTATTGACAATGGACTTTATAAGAGTGATTGCTAGAGGATCAGAT
PIK3CA_TROSSULUS                    CATTTGTATTGACGATGGACTTTATAAGAGTAATTGCTAGAGGATCAGAT
PIK3CA_ CANCER                      CGTTTGTATTGACGATGGACTTTATAAGAGTAATTGCTAGAGGTTCTGAT
                                    * *********** ***************** *********** ** *** 2850

PIK3CA_EDULIS                       CAACCATTAAAGCACAAAGAATTTAAAAAATTTCAGCAACTCTGCTGTGA
PIK3CA_TROSSULUS                    CAACCACTAAAGCACAAAGAATTTAAAAAATTTCAGCAACTCTGCTGTGA
PIK3CA_ CANCER                      CAACCACTAAAGCACAAAGAATTTAAAAAATTTCAGCAACTCTGCTGTGA
                                    ****** ******************************************* 2900

PIK3CA_EDULIS                       AGCTTATTTAATTATTCGTAAAAATGCCTATTTATTCATCAATCTTATGA
PIK3CA_TROSSULUS                    AGCTTATTTAATTATTCGTAAAAATGCCTATTTATTCATCAATCTTATGA
PIK3CA_ CANCER                      AGCTTATTTAATTATTCGTAAAAATGCCTATTTATTCATCAATCTTATGA
                                    ************************************************** 2950

PIK3CA_EDULIS                       CTATGATGTTGTCGTGTGGTATCCCAGAGTTACAATCTCTAGATGACATA
PIK3CA_TROSSULUS                    CTATGATGTTGTCATGTGGTATCCCAGAGTTACAATCTCTAGATGACATA
PIK3CA_ CANCER                      CTATGATGTTGTCATGTGGTATCCCAGAGTTACAATCTCTTGATGACATA
                                    ************* ************************** ********* 3000


PIK3CA_EDULIS                       AGTTATCTTCGTAAAACATTAGCCGTGGAAGAGAAAGACGATGAAAAAGC
PIK3CA_TROSSULUS                    AGTTATCTTCGTAAAACATTAGCCGTGGAAGAGAAAGACGATGAAAAAGC
PIK3CA_ CANCER                      AGTTATCTTCGTAAAACATTAGCCGTGGAAGAGAAAGACGATGAAAAAGC
                                    ************************************************** 3050

PIK3CA_EDULIS                       CTTAAAATATTTCATAGCCAAGTTCAATAGTGCTTATTCCGATGCTTGGA
PIK3CA_TROSSULUS                    CTTAAAATATTTCATAGCCAAGTTTAATAGTGCTTATTCCGATGCTTGGA
PIK3CA_ CANCER                      CTTAAAATATTTCATAGCCAAGTTTAATAGTGCTTATTCCGATGCTTGGA
                                    ************************ ************************* 3100

PIK3CA_EDULIS                       CCGTCAAAACTGATTGGTTGTTCCATTATATGAAAAACAGATGA
PIK3CA_TROSSULUS                    CCGTCAAAACTGATTGGTTGTTCCATTATATGAAAAACAGATGA
PIK3CA_ CANCER                      CCGTCAAAACTGATTGGTTGTTCCATTATATGAAAAACAGATGA
                                    ******************************************** 3144
```

**ALIGNMENTS PIKACB-GENE_NUCLEOTIDE SEQUENCES**

```
PIK3CB_TROSSULUS      ATGCCTCCTGTGATAGTCTCCCCTGATCTGGATGTCACATCCTTCCAGAATGATCTTGGA
PIK3CB_MTRBTN2        ATGCCTCCTGTGATAGTCTCCCCTGATCTGGATGTCACATCCTTCCAGAATGATCTTGGA
PIK3CB_EDULIS         ATGCCTCCAGTGATAGTCTCCCCTGATCTGGATGTGACGTCCTTCCAGAATGATCTTGGA
                      ******** ************************** ** *********************  60

PIK3CB_TROSSULUS      CAACTGGACATTGACTTCCTGTTACCTAATGGTATATGTGTACCATTACAGGTAGACATA
PIK3CB_MTRBTN2        CAGCTGGACATTGACTTCCTGTTACCTAATGGTATATGTGTACCATTACAGGTAGACATA
PIK3CB_EDULIS         CAGCTGGACATTGACTTCCTCTTACCTAATGGTATATGTGTACCGTTACAGGTAGACATA
                      ** ***************** *********************** ***************  120

PIK3CB_TROSSULUS      GATAGACCACTTGATCAAATAAAACAGGAACTATGGAAACAAGCAGCAAATTACCCACTG
PIK3CB_MTRBTN2        GATAGACCACTTGATCAAATAAAACAGGAACTATGGAAACAAGCAGCAAATTACCCACTG
PIK3CB_EDULIS         GATAGACCACTTGATCAAATAAAACAGGAACTATGGAAACAAGCAGCAAATTACCCACTG
                      ************************************************************  180

PIK3CB_TROSSULUS      TTCCAGCAGCTGGCAACATTTGACAAATATGGCTTTCTTTACATCAGTGGTGAAGGACAA
PIK3CB_MTRBTN2        TTCCAACAGCTGGCAACATTTGACAAATATGGCTTTCTTTACATCAGTGGTGAAGGACAA
PIK3CB_EDULIS         TTCCAGCAGCTGGCAACATTTGACAAATATGGCTTTCTTTACATCAGTGGTGAAGGACAA
                      ***** ******************************************************  240

PIK3CB_TROSSULUS      AAGGAGGATGTTATGGATGAAAGTGTTGGAATAGCCGAGTTACGCTTGGCTGCCTCTTTC
PIK3CB_MTRBTN2        AAGGAGGATGTTATGGATGAAAGTGTTGGAATAGCCGAGTTACGCTTGGCTGCCTCTTTC
PIK3CB_EDULIS         AAGGAGGATGTTATGGATGAAAGTGTTGGGATATCCGAGTTACGCTTGGCTGCATCTTTC
                      ***************************** *** ******************* ******  300

PIK3CB_TROSSULUS      CTTAAAGTAGTAGAGAAACCTGCTGATGACAAACGTAAAACAATTGACAGACAGATTAAC
PIK3CB_MTRBTN2        CTTAAAGTAGTTGAGAAACCTGCTGATGACAAACGTAAAACAATTGACAGACAGATTAAC
PIK3CB_EDULIS         CTTAAAGTAGTAGAGAAACCTGCTGATGACAAACGTAAAACAATTGACAGACAGATTAAC
                      ************************************************************  360

PIK3CB_TROSSULUS      AGTTTGATTGGCATGAAAGGTTCTAACTACATTCAAGCTCAGCATTCAGTGAGTAAGGCA
PIK3CB_MTRBTN2        AGTTTGATTGGCATGAAAGGTTCTAACTACATTCAAGCTCAGCATTCAGTGAGTAAGGCA
PIK3CB_EDULIS         AGTTTGATTGGCATGAAAGGTTCTAACTACATTCAAGCTCAGCATTCAGTGAGTAAGGCA
                      ************************************************************  420

PIK3CB_TROSSULUS      GAGATATCAGAGTTCAGACAGAAGATGACCACTTATTGTAATAATGATGTACTGAAACAA
PIK3CB_MTRBTN2        GAGATATCAGAGTTCAGACAGAAGATGACCACTTATTGTAATAATGATGTACTGAAACAA
PIK3CB_EDULIS         GAGATATCAGAGTTTAGACAGAAGATGACCACTTATTGTAACAATGATGTACTGAAGCAA
                      ************** ************************** ************** ***  480

PIK3CB_TROSSULUS      GCCAATCAGGATCCAACAGGTTTGGTTTATTTTGAGTACAAGCATCCCTGTCGACTTAGA
PIK3CB_MTRBTN2        GCCAATCAGGATCCAACAGGTTTGGTTTATTTTGAGTACAAGCATCCCTGTCGACTTAGA
PIK3CB_EDULIS         GCCAATCAGGATCCAACAGGTTTGGTTTATTTTGAGTACAAACATCCCTGTCGACTTAGA
                      ***************************************** ******************  540

PIK3CB_TROSSULUS      AGTACCAGTGAGCTGCCTCCAGATGTAGAGAACATGGTTATAGATAATATTGTACTGACT
PIK3CB_MTRBTN2        AGTACCAGTGAGCTGCCTCCAGATGTAGAGAACATGGTTATAGATAATATTGTACTGACT
PIK3CB_EDULIS         AGTACCAGTGAGCTACCACCAGATGTAGAGAACATGGTTATAGATTATATTGTACTTACT
                      ************** ** *************************** ********** ***  600

PIK3CB_TROSSULUS      GTGATTGTACAGGAAACTGGGACTTCATACAAATTGCGAGTTAAGAAAGTGTCCACACCA
PIK3CB_MTRBTN2        GTGATTGTACAGGAAACAGGGACTTCATACAAATTGCGAGTTAAGAAAGTGTCCACACCA
PIK3CB_EDULIS         GTGATTGTACAGGAAACTGGGACTTCATACAAATTGCGAGTTAAGAAATTGTCCACACCA
                      ***************** ****************************** ***********  660

PIK3CB_TROSSULUS      GATGAGTTAGTGAAAATGACTGTTGATAAATGGTCAAGCAAATCAGGACAGAAGACTATA
PIK3CB_MTRBTN2        GATGAGTTAGTGAAAATGACTGTTGATAAATGGTCAAGCAAATCAGGACAGAAGACTATA
PIK3CB_EDULIS         GACGAGTTAGTGAAAATGACTGTAGATAAATGGTCAAGCAAATCAGGACAGAAGACTATA
                      ** ******************** ************************************  720

PIK3CB_TROSSULUS      AACTGCAATGATTATGTTCTTCAAGTTCTTGGAATGAATGATTTTCTTTATGGTGATAAT
PIK3CB_MTRBTN2        AACTGCAATGATTATGTTCTTCAAGTTCTTGGAATGAATGATTTTCTTTATGGTGATAAT
PIK3CB_EDULIS         AACTGCAATGATTATGTTCTTCAAGTTCTTGGAATGAATGATTTTCTTTATGGTGATAAT
                      ************************************************************  780

PIK3CB_TROSSULUS      CCTCTGGTTATGTTTAAGTATGTATATTCCTGTATTGTGAAAAACACAGCACCAGAATTT
PIK3CB_MTRBTN2        CCTCTGGTTATGTTTAAGTATGTATATTCCTGTATTGTGAAAAACACAGCACCAGAATTT
PIK3CB_EDULIS         CCTCTGGTTATGTTTAAGTATGTTTATTCCTGTATTGTGAAAAACACAGCGCCAGAATTT
                      *********************** ************************** *********  840

PIK3CB_TROSSULUS      TGGCTGAAGAATAAATCATCTGTCTTAGAAAAACAAGAGGTGTCACAACCACACAAATCT
PIK3CB_MTRBTN2        TGGCTGAAGAATAAATCATCTGTCTTAGAAAAACAAGAGGTGTCACAACCACACAAATCT
PIK3CB_EDULIS         TGGCTGAAGAATAAATCATCTGTCTTAGAAAAACAAGAGGTCTCACAACCACACAAATCT
                      ***************************************** ******************  900


PIK3CB_TROSSULUS      TCCAGAACGTCTTTCCAGGAAGAAAGAAGAAATACTGTGCGGTCAGAGAAGAGAAGTGGA
PIK3CB_MTRBTN2        TCCAGAACGTCTTTCCAGGAAGAAAGAAGAAATACTGTGCGGTCAGAGAAGAGAAGTGGA
PIK3CB_EDULIS         TCCAGAACATCTTTCCAGGAAGAAAGAAGAAATACTGTACGGTCAGAGAAGAGAAGTGGA
                      ******** ***************************** *********************  960

PIK3CB_TROSSULUS      CACTGTGTGTGGGACATACACAATAACTTTTATATAACCTTACACACAGCATGCAATGTT
PIK3CB_MTRBTN2        CACTGTGTGTGGGACATACACAATAACTTTTATATAACCTTACACACAGCATGCAATGTT
PIK3CB_EDULIS         CACTGTGTGTGGGACATACACCAGAACTTTTATATAACCTTACACACAGCATGCAATGTT
                      ********************* * ************************************  1020

PIK3CB_TROSSULUS      CAAGTTCCAGAAAACTGCAGAAATCTACAGGTACGTTTGTGTGTAGGTGTGTTCCATGGA
PIK3CB_MTRBTN2        CAAGTTCCAGAAAACTGCAGAAATCTACAGGTACGTTTGTGTGTAGGTGTGTTCCATGGA
PIK3CB_EDULIS         CAAGTTCCAGAAAACTGCAGAAATCTA---GTACGTTTGTGTGTAGGTGTGTTCCATGGA
                      ***************************   ******************************  1080

PIK3CB_TROSSULUS      CCTGATCCTTTGTGTAGTATTCAGGAAACCAAAGAAGTCTTTGTCTCACCAGATGGACTA
PIK3CB_MTRBTN2        CCTGATCCTTTGTGTAGTATTCAGGAAACCAAAGAAGTCTTTGTCTCACCAGATGGACTA
PIK3CB_EDULIS         CCTGATCCTTTGTGTAGTATACAGGAAACCAAAGAAGTCTTCGTCTCACAAGATGGACTG
                      ******************** ******************** ******* *********   1140

PIK3CB_TROSSULUS      TGTAGCTTTAATGACTGTATAAACTTTGATATTAAAGTACAGGATCTTCCTCAAATGAGC
PIK3CB_MTRBTN2        TGTAGCTTTAATGACTGTATAAACTTTGATATTAAAGTACAGGATCTTCCTCAAATGAGC
PIK3CB_EDULIS         TGTAGCTTTAATGATTGTATAAACTTTGATATTAAAGTACAGGATCTTCCTCAAATGAGC
                      ************** *********************************************  1200

PIK3CB_TROSSULUS      AGGTTATGTTTTGGACTCCATTGCAAGAAAGGAAAGGAACCATTATTATTAGCTTGGGCT
PIK3CB_MTRBTN2        AGGTTATGTTTTGGACTCCATTGCAAGAAAGGAAAGGAACCATTATTATTAGCTTGGGCT
PIK3CB_EDULIS         CGATTATGTTTTGGACTCCATTGCAAGAAAGGAAAGGAACCATTATTATTAGCTTGGGCT
                       * *********************************************************  1260

PIK3CB_TROSSULUS      AATATTCCAATTTTTGACTACAAATCAAATCTACAAAAAGGGAAACTGAAGTTACCATTA
PIK3CB_MTRBTN2        AATATTCCAATTTTTGACTACAAATCAAATCTACAAAAAGGGAAACTGAAGTTACCATTA
PIK3CB_EDULIS         AATATTCCAATTTTTGACTACAAATCAAATCTACAAAAAGGGAAACTGAAGTTACCATTA
                      ************************************************************  1320

PIK3CB_TROSSULUS      TGGCCAAGATCAGAACTTATACAACAGGAAGAATCAAACTGTTTTCCTGTTGGAACAGTA
PIK3CB_MTRBTN2        TGGCCAAGATCAGAACTTATACAACAGGAAGAATCAAACTGTTTTCCTGTTGGAACAGTA
PIK3CB_EDULIS         TGGCCAAGATCAGAACTTATACAACAGGAAGAGTCATACTGTTTCCCTGTTGGAACAGTG
                      ******************************** *** ******* **************   1380

PIK3CB_TROSSULUS      GCCACAAATCCTCAGGGAGATTGTTCTATGATAGAATTTAGTGTTCCAGATTTCACTAAG
PIK3CB_MTRBTN2        GCCACAAATCCTCAGGGAGATTGTTCTATGATAGAATTTAGTGTTCCAGATTTCACTAAG
PIK3CB_EDULIS         GCCACAAATCCTCAGGGAGATTGTTCTATGATAGAATTTAGTGTTCCAGATTTTACTAAG
                      ***************************************************** ******  1440

PIK3CB_TROSSULUS      AAAGGATCCATATATTACCCTCCTATTGAAAAGGTTTTACAATGTGCCAGTGACCATATG
PIK3CB_MTRBTN2        AAAGGATCCATATATTACCCTCCTATTGAAAAGGTTTTACAATGTGCCAGTGACCATATG
PIK3CB_EDULIS         AAAGGATCCATATATTACCCTCCAATTGAAAAGGTTTTACAATGTGCCAGTGATCATATG
                      *********************** ***************************** ******  1500

PIK3CB_TROSSULUS      GAAGAGCCAGGGAAATGTGGGTCACCAACTTGGAGACCAAACAAGAACATTGTTCAGCAG
PIK3CB_MTRBTN2        GAAGAGCCAGGGAAATGTGGGTCACCAACTTGGAGACCAAACAAGAACATTGTTCAGCAG
PIK3CB_EDULIS         GAAGAGCCAGGGAAATGTGGGTCACCAACATGGAGACCAAACAAGAACATTGTTCAGCAG
                      ***************************** ******************************  1560

PIK3CB_TROSSULUS      CTAGACACCATACTAAGAGCTAACCAGTACACTGTACTTCAGACTTTGGACGAACAGCAG
PIK3CB_MTRBTN2        CTAGACACCATACTAAGAGCTAACCAGTACACTGTACTTCAGACTTTGGACGAACAGCAG
PIK3CB_EDULIS         CTAGACAGCATACTGAGGGCTAACCAGTACACAATACTTCAGACATTGGACGAACAGCAG
                      ******* ****** ** **************  ********** ***************  1620

PIK3CB_TROSSULUS      AAACAACTTATTTGGTGGATGCGTTATGACATACGTGACAAATGCAGCGAGTTCCCTCAT
PIK3CB_MTRBTN2        AAACAACTTATTTGGTGGATGCGTTATGACATACGTGACAAATGCAGCGAGTTCCCTCAT
PIK3CB_EDULIS         AAACAACTGATTTGGTGGATGCGTTATGACATACGTGACAAATGCAGCGAGTTCCCTCAC
                      ******** **************************************************   1680

PIK3CB_TROSSULUS      GCTCTGTTGTATGTGTTATATTCAGTTAGCTGGGACAACCATATTGAAGTTTCCAAGATG
PIK3CB_MTRBTN2        GCTCTGTTGTATGTGTTATATTCAGTTAGCTGGGACAACCATATTGAAGTTTCCAAGATG
PIK3CB_EDULIS         GCTCTGTTGTATGTGTTATATTCTGTCAGCTGGGACAACCATATTGAAGTTTCCAAGATG
                      *********************** ** *********************************  1740

PIK3CB_TROSSULUS      CAGGCACTATTACAGACATGGCCTACACTAGAGGCAGATCAAGCCTTAACATTGCTAGAT
PIK3CB_MTRBTN2        CAGGCACTATTACAGACATGGCCTACACTAGAGGCAGATCAAGCCTTAACATTGCTAGAT
PIK3CB_EDULIS         CAGGCTCTGTTACAGACATGGCCTACACTAGAGGCAGATCAAGCCTTAACTTTGCTAGAT
                      ***** ** ***************************************** *********  1800

PIK3CB_TROSSULUS      TTTACATTCCCAGATAAATATGTCAGAAAGAAAGCCACTGAATGGTTAGATGAACTGCCT
PIK3CB_MTRBTN2        TTTACATTCCCAGATAAATATGTCAGAAAGAAAGCCACTGAATGGTTAGATGAACTGCCT
PIK3CB_EDULIS         TTTACATTCCCTGATAAATATGTCAGAAAGAAAGCCACTGAATGGTTAGATGAACTGCCT
                      *********** ************************************************  1860

PIK3CB_TROSSULUS      GATGAAGAATTGGCTCAGTATTTGCTACAGTTAGTACAGGCTTTAAAGTATGAGAACTAT
PIK3CB_MTRBTN2        GATGAAGAATTGGCTCAGTATTTGCTACAGTTAGTACAGGCTTTAAAGTATGAGAACTAT
PIK3CB_EDULIS         GATGAAGAATTGGCTCAGTATTTGCTACAGTTGGTACAGGCTTTGAAGTATGAGAACTAT
                      ******************************** *********** ***************  1920

PIK3CB_TROSSULUS      TTAAACTGTGATCTAGTCAAATTCCTGTTACGGAGAGCACTACAGAATCGTAATATAGGA
PIK3CB_MTRBTN2        TTAAACTGTGATCTAGTCAAATTCCTGTTACGGAGAGCACTACAGAATCGTAATATAGGA
PIK3CB_EDULIS         TTAAATTGTGATCTAGTCAAATTCCTGTTACGGAGAGCACTTCAGAATCGTAATATAGGA
                      ***** *********************************** ******************  1980

PIK3CB_TROSSULUS      CATAAGTTATTTTGGTTGCTCAAGTCAGACATGCATGAACCCTCTGTTACCGTCCAGTAT
PIK3CB_MTRBTN2        CATAAGTTATTTTGGTTGCTCAAGTCAGACATGCATGAACCCTCTGTTACCGTCCAGTAT
PIK3CB_EDULIS         CATAAGTTATTTTGGTTGCTCAAGTCAGACATGCATGAACCCTCTGTTACCGTCCAGTAT
                      ************************************************************  2040

PIK3CB_TROSSULUS      GGTTTAATCATAGAAATATATCTCAAGGCAAACCCCAGTCATATGTCCATCCTAGACAGG
PIK3CB_MTRBTN2        GGTTTAATCATAGAAATATATCTCAAGGCAAACCCCAGTCATATGTCCATCCTAGACAGG
PIK3CB_EDULIS         GGTTTAATCATAGAAATATATCTCAAGGCAAACCCCAGTCACATGTCCATCCTTGATAGG
                      ***************************************** *********** ** ***  2100

PIK3CB_TROSSULUS      CAGCAAATCATCCTACAGAAACTCAAATCTATCACAGATATACATACCAAAGCAGCAAAT
PIK3CB_MTRBTN2        CAGCAAATCATCCTACAGAAACTCAAATCTATCACAGATATACATACCAAAGCAGCAAAT
PIK3CB_EDULIS         CAGCAAATCATCCTACAGAAACTCAAATCTATCACAGATATACATACCAAAGCAGCAAAT
                      ************************************************************  2160

PIK3CB_TROSSULUS      TTGAAGAAAAAGGCTAAAGAAAAAGATGACTCTCCCTTGCAGAAGGTTATGCAGCAATCT
PIK3CB_MTRBTN2        TTGAAGAAAAAGGCTAAAGAAAAAGATGACTCTCCCTTGCAGAAGGTTATGCAGCAATCT
PIK3CB_EDULIS         TTGAAGAAAAAGGCTAAAGAAAAAGATGACTCTCCCTTGCAGAAGGTTATGCAGCAATCT
                      ************************************************************  2220

PIK3CB_TROSSULUS      GCATACAAAGAGGCGTTCTGTCAAATATACAGTCCTATCACCTTAATGTATAAATTAGAT
PIK3CB_MTRBTN2        GCATACAAAGAGGCGTTCTGTCAAATATACAGTCCTATCACCTTAATGTATAAATTAGAT
PIK3CB_EDULIS         GCATACAAAGAGGCGTTCTGTAAAATATATAGTCCTATCACCTTAATGTATAAATTAGAC
                      ********************* ******* *****************************   2280

PIK3CB_TROSSULUS      AAAATTGTTGAGAAATCCTGTAAAGTTATGGACTCTAAAAAGAAGCCATTTTGGTTGGAA
PIK3CB_MTRBTN2        AAAATTGTTGAGAAATCCTGTAAAGTTATGGACTCTAAAAAGAAGCCATTTTGGTTGGAA
PIK3CB_EDULIS         AAAATTGTTGAGAAATCCTGTAAAGTAATGGACTCTAAAAAGAAGCCGTTTTGGTTGGAA
                      ************************** ******************** ************  2340

PIK3CB_TROSSULUS      TGGACAAATGATGATGAGAAAGGACAAAACATACAGTTGATATATAAATGTGGAGATGAT
PIK3CB_MTRBTN2        TGGACAAATGATGATGAGAAAGGACAAAACATACAGTTGATATATAAATGTGGAGATGAT
PIK3CB_EDULIS         TGGACAAATGATGATGAGAAAGGACAAAACATACAGTTGATTTATAAATGTGGAGATGAT
                      ***************************************** ******************  2400

PIK3CB_TROSSULUS      TTACGACAAGATATGCTGACATTACAGATTTTAGAAGTCATGGATACTATATGGCAATCA
PIK3CB_MTRBTN2        TTACGACAAGATATGCTGACATTACAGATTTTAGAAGTCATGGATACTATATGGCAATCA
PIK3CB_EDULIS         TTACGACAAGATATGCTGACATTACAGATTTTAGAAGTCATGGATACTATATGGCAATCA
                      ************************************************************  2460

PIK3CB_TROSSULUS      CAAGGCTATGATCTCAGACTAAATCCTTATGGATGTGTTGCAACTGGATGTGAAGAGGGT
PIK3CB_MTRBTN2        CAAGGCTATGATCTCAGACTAAATCCTTATGGATGTGTTGCAACTGGATGTGAAGAGGGT
PIK3CB_EDULIS         GAAGGCTATGATCTCAGACTAAATCCTTATGGATGTGTTGCAACTGGATGTGAAGAGGGT
                       ***********************************************************  2520


PIK3CB_TROSSULUS      ATGATTGAAGTTGTACAGAAATCTTTAACACTTGCTGGAATACAAAAATGGAGAAAACTA
PIK3CB_MTRBTN2        ATGATTGAAGTTGTACAGAAATCTTTAACACTTGCTGGAATACAAAAATGGAGAAAACTA
PIK3CB_EDULIS         ATGATTGAAGTTGTACAGAAATCTTTAACACTTGCCGGAATACAAAAATGGAGAAAACTA
                      *********************************** ************************  2580

PIK3CB_TROSSULUS      GGACTGGATAAGAGATCACTTTATGATTGGTTAAAACATAAAAATCCAACTGAACATAGT
PIK3CB_MTRBTN2        GGACTGGATAAGAGATCACTTTATGATTGGTTAAAACATAAAAATCCAACTGAACATAGT
PIK3CB_EDULIS         GGACTGGATAAGAGATCACTTTATGATTGGTTAAAACATAAAAATCCAACTGAACACAGT
                      ******************************************************** ***  2640

PIK3CB_TROSSULUS      TTACAAAGAGCAGTTGAAGAATTTAAGCTGTCCTGTGCCGGATATGCTGTGGCCACCTAC
PIK3CB_MTRBTN2        TTACAAAGAGCAGTTGAAGAATTTAAGCTGTCCTGTGCCGGATATGCTGTGGCCACCTAC
PIK3CB_EDULIS         TTACAGAGAGCAGTTGAAGAATTTAAGCTGTCCTGTGCCGGATATGCTGTGGCCACCTAC
                      ***** ******************************************************  2700

PIK3CB_TROSSULUS      ATCCTGGGAGTTGGGGACAGACATAACGACAATATAATGATGAAAGAAAGTGGACAGTTG
PIK3CB_MTRBTN2        ATCCTGGGAGTTGGGGACAGACATAACGACAATATAATGATGAAAGAAAGTGGACAGTTG
PIK3CB_EDULIS         ATCCTCGGAGTTGGTGACAGACATAACGACAATATAATGATGAAAGAAACTGGACAGTTG
                      ***** ******** ********************************** **********  2760

PIK3CB_TROSSULUS      TTTCATATAGACTTTGGACATTTTCTTGGAAATAAAAAAACAAAGTTTAACATTAACAGA
PIK3CB_MTRBTN2        TTTCATATAGACTTTGGACATTTTCTTGGAAATAAAAAAACAAAGTTTAACATTAACAGA
PIK3CB_EDULIS         TTTCATATAGACTTTGGACATTTTCTGGGAAATAAAAAAACAAAGTTTAACATTAACAGA
                      ************************** *********************************  2820

PIK3CB_TROSSULUS      GAACGAGTGCCATTTATTCTTACCAGCCATTTTGAGTATATCATAAAAGATGGGGATCAA
PIK3CB_MTRBTN2        GAACGAGTGCCATTTATTCTTACCAGCCATTTTGAGTATATCATAAAAGATGGGGATCAA
PIK3CB_EDULIS         GAACGAGTGCCATTTATTCTAACTAGCCATTTTGAGTATATCATAAAAGACGGTGAAAAA
                      ******************** ** ************************** ** **  **  2880

PIK3CB_TROSSULUS      AAGCCACAAAATTTCACAGATTTTAAAGACATTTGTGAGAGGGCTTACTTAATCATTCGT
PIK3CB_MTRBTN2        AAGCCACAAAATTTCACAGATTTTAAAGACATTTGTGAGAGGGCTTACTTAATCATTCGT
PIK3CB_EDULIS         AAGCCACAAAATTTCACACATTTTAAAGAAATTTGTGAGAGAGCTTACTTAATCATTCGT
                      ****************** ********** *********** ******************  2940

PIK3CB_TROSSULUS      AGCAAAGCACACCTACTGATTCAGTTGTTTACCATGATGCTGTCATCAGGAATACCTCAG
PIK3CB_MTRBTN2        AGCAAAGCACACCTACTGATTCAGTTGTTTACCATGATGCTGTCATCAGGAATACCTCAG
PIK3CB_EDULIS         AGCAGAGCACACCTACTGATTCAGTTGTTTATGATGATGCTCTCATCAGGAATACCTCAG
                      **** **************************  ******** ******************  3000

PIK3CB_TROSSULUS      CTTAATAATGTCTCAGATATTGATTACATTAAAGACGTTCTTGCCTTGAACTCAACTGAA
PIK3CB_MTRBTN2        CTTAATAATGTCTCAGATATTGATTACATTAAAGACGTTCTTGCCTTGAACTCAACTGAA
PIK3CB_EDULIS         CTCAATAATGTCTCAGATATTGATTACATTAAAGAAGTTCTTGCCTTGAACTCGACACAA
                      ** ******************************** ***************** **  **  3060

PIK3CB_TROSSULUS      GAAGTGGCTCTAGACAAATTCAGGAAAAAATTCAAAGAAGCTCAAGACAGCAGTTGGTCA
PIK3CB_MTRBTN2        GAAGTGGCTCTAGACAAATTCAGGAAAAAATTCAAAGAAGCTCAAGACAGCAGTTGGTCA
PIK3CB_EDULIS         GAAGAGGCTCTAGAAAAATTCAGGAAAAAATTCAAAGAAGCTCAAGAAAGCAGTTGGTCA
                      **** ********* ******************************** ************  3120

PIK3CB_TROSSULUS      ACTACAGTAAATTGGTGGTTCCATATGAGAGTCCATTGA
PIK3CB_MTRBTN2        ACTACAGTAAATTGGTGGTTCCATATGAGAGTCCATTGA
PIK3CB_EDULIS         ACTACAGTGAATTGGTGGTTTCATATGAGAGTCCATTGA
                      ******** *********** ******************  3159
```

**ALIGNMENTS PTEN-GENE_NUCLEOTIDE SEQUENCES**

```
PTEN_TROSSULUS      ATGGATTTTTTAAAAGGTCTTGTCAGTAAAAACAAGAGACGACACAAAGTGGACGGATTT
PTEN_CANCER         ATGGATTTTTTAAAAGGTCTTGTCAGTAAAAACAAGAGACGACACAAAGTGGACGGATTT
PTEN_EDULIS         ATGGATTTTTTAAAAGGTCTTGTCAGTAAAAACAAGAGACGGCACAAAGTCGACGGATTC
                    ***************************************** ******** ********   60

PTEN_TROSSULUS      GATCTAGATCTGACATACATCTATCCTAACATTATAGCCATGGGCTTTCCTGCAGAAAAG
PTEN_CANCER         GATCTAGATCTGACATACATCTATCCTAACATTATAGCCATGGGCTTTCCTGCAGAAAAG
PTEN_EDULIS         GATCTAAATCTGACATACATCTATCCTAACATTATAGCCATGGGCTTTCCTGCAGAAAAA
                    ****** ****************************************************   120

PTEN_TROSSULUS      CTGGAGGGAGTGTACAGAAACCATATTGACGATGTTATTGAGTTTCTAGACACAAAGCAT
PTEN_CANCER         CTGGAGGGAGTTTACAGAAACCATATTGACGATGTTATTGAGTTTCTAGACACAAAGCAT
PTEN_EDULIS         CTGGAGGGAGTTTACAGAAACCATATTGATGATGTTATTGAATTTCTAGACACTAAGCAT
                    *********** ***************** *********** *********** ******  180

PTEN_TROSSULUS      AAAGACCACTACAAAGTATATAATTTATGTACAGAAAGGTCCTACGACCCTGACAGATTC
PTEN_CANCER         AAAGACCACTACAAAGTATATAATTTATGTACAGAAAGGTCCTACGACCCTGACAGATTC
PTEN_EDULIS         AAAGACCACTACAAAGTATATAATCTATGTACAGAAAGGTCCTACGACCCTGACAGATTC
                    ************************ ***********************************  240

PTEN_TROSSULUS      CATGGGAGAGTTGTAGCCTATCCATTTGAGGACCACAATCCACCGAGATTAGAGTTAATA
PTEN_CANCER         CATGGGAGAGTTGTAGCCTATCCATTTGAGGACCACAATCCACCGAGATTAGAGTTAATA
PTEN_EDULIS         CATGGGAGAGTTGTAGCCTATCCATTTGAGGACCACAATCCACCGAGATTAGAGTTAATA
                    ************************************************************  300

PTEN_TROSSULUS      AAACCATTCTGTGAGGACCTGGATGAGTGGTTGAAGAAAAGCAATGAGAACATAGCAGCC
PTEN_CANCER         AAACCATTCTGTGAGGACCTGGATGAGTGGTTGAAGAAAAGCAATGAGAACATAGCAGCC
PTEN_EDULIS         AAACCATTCTGTGAGGACCTGGATGAGTGGTTGAAGAAAAGCGATGAAAACATAGCAGCC
                    ****************************************** **** ************  360

PTEN_TROSSULUS      ATTCATTGTAAGGCAGGAAAGGGAAGAACTGGTGTTATGATATGTGCCTATATGTTACAC
PTEN_CANCER         ATTCATTGTAAGGCAGGAAAGGGAAGAACTGGTGTTATGATATGTGCCTATATGTTACAC
PTEN_EDULIS         ATTCATTGTAAGGCAGGAAAGGGAAGAACTGGTGTTATGATATGTGCCTATATGTTACAC
                    ************************************************************  420

PTEN_TROSSULUS      AGAAACAAATTTGATAACTCTAAAGAGGCACTGCGGTTCTATGGCCAGGCACGGACACAG
PTEN_CANCER         AGAAACAAATTTGATAACTCTAAAGAGGCACTGCGGTTCTATGGCCAGGCACGGACACAG
PTEN_EDULIS         AGGAACAAATTTGATAACACTAAAGAAGCACTGCGGTTCTATGGCCAGGCACGGACACAG
                    ** *************** ******* *********************************  480

PTEN_TROSSULUS      GATGAAAAGGGGGTAACCATTCCTAGCCAGAGACGATATGTAGAATATTATGAATATTTA
PTEN_CANCER         GATGAAAAGGGGGTAACCATTCCTAGCCAGAGACGATATGTAGAATATTATGAATATTTA
PTEN_EDULIS         GATGAAAAGGGGGTAACCATTCCTAGCCAGAGACGATATGTAGAATATTATGAATATTTA
                    ************************************************************  540

PTEN_TROSSULUS      ATTAGAAACAAATTAAACTACAAGCCAGTAGCCTTGTTGTTAAAAGGCATAGAATTTATC
PTEN_CANCER         ATTAGAAACAAATTAAACTACAAGCCAGTAGCCTTGTTGTTAAAAGGCATAGAATTTATC
PTEN_EDULIS         ATTAGAAACAAATTAAACTACAAGCCAGTAGCATTGTTGTTAAAAGGCATAGAATTTATA
                    ******************************** **************************   600

PTEN_TROSSULUS      ACAGTTCCCATGCACAATGGGAGTGGATGTTCTCCATTTTTTGAAGTATATCAACTTAAG
PTEN_CANCER         ACAGTTCCCATGCACAATGGGAGTGGATGTTCTCCATTTTTTGAAGTATATCAACTTAAG
PTEN_EDULIS         ACAGTTCCCATGCACAATGGGAGTGGATGTTCTCCATTTTTTGAAGTATATCAACTTAAG
                    ************************************************************  660

PTEN_TROSSULUS      GTGCGAGTATACACATCAAAAGTTTATGAAGACATCAAAAAGGGCCAAACCTCGTTTTAC
PTEN_CANCER         GTGCGAGTATACACATCAAAAGTTTATGAAGACATCAAAAAGGGCCAAACCTCGTTTTAC
PTEN_EDULIS         GTGCGAGTATACACATCAAAAGTTTACGAAGACATCAAAAAGGGCCAAACATCGTTTTAC
                    ************************** *********************** *********  720

PTEN_TROSSULUS      ATGCCCATAGAACAGTCGGTACCGCTGTGTGGAGATATCAAAGTGGTATTTTATAATAAA
PTEN_CANCER         ATGCCCATAGAACAGTCGGTACCGCTGTGTGGAGATATCAAAGTGGTATTTTATAATAAA
PTEN_EDULIS         ATGCCCATAGAACAGTCTGTACCGCTGTGTGGAGATATCAAAGTTGTATTTTATAATAAA
                    ***************** ************************** ***************  780

PTEN_TROSSULUS      CCAAGAATGAAAAAAAAGGATAAAATGTTTCAATTTTGGTTAAACACATTTTTTGTGGAA
PTEN_CANCER         CCAAGAATGAAAAAAAAGGATAAAATGTTTCAATTTTGGTTAAACACATTTTTTGTGGAA
PTEN_EDULIS         CCAAGAATGAAAAAGAAGGATAAAATGTTTCAATTTTGGTTAAACACATTTTTTGTGGAA
                    ************** *********************************************  840

PTEN_TROSSULUS      GCAGACGACAAGCCAAGAGAGAATGGGCGGAAGTCTTGGGCAGGAGAGGTTCCAGGCTCC
PTEN_CANCER         GCAGACGACAAGCCAAGAGAGAATGGCCGGAAGTCTTGGGCCGGAGAGGTTCCAGGGTCC
PTEN_EDULIS         GCAGACGACAAGCCAAAAGAAAATGGACGGAAATCTTGGGCAGGAGAGGTGCCAGGGTCC
                    **************** *** ***** ***** ******** ******** ***** ***  900

PTEN_TROSSULUS      AGTAATTATTACACAGTGACTATTCCTAAGTGTGAACTAGACAAAGCTAATAAAGATAAG
PTEN_CANCER         AGTAATTATTACACAGTGACTATTCCTAAGTGTGAACTAGACAAAGCTAATAAAGATAAG
PTEN_EDULIS         AGTAATTATTACACAGTGACTATTCCTAAGTGTGAACTAGACAAAGCTAATAAAGATAAG
                    ************************************************************  960

PTEN_TROSSULUS      GCTCACAAGTTATTTAGTCCTAATTTCCAGGTCAAACTTCATTTCACCAATCCTGAAAAT
PTEN_CANCER         GCTCACAAGTTATTTAGTCCTAATTTCCAGGTCAAACTTCATTTCACCAATCCTGAAAAT
PTEN_EDULIS         GCTCACAAGTTATTTAGTCCTAATTTCCAGGTCAAACTTCATTTCACCAATCCTGAAAAT
                    ************************************************************  1020

PTEN_TROSSULUS      TGTGGATTATATAACTCCTCATTAAAAGTCCCAGATCAGCAAGGATTACGACACCGATCA
PTEN_CANCER         TGTGGATTATATAACTCCTCATTAAAAGTCCCAGATCAGCAAGGATTACGACACCGATCA
PTEN_EDULIS         TGTGGATTATATAACTCCTCATTAAAAGTCCCCGATCAGCAAGGATTACGACACCGATCA
                    ******************************** ***************************  1080


PTEN_TROSSULUS      AAAACATTAGAACAAATATGTGATAAAAACTCCACTACAAATCACTCGGACCATAGTGTG
PTEN_CANCER         AAAACATTAGAACAAATATGTGATAAAAACTCCACTACAAATCACTCGGACCATAGTGTG
PTEN_EDULIS         AAAACATTAGAACAAATATGTGATAAAAACTCCACTACAAATCACTCGGACCATAGTGTG
                    ************************************************************  1140

PTEN_TROSSULUS      GATGGGATGCCACATTCTATAACGGCACTGACACTTAACTCTCGACGAGAAAATCAGTCG
PTEN_CANCER         GATGGGATGCCACATTCTATAACGGCACTGACACTTAACTCTCGACGAGAAAATCAGTCG
PTEN_EDULIS         GACGGGATGCCACATTCTATAACGGCACTGACACTTAACTCTCGACGAGAAAATCAGTCG
                    ** *********************************************************  1200

PTEN_TROSSULUS      CCTGTGATAACACGGACACGATCACCTAGTCACATGACTCATGAACGACCAAAATTCAAT
PTEN_CANCER         CCTGTGATAACACGGACACGATCACCTAGTCACATGACTCATGAACGACCAAAATTCAAT
PTEN_EDULIS         CCTGTGATAACACGGACACGATCACCTAGTCACATGACTCATGAACGACCAAAATTCATG
                    **********************************************************    1260

PTEN_TROSSULUS      TTGCCTTCTAATGGGGAGGGCAGTAGTGACCAATTAAGTAGTGAGGCCAATAGTGACACA
PTEN_CANCER         TTGCCTTCTAATGGGGAGGGCAGTAGTGACCAATTAAGTAGTGAGGCCAATAGTGACACA
PTEN_EDULIS         TTGCCGTCTAACGGGGAGGGCAGTAGTGACCAATTAAGTAGTGAGGCAAATAGTGACACA
                    ***** ***** *********************************** ************  1320

PTEN_TROSSULUS      GACTTTCCCGAGGAGGATTTGTCAGACACGGACGATGAAGAAGAATGGAACGATCTAGAA
PTEN_CANCER         GACTTTCCCGAGGAGGATTTGTCAGACACGGACGATGAAGAAGAATGGAACGATCTAGAA
PTEN_EDULIS         GACTTTCCCGAGGAGGATTTGTCAGACACGGACGATGAAGAAGAATGGAATGATCTAGAA
                    ************************************************** *********  1380

PTEN_TROSSULUS      ACGACTGCTGTTTGA
PTEN_CANCER         ACGACTGCTGTTTGA
PTEN_EDULIS         ACGACTGCTGTTTGA
                    *************** 1395
```

**ALIGNMENTS AKT1-GENE_NUCLEOTIDE SEQUENCES**

```
AKT_EDULIS               ATGAACCCGTCTGGAAGTAGCATTCCTGTGGTGAAGGAAGGATATTTAATGAAAAGAGGA
AKT_TROSSULUS            ATGAACCCGTCTGGAAGTAGCATTCCTGTGGTGAAGGAAGGATATTTAATGAAAAGAGGA
AKT_CANCER               ATGAACCCGTCTGGAAGTAGCATTCCTGTGGTGAAGGAAGGATATTTAATGAAAAGAGGA
                         ************************************************************  60

AKT_EDULIS               GAACATATAAAAAATTGGAGAAAAAGATATTTTATTTTAAGAAAAGATGGTTCCTTTCTA
AKT_TROSSULUS            GAACATATAAAAAATTGGAGAAAAAGATATTTTATTTTAAGAAAAGATGGTTCCTTTCTA
AKT_CANCER               GAACATATAAAAAATTGGAGAAAAAGATATTTTATTTTAAGAAAAGATGGTTCCTTTCTA
                         ************************************************************  120

AKT_EDULIS               GGCTTCAGGGCAAAGCCAGAACAAGGATTTAATGACCCACTTAATGATTTTACTGTTAGA
AKT_TROSSULUS            GGCTTCAGGGCAAAGCCTGAACAAGGATTTAATGACCCACTTAATGATTTTACTGTTAGA
AKT_CANCER               GGCTTCAGGGCAAAGCCTGAACAAGGATTTAATGACCCACTTAATGATTTTACTGTTAGA
                         ***************** ******************************************  180

AKT_EDULIS               GATTGTCAGTTAATGAGACAAGAACGTCCAAAGGCCAATACGTTTATAATCCGAGGATTA
AKT_TROSSULUS            GATTGTCAGTTAATGAGACAAGAACGTCCAAAGGCCAATACATTTATAATCAGAGGATTA
AKT_CANCER               GATTGTCAGTTAATGAGACAAGAACGTCCAAAGGCCAATACGTTTATAATCAGAGGATTA
                         ***************************************** ********* ********  240

AKT_EDULIS               CAATGGACAACAGTAGTGGAGAGGATGTTCTGTGTAGAATCTCAGGAGGAGAGAGAAGAG
AKT_TROSSULUS            CAATGGACAACAGTGGTAGAGAGGATGTTCTGTGTAGAATCTCAGGAGGAGAGAGAAGAG
AKT_CANCER               CAATGGACAACAGTGGTAGAGAGGATGTTCTGTGTAGAATCTCAGGAGGAGAGAGAAGAG
                         ************** ** ******************************************  300

AKT_EDULIS               TGGATAGCAGCTATCAATTCTATCTCGGATGAGTTGAAAAACTCTGATGAAGGAGCAGAG
AKT_TROSSULUS            TGGATAGCAGCTATCAATTCTATCTCAGATGAGTTGAAAAACTCGGATGAAGGAGCAGAG
AKT_CANCER               TGGATAGCAGCTATCAATTCTATCTCAGATGAGTTGAAAAACTCGGATGAAGGAGCAGAG
                         ************************** ***************** ***************  360

AKT_EDULIS               GGACCATCACTTTTTGACAAATCAAAGAAAAAAGTGTCATTAGATGACTTTGAGTTTTTG
AKT_TROSSULUS            GGACCATCACTTTTTGACAAATCAAAGAAAAAAGTGTCATTAGATGACTTTGAGTTTTTG
AKT_CANCER               GGACCATCACTTTTTGACAAATCAAAGAAAAAAGTGTCATTAGATGACTTTGAGTTTTTG
                         ************************************************************  420

AKT_EDULIS               AAAGTACTAGGAAAAGGAACATTTGGAAAAGTCATCCTTTGTAAAGAGAAGGTGACAGGT
AKT_TROSSULUS            AAAGTACTAGGAAAAGGAACATTTGGAAAAGTCATCCTTTGTAAAGAGAAGGTGACAGGT
AKT_CANCER               AAAGTACTAGGAAAAGGAACATTTGGAAAAGTCATCCTTTGTAAAGAGAAGTTGACAGGT
                         *************************************************** ********  480


AKT_EDULIS               CATCTATTAGCAATCAAAATATTAAAGAAGCAAGTAATCATTCAAAAGGATGAAGTTGCA
AKT_TROSSULUS            CATCTATTAGCAATCAAAATATTAAAGAAGCAAGTAATCATTCAAAAGGATGAAGTTGCA
AKT_CANCER               CATCTATTAGCAATCAAAATATTAAAGAAGCAAGTAATCATTCAAAAGGATGAAGTTGCA
                         ************************************************************  540

AKT_EDULIS               CATACTTTGACTGAAAACAGAGTATTACAGACTACAAAACATCCATTCTTAACACAATTG
AKT_TROSSULUS            CATACTTTGACTGAAAACAGAGTATTACAGACAACAAAACATCCATTCTTAACACAATTG
AKT_CANCER               CATACTTTGACTGAAAACAGAGTATTACAGACAACAAAACATCCATTCTTAACACAATTG
                         ******************************** ***************************  600

AKT_EDULIS               AAGTACTCATTCCAGACGACAGATAGGTTATGTTTTGTTATGGAGTATGTAAATGGTGGT
AKT_TROSSULUS            AAGTACTCATTCCAGACGACAGATAGGTTATGTTTTGTTATGGAGTATGTAAATGGTGGT
AKT_MTRBTN2              AAGTACTCATTCCAGACGACAGATAGGTTATGTTTTGTTATGGAGTATGTAAATGGTGGT
                         ************************************************************  660

AKT_EDULIS               GAACTTTTCTTTCATTTATCACGAGAACGTGTGTTTTCAGAAGAAAGAACAAAATTTTAT
AKT_TROSSULUS            GAACTTTTCTTTCATTTATCACGAGAACGTGTGTTTTCAGAAGAAAGAACAAAATTTTAT
AKT_CANCER               GAACTTTTCTTTCATTTATCACGAGAACGTGTGTTTTCAGAAGAAAGAACAAAATTTTAT
                         ************************************************************  720

AKT_EDULIS               GGTGCAGAAATAATTTCAGCATTAGGATATTTACATGAAAATAATATTGTTTATAGAGAT
AKT_TROSSULUS            GGTGCAGAAATAATTTCAGCATTAGGATATTTACATGAAAATAATATTGTTTATAGAGAT
AKT_CANCER               GGTGCAGAAATAATTTCAGCATTAGGATATTTACATGAAAATAATATTGTTTATAGAGAT
                         ************************************************************  780

AKT_EDULIS               CTGAAGTTAGAAAATTTGTTGTTAGATAAAGATGGACATATCAAAATAGCTGATTTTGGT
AKT_TROSSULUS            CTGAAGTTAGAAAATTTGTTGTTAGATAAAGATGGACATATCAAAATAGCTGATTTTGGT
AKT_CANCER               CTGAAGTTAGAAAATTTGTTGTTAGATAAAGATGGACATATCAAAATAGCTGATTTTGGT
                         ************************************************************  840

AKT_EDULIS               CTTTGTAAAGAAGAAATGTTTTATGGAGCTAGTACAAAAACATTTTGTGGAACTCCAGAG
AKT_TROSSULUS            CTTTGTAAAGAAGAAATGTTTTATGGAGCTAGTACAAAAACATTTTGTGGAACTCCAGAG
AKT_CANCER               CTTTGTAAAGAAGAAATGTTTTATGGAGCTAGTACAAAAACATTTTGTGGAACTCCAGAG
                         *********************************** ************** *********  900

AKT_EDULIS               TATTTGGCACCTGAAGTTTTAGAAGACAATGATTATGGTAGAGCAGTAGATTGGTGGGGC
AKT_TROSSULUS            TATTTGGCACCTGAAGTTTTAGAAGACAATGATTATGGTCGAGCAGTAGATTGGTGGGGC
AKT_CANCER               TATTTGGCACCTGAAGTTTTAGAAGACAATGATTATGGTCGAGCTGTAGATTGGTGGGGC
                         *************************************** **** ***************  960

AKT_EDULIS               ACAGGTGTAGTTATGTATGAAATGATGTGTGGACGTTTACCTTTTTACAATCGTGATCAT
AKT_TROSSULUS            ACAGGTGTAGTTATGTATGAAATGATGTGTGGACGTTTACCTTTTTACAATCGTGATCAT
AKT_CANCER               ACAGGTGTAGTTATGTATGAAATGATGTGTGGACGTTTACCTTTTTACAATCGTGATCAT
                         ************************************************************ 1020

AKT_EDULIS               GACGTTTTATTTGAACTTATATTACTACAATCAGTGCGGTTTCCCAGGACATTGTCAGAA
AKT_TROSSULUS            GATGTTTTATTTGAACTTATATTACTACAATCAGTGCGGTTTCCCAGGACATTGTCAGAA
AKT_CANCER               GATGTTTTATTTGAACTTATATTACTACAATCAGTGCGGTTTCCCAGGACATTGTCAGAA
                         ** ********************************************************* 1080

AKT_EDULIS               GATGCCAAGAGTCTTCTGGAAGGGTTTTTAAAGAAAAATCCACACGAAAGACTTGGAGGA
AKT_TROSSULUS            GATGCCAAGAGTCTTCTGGAAGGGTTTTTGAAGAAAAATCCACATGAAAGACTTGGAGGT
AKT_CANCER               GATGCCAAGAGTCTTCTGGAAGGGTTTTTGAAGAAAAATCCACATGAAAGACTTGGAGGT
                         ***************************** ************** **************  1140

AKT_EDULIS               AGTGAACAGGATGTCAAAGAAATTATAGCACATCCATTCTTTAAAACAGTCAACTGGCAG
AKT_TROSSULUS            AGTGAGCAGGATGTCAAAGAAATAATAGCACATCCATTCTTTAAAACAGTCAACTGGCAG
AKT_CANCER               AGTGAGCAGGATGTCAAAGAAATAATAGCACATCCATTCTTTAAAACAGTCAACTGGCAG
                         ***** ***************** ************************************ 1200

AKT_EDULIS               GACTTAATTGAAAAGAAGATAACACCACCTTGGAAGCCTGACGTCAAAAATGACTATGAC
AKT_TROSSULUS            GACTTAATTGAGAAGAAGATAACACCACCTTGGAAGCCTGACGTCAAAAATGACTATGAT
AKT_CANCER               GACTTAATTGAGAAGAAGATAACACCACCTTGGAAGCCTGACGTCAAAAATGACTATGAT
                         *********** ***********************************************  1260

AKT_EDULIS               ACAAAATACATACCTGATGAGTTTGCCCAGGTTCCTCTGTCCTTGACACCTAACTCTAAA
AKT_TROSSULUS            ACAAAATACATACCTGATGAGTTTGCCCAGGTTCCTCTGTCCTTGACACCTAACTCTAAA
AKT_CANCER               ACAAAATACATACCTGATGAGTTTGCCCAGGTTCCTCTGTCCTTGACACCTAACTCTAAA
                         ************************************************************ 1320

AKT_EDULIS               GAAAGTCACGTAGAGCTCAGTTCCATAGCAGAGGAAAGTGAACTTCCTTACTTTGAACAA
AKT_TROSSULUS            GAAAGTCATGTAGAGCTCAGTTCCATAGCAGAGGAAAGTGAACTTCCTTACTTTGAACAA
AKT_CANCER               GAAAGTCATGTAGAGCTCAGTTCCATAGCAGAGGAAAGTGAACTTCCTTACTTTGAACAA
                         ******** *************************************************** 1380


AKT_EDULIS               TTCTCATACCACGGATCAAGAGCTGGTATGGGAGAAAGCTACCTCCAATCACATGTAAAT
AKT_TROSSULUS            TTCTCATACCACGGATCAAGAGCTGGTATGGGAGAAAGCTACCTCCAATCACATGTAAAT
AKT_MTRBTN2              TTCTCATACCACGGATCAAGAGCTGGTATGGGAGAAAGCTACCTCCAATCACATGTAAAT
                         ************************************************************ 1440

AKT_EDULIS               GATGGGGAAATTTTCTAG
AKT_TROSSULUS            GATGGGGAAATTTTCTAG
AKT_CANCER               GATGGGGAAATTTTCTAG
                         ****************** 1458


ALIGNMENTS P53-GENE_NUCLEOTIDE SEQUENCES

p53_TROSSULUS                       ATGTCACAAGCTTCAGTTTCAACTACATGCACACCATCAGGCCCACCAAT
p53_CANCER                          ATGTCACAAGCTTCAGTTTCAACTACATGCACACCATCAGGCCCACCAAT
p53_EDULIS                          ATGTCACAAGCTTCAGTTTCAACTACATGCACACCATCAGGCCCACCAAT
                                    **************************************************  50

p53_TROSSULUS                       GAGCCAAGAAACATTTGAATACTTATGGAATACTCTGGGAGAAGTCACAC
p53_CANCER                          GAGCCAAGAAACATTTGAATACTTATGGAATACTCTGGGAGAAGTCACAC
p53_EDULIS                          GGGCCAAGAAACATTTGAATATTTATGGAATACTCTGGGAGAGGTCACAC
                                    * ******************* ******************** *******  100

p53_TROSSULUS                       AAGAAGGGGGATACACAAACATCACATCTAAAGAATCAATCGACTATGCA
p53_CANCER                          AAGAAGGGGGTTACACAAACATCACATCTAAAGAATCAATCGACTATGCA
p53_EDULIS                          AAGAAGGGGGATACACAAACATCACATCTAAAGAATCAATCGACTATGCA
                                    ********** ***************************************  150

p53_TROSSULUS                       TTCAGTGAAGCTGAGGATGAAACCAGTATATCTGTTGAGAAATACAGAAT
p53_CANCER                          TTCAGTGAAGCTGAGGATGAAACCAGCATATCTGTTGAGAAATACAGAAT
p53_EDULIS                          TTCAGTGAAGCTGAGGATGAAACCAGTATATCTGTTGAGAAATACAGGAT
                                    ************************** ******************** **  200

p53_TROSSULUS                       TACATCCAATGACTCCATTTCAGACTTGTTGAATCCTATCATAGGTCAAA
p53_CANCER                          TACATCCAATGACTCCATTTCAGACTTGTTGAATCCTATCATAGGTCAAA
p53_EDULIS                          TACATCCAATGATTCCATTTCAGACTTGTTGAATCCTATAATAGGTCAAA
                                    ************ ************************** **********  250

p53_TROSSULUS                       CGACCTCAGCGAGCTCTATGAGCCCTGACAGTCAGACCAACATCATCGGT
p53_CANCER                          CGACCTCAGCGAGCTCTATGAGCCCTGACAGTCAGACCAACATCATCGGT
p53_EDULIS                          CAACTTCAGCTAGCTCTATGAGCCCTGACAGTCAGACCAATATCATCGGT
                                    * ** ***** ***************************** *********  300

p53_TROSSULUS                       AGCTCTGCATCAAGTCCTTACAATGATACAATCACATCACCTCCACCTTA
p53_CANCER                          AGCTCTGCATCAAGTCCTTACAATGATACAATCACATCACCTCCACCTTA
p53_EDULIS                          AGCTCTGCATCAAGTCCTTACAATGATACAATCACATCACCCCCACCTTA
                                    ***************************************** ********  350

p53_TROSSULUS                       TTCGCCTCATACTAGTATGCAGTCGCCTATACCCTCGGTGCCATCAAACA
p53_CANCER                          TTCGCCTCATACTAGTATGCAGTCGCCTATACCCTCGGTGCCATCAAACA
p53_EDULIS                          TTCGCCTCATACTAGTATGCAGTCACCTATACCCTCTGTGCCATCAAACA
                                    ************************ *********** *************  400

p53_TROSSULUS                       CTGACTATCCAGGAGATTATGGCTTCACCATTTCATTTTCTCAGCCATCA
p53_CANCER                          CTGACTATCCAGGAGATTATGGCTTCACCATTTCATTTTCTCAGCCATCA
p53_EDULIS                          CAGACTATCCCGGCGATTATGGGTTCACCATTTCATTTTCTCAGCCATCA
                                    * ******** ** ******** ***************************  450

p53_TROSSULUS                       AAAGAGACAAAGTCTACCACATGGACATACTCAGAGTCACTTAAGAAATT
p53_CANCER                          AAAGAGACAAAGTCTACCACATGGACATACTCAGAGTCACTTAAGAAATT
p53_EDULIS                          AAAGAGACCAAGTCTACCACATGGACATACTCAGAGTCACTTAAGAAATT
                                    ******** *****************************************  500

p53_TROSSULUS                       ATACGTCAGAATGGCAACAACTTGCCCAATCCGATTTAAATGTTTGAGAC
p53_CANCER                          ATACGTCAGAATGGCAACAACTTGCCCAATCCGATTTAAATGTTTGAGAC
p53_EDULIS                          ATACGTCAGAATGGCAACAACTTGCCCAATCCGATTTAAATGTTTGAGAC
                                    **************************************************  550

p53_TROSSULUS                       AGCCCCCACAGGGATGTGTAATTCGTGCAATGCCAATATTCATGAAACCT
p53_CANCER                          AGCCCCCACAGGGATGTGTAATTCGTGCAATGCCAATATTCATGAAACCT
p53_EDULIS                          AGCCCCCACAGGGATGTGTTATTCGTGCAATGCCAATATTCATGAAACCT
                                    ******************* ******************************  600


p53_TROSSULUS                       GAACATGTCCAAGAACCTGTAAAAAGATGCCCAAATCATGCCACATCTAA
p53_CANCER                          GAACATGTCCAAGAACCTGTAAAAAGATGCCCAAATCATGCCACATCTAA
p53_EDULIS                          GAACATGTCCAAGAACCTGTAAAAAGATGCCCAAATCATGCCACATCTAA
                                    **************************************************  650

p53_TROSSULUS                       AGAGCACAATGAAAATCATCCAGCCCCAACACATTTATGTCGATGTGAGC
p53_CANCER                          AGAGCACAATGAAAATCATCCAGCCCCAACACATTTATGTCGATGTGAGC
p53_EDULIS                          AGAGCACAATGAAAATCATCCAGCTCCAACACATTTATGTCGATGTGAGC
                                    ************************ *************************  700

p53_TROSSULUS                       ACAAACTTGCTAAATTTGTTGAAGACCCATACACAAGCCGCCAGAGTGTT
p53_CANCER                          ACAAACTTGCTAAATTTGTTGAAGACCCATACACAAGCCGCCAGAGTGTT
p53_EDULIS                          ACAAACTTGCTAAATTTGTTGAAGATCCATATACCAGCCGCCAGAGTGTT
                                    ************************* ***** ** ***************  750

p53_TROSSULUS                       CTAATTCCACATGAGATACCTCAAGCTGGCTCAGAATGGGTCACCAATTT
p53_CANCER                          CTAATTCCACATGAGATACCTCAAGCTGGCTCAGAATGGGTCACCAATTT
p53_EDULIS                          CTAATTCCACATGAGATACCTCAAGCTGGCTCAGAATGGGTCACCAATTT
                                    **************************************************  800

p53_TROSSULUS                       GTTCCAGTTCATGTGCCTGGGGTCATGTGTAGGAGGACCAAACAGAAGAC
p53_CANCER                          GTTCCAGTTCATGTGTCTGGGGTCATGTGTAGGAGGACCAAACAGAAGAC
p53_EDULIS                          GTTCCAGTTCATGTGCCTGGGGTCATGTGTAGGAGGACCAAACAGAAGGC
                                    *************** ******************************** *  850

p53_TROSSULUS                       CTATTCAGATTGTTCTGACTTTAGAAAAAGATAATCAAGTGCTAGGTAGA
p53_CANCER_HAPLOTYPE1_AetB          CTATTCAGATTGTTCTGACTTTAGAAAAAGATAATCAAGTGCTAGGTAGA
p53_CANCER                          CTATTCAGATTGTTCTGACTTTAGAAAAAGATAATCAAGTGCTAGGTAGA
p53_EDULIS                          CTATTCAGATTGTTCTGACTTTAGAAAAAGATAATCAAGTGCTAGGTAGA
                                    **************************************************  900

p53_TROSSULUS                       CGTGCCGTAGAAGTGAGAATTTGTGCCTGTCCTGGAAGAGATCGTAAGGC
p53_CANCER                          CGTGCCGTAGAAGTGAGAATTTGTGCCTGTCCTGGAAGAGATCGTAAGGC
p53_EDULIS                          CGGGCAGTAGAAGTTAGAATTTGTGCCTGTCCTGGGAGAGACAGAAAGGC
                                    ** ** ******** ******************** *****  * *****  950

p53_TROSSULUS                       TGATGAGAAGGCAGCTCTCCCACCATGTAAACAGTCCCCAAAGAAAGGCC
p53_CANCER                          TGATGAGAAGGCAGCTCTCCCACCATGTAAACAGTCCCCAAAGAAAGGCC
p53_EDULIS                          TGATGAGAAGGCAGCTCTCCCACCATGCAAACAGTCCCCAAAGAAAGGCC
                                    *************************** ********************** 1000

p53_TROSSULUS                       AGAAAGTTAATATTATCAATGAAATCACTACAGTAACACCAGGAGGCAAA
p53_CANCER                          AGAAAGTTAATATTATCAATGAAATCACTACAGTAACACCAGGAGGCAAA
p53_EDULIS                          AGAAAGTTAATATTATCAATGAAATCACTACAGTAACACCAGGAGGCAAA
                                    ************************************************** 1050

p53_TROSSULUS                       AAGAGGAAAGCAGAAGATGAACCCTTCACATTGTCTGTACGAGGACGAGA
p53_CANCER                          AAGAGGAAAGCAGAAGATGAACCCTTCACATTGTCTGTACGAGGACGAGA
p53_EDULIS                          AAGAGGAAAGCAGAAGACGAACCATTCACATTATCTGTACGAGGACGAGA
                                    ***************** ***** ******** ***************** 1100

p53_TROSSULUS                       AAACTACGAAATTCTGTGTAGACTGAGGGATTCATTGGAATTGTCATCCA
p53_CANCER                          AAACTACGAAATTCTGTGTAGACTGAGGGATTCATTGGAATTGTCATCCA
p53_EDULIS                          AAACTACGAAATTCTGTGTAGACTGAGGGATTCATTGGAACTGTCATCCA
                                    **************************************** ********* 1150

p53_TROSSULUS                       TGGTTCCCCAGAATCAAATAGATGTATACAAACAGAAACAACTGGATACA
p53_CANCER                          TGGTTCCCCAGAATCAAATAGATGTATACAAACAGAAACAACTGGATACA
p53_EDULIS                          TGGTTCCCCAGAATCAAATAGATGTATACAAACAGAAACAACTTGATACA
                                    ******************************************* ****** 1200

p53_TROSSULUS                       AATAGACAATCATCGCCAACAACA------GCAAGGGTTGTTACTCTTCC
p53_CANCER                          AATAGACAATCATCGCCAACAACA------GCAAGGGTTGTTACTCTTCC
p53_EDULIS                          AACAGACAATCATCGCCAACAACATCAACAGCAAGGGTTGTTACTCTTCC
                                    ** *********************      ******************** 1250

p53_TROSSULUS                       TACTCACGACAACACACCAATAACTATACAGGGTGAAGGTCGGCAGACGA
p53_CANCER                          TACTCACGACAACTCACCAATAACTATACAGGGTGAAGGTCGGCAGACGA
p53_EDULIS                          TACTCACGACAACACACCAATAACTATACAGGGTGAAGGTCGGCAGACGA
                                    ************* ************************************ 1300

p53_TROSSULUS                       CACTACCATTCACTGCAGACTTAAATGGTCAGGTGACAAGTTCACAAAAT
p53_CANCER                          CACTACCATTCACTGCAGACTTAAATGGTCAAGTGACAAGTTCACAAAAT
p53_EDULIS                          CACTACCATTCACTGCAGACTTAAATGGTCAGGTGACAAGTTCACAAAAT
                                    ******************************* ****************** 1350


p53_TROSSULUS                       GGTGTGGTGGAAAATCATGGAAATATTAAAGAAGAACTGATGGCCAATGG
p53_CANCER_HAPLOTYPE1_AetB          GGTGTGGTGGAAAATCATGGAAATATTAAAGAAGAACTGATGGCCAATGG
p53_EDULIS                          GGTGTGGTGGAAAATCATGGAAATATTAAAGAAGAACTGATGGCTAATGG
                                    ******************************************** ***** 1400

p53_TROSSULUS                       GGACCATTCAATTTCAAATTGGTTGACTACTCTTGGTTTATCAGCATACA
p53_CANCER                          GGACCATTCAATTTCAAATTGGTTGACTACTCTTGGTTTATCAGCATACA
p53_EDULIS                          GGACCATTCTATTTCAAATTGGTTGACTACCCTTGGTTTATCAGCCTACA
                                    ********* ******************** ************** **** 1450

p53_TROSSULUS                       TAGATAACTTTCATCAACAGAATCTGTTTACAATGGAGCAGCTTGATGAT
p53_CANCER                          TAGATAACTTTCATCAACAGAATCTATTTACAATGGAGCAGCTTGATGAT
p53_EDULIS                          TCGATAACTTTCATCAACAGAATCTATTTACAATGGAGCAACTCGATGAT
                                    * *********************** ************** ** ****** 1500

p53_TROSSULUS                       TTTACTGTTGAGGACTTGCAAAAGATGAGAATAGGAACAAGCCACAGAAA
p53_CANCER                          TTTACTGTTGAGGACTTGCAAAAGATGAGAATAGGAACAAGCCACAGAAA
p53_EDULIS                          TTTACTGTTGAGGACTTACAAAAGATGAGAATAGGAACAAGCCACAGAAA
                                    ***************** ******************************** 1550

p53_TROSSULUS                       CAAAATCTGGAAAGCTCTAGTAGAATTCCATAGCGAGTCTATCACCATTA
p53_CANCER                          CAAAATCTGGAAAGCTCTAGTAGAATTCCATAGCGAGTCTATCACCATTA
p53_EDULIS                          TAAAATCTGGAAAGCTCTAGTAGAATTCCATAGCGAGTCTATTACCATTA
                                     ***************************************** ******* 1600

p53_TROSSULUS                       GTGATTCACAAAGTCTGCAGAGGGACGTCAGCACTGCATCCACCATTAGT
p53_CANCER                          GTGATTCACAAAGTCTGCAGAGGGACGTCAGCACTGCATCCACCATTAGT
p53_EDULIS                          GTGATTCACAAAGTCTGCAGAGGGACGTCAGCACTGCATCCACCATTAGT
                                    ************************************************** 1650

p53_TROSSULUS                       ATGACATCACAGGCCAGTATCTCCCAGAATTCTCAAAATTCAACGTATTG
p53_CANCER                          ATGACATCACAGGCCAGTATCTCCCAGAATTCTCAAAATTCAACGTATTG
p53_EDULIS                          ATGACATCACAGGCCAGTATCTCCCAGAATTCTCAAAATTCAACGTATTG
                                    ************************************************** 1700

p53_TROSSULUS                       TCCAGGATACTATGAAGTTACCAGATATACATTCAAACATACAGTTTCTT
p53_CANCER                          TCCAGGATACTATGAAGTTACCAGATATACATTCAAACATACAGTTTCTT
p53_EDULIS                          TCCGGGATACTATGAAGTTACCAGATATACATTCAAACATACAGTTTCTT
                                    *** ********************************************** 1750

p53_TROSSULUS                       ATACGAAAGTGGATGAAAGAAGTCCAAAACGGGCTAAAGTAGACTGA
p53_CANCER                          ATACGAAAGTGGATGAAAGAAGTCCAAAACGGGCTAAAGTAGACTGA
p53_EDULIS                          ATACGAAAGTGGATGAAAGAAGTCCAAAACGGGCTAAGGTAGACTGA
                                    ************************************* ********* 1797


ALIGNMENTS MDM2-GENE_NUCLEOTIDE SEQUENCES


MDM2_TROSSULUS             ATGTGCAATAATCCAAATTTAATTGTACGACAGCCTCCTGATACATACAG
MDM2_EDULIS                ATGTGCAATAATCCAAATTTAATTGTACGACAGCCTCCTGATACATACAG
MDM2_CANCER                ATGTGCAATAATCCAAATTTAATTGTACGACAGCCTCCTGATACATACAG
                           **************************************************  50

MDM2_TROSSULUS             CACACCTGCAGGATCAAATGTATCTACTTTGAAGCAACTCACAGAGTCCC
MDM2_EDULIS                CACACCTGCAGGATCAAATGTATCTACTTTGAAGCAACTCACAGAGTCCC
MDM2_CANCER                CTCAGCTGCAGGATCAAATGTATCTACTTTGGAGCAACTTACAGAGTCCC
                           ** **************************** ******* **********  100

MDM2_TROSSULUS             ATTACATAACAACTGAAGAATGTGAGGTCAATTATCCTATGGTCAGACCC
MDM2_EDULIS                ATTACATAACAACTGAAGAATGTGAGGTCAATTATCCTATGGTCAGACCC
MDM2_CANCER                ATTACATAACAACTGAAGAATGTGAGGTCAATTATCCTATGGTCAGACCC
                           **************************************************  150

MDM2_TROSSULUS             AGACGTGAATTTCTTGACCTACTTAAACAAGTAGGTGCATCATGGGAAGT
MDM2_EDULIS                AGACGTGAATTTCTTGACCTACTTAAACAAGTAGGAGCATCATGGGAAGT
MDM2_CANCER                AGACGTGAATTTCTTGACCTACTTAAACAAGTAGGTGCATCATGGGAAGT
                           ************************************************** 200

MDM2_TROSSULUS             GTTATCCAGAAAAGAGGTCTGTAGTTATCTGAAGGATTACATTGGTAGCA
MDM2_EDULIS                GTTATCCAGAAAAGAGGTCTGTAGTTATCTGAAGGATTACATTGGTAGCA
MDM2_CANCER                GTTATCCAGAAAAGAGGTCTGTAGTTATCTGAAGGATTACATTGGTAGCA
                           ************************************************** 250

MDM2_TROSSULUS             GACAGTTGTATAATCCAAATGACCCTAGTATCGTGTTCTGTAAAAAAGAT
MDM2_EDULIS                GACAGTTGTATAATCCAAATGACCCTAGTATCGTGTTCTGTAAAAAAGAT
MDM2_CANCER                GACAGTTGTATAATCCAAATGACCCTAGTATCGTGTTCTGTAAAAAAGAT
                           ************************************************** 300

MDM2_TROSSULUS             CCTCTGGGAAAAGTATTCGGAGTTGATAAGTTTACCTTCACAGAAGCAAC
MDM2_EDULIS                CCTCTGGGAAAAGTATTTGGAGTTGATAAGTTTACCTTCACAGAAGCAAC
MDM2_CANCER                CCTCTGGGAAAAGTATTTGGAGTTGAAAAGTTTACCTTCACAGAAGCAAC
                           ***************** ******** *********************** 350

MDM2_TROSSULUS             GAAATTGATAGATGCCAACTGCTACCAGTTACCTGACTCATGTATGGTTA
MDM2_EDULIS                GAAATTGATAGATGCCAACTGCTACCAGTTACCTGACTCATGTATGGTTA
MDM2_CANCER                GAAATTGATAGATGCCAACTGCTACCAGTTACCTGACTCATGTATGGTTA
                           ************************************************** 400

MDM2_TROSSULUS             AGAAAGAACAATTAGTCGCTATACCAAAGAAAAGAAGTAATTCATCAAAT
MDM2_EDULIS                AGAAAGAACAATTAGTCGCTATACCAAAGAAAAGAAGTAATTCATCAAAT
MDM2_CANCER                AGAAAGAACAATTAGTCGCTATACCAAAGAAAAGAAATAATTCATCAAAT
                           ************************************ ************* 450

MDM2_TROSSULUS             CAAGTAAATCAGATAGTGACTGTTTCTTGTCCAAAGTTGATTGACATACT
MDM2_EDULIS                CAAGTAAATCAGATAGTGACTGTTTCTTGTCCAAAGTTGATTGACATACT
MDM2_CANCER                CAAGTAAATCAGATAGTGACTGTTTCTTGTCCAAAGTTGATTGACATACT
                           ************************************************** 500

MDM2_TROSSULUS             CGTACATAGTCCCAGTACACAGACAGGGGAACAGCAGAAAAAACAAAATA
MDM2_EDULIS                CGTACATAGTCCCAGTACACAGACAGGGGAACAGCAGAAAAAACAAAATA
MDM2_CANCER                CGTACATAGTCCCAGTACACAGACAGGGGAACAACAGAAAAAACAAAATA
                           ********************************* **************** 550

MDM2_TROSSULUS             GTTTCGAAGGTCAAACATCAGACACGAGTTCTAACAGAGCACAAAACTTG
MDM2_EDULIS                GTTTGGAAGGTCAAACATCAGACACGAGTTCTAACAGAGCACAAAACTTG
MDM2_CANCER                GTTTCGAAGGTCAAACATCAGACACGAGTTCTAACAGAGCACAAAACTTG
                           **** ********************************************* 600

MDM2_TROSSULUS             TCAGCTGCATCTACTGGTAGGAGAAGACATAGATGGGTTTCTGAAAGGGG
MDM2_EDULIS                TCAGCTGCATCTACTGGTAGGAGAAGACATAGATGGGTTTCTGAAAGGGG
MDM2_CANCER                TCAGCTGCATCTACTGGTAGGAGAAGACATAGATGGGTTTCTGAAAGGGG
                           ************************************************** 650

MDM2_TROSSULUS             AGATTCTGAACCAGATATAATAGCCATCGAGCTAGATGCTGACAATGATG
MDM2_EDULIS                AGATTCTGAACCAGATATAATAGCCATCGAGCTAGATGCTGACAATGATG
MDM2_CANCER                AGATTCAGAGCCAGATATAATAGCCATAGAGTTAGATGCTTACAATGATG
                           ****** ** ***************** *** ******** ********* 700

MDM2_TROSSULUS             ATGATGCTATAGGTGTAGAGGTTGGAACCAATGGTACTGAGGATGTTATG
MDM2_EDULIS                ATGATGCTATAGGTGTAGAGGTTGGAACCAATGGTACTGATGATGTTATG
MDM2_CANCER                AGGAAGCTATAGGTGTAGAGGTTGAAACCAATGGTACTGAGGATGTTATG
                            *** ******************* *************** ********* 750

MDM2_TROSSULUS             TCTGTAGAGGTTGATGCTGACAGAATCTCATTCGAGTATGAAGTCTGTTC
MDM2_EDULIS                TCTGTAGAGGTTGATGCTGACAGAATCTCATTCGAGTATGAAGTCTGTTC
MDM2_CANCER                TCTGTAGAGGTGGATGCTGACAGAATCTCATTCGAGTATGAAGTCTGTTC
                           *********** ************************************** 800

MDM2_TROSSULUS             AGATGAGGATGATATTATATCCAGCACAGCCAGTGTATCAGGAATCTCAG
MDM2_EDULIS                AGATGAGGATGATATTATATCCAGCACAGCCAGTGTATCAGGAATCTCAG
MDM2_CANCER                AGATGAGGATGATATCATATCCAGCACAGCCAGTGTATCAGGAATCTCAG
                           *************** ********************************** 850

MDM2_TROSSULUS             ATACTAATGTAGTGATAATATGTGGTGATAGTGATATGGAATTCTGGGGT
MDM2_EDULIS                ATACTAATGTAGTGATAATATGTGGTGATAGTGATATGGAATTCTGGGGT
MDM2_CANCER                ATACTAATGTAGTGATAATATGTGGTGATAGTGATATGGAATTCTGGGGT
                           ************************************************** 900

MDM2_TROSSULUS             GATTCTGATTCAAGTGATTCAGAATTATCAGATGGGGATAAATGGACGTG
MDM2_EDULIS                GATTCTGATTCAAGTGATTCAGAATTATCAGATGGGGATAAATGGACGTG
MDM2_CANCER                GATTCTGATTCAAGTGATTCAGAATTATCAGACGGGGATAAATGGACATG
                           ******************************** ************** ** 950

MDM2_TROSSULUS             TACAGAATGTGATACAAAAAATTCTCCTTTAAATGGTCACTGCGGACATT
MDM2_EDULIS                TACAGAATGTGATACAAAAAATTCTCCTTTAAATGGTCACTGCGGACATT
MDM2_CANCER                TACAGAATGTGATACAAAAAATTCTCCTTTAAATGGTCACTGCGGACATT
                           ************************************************** 1000


MDM2_TROSSULUS             GTAAAAAAGTACGTCCAAACTGGTTACCAGAGACTACAAGAAAGAGACGT
MDM2_EDULIS                GTAAAAAAGTACGTCCAAACTGGTTACCAGAGACTACAAGAAAGAGACGT
MDM2_CANCER                GTAAAAAAGTACGTCCAAACTGGTTACCAGAGACTACAAGAAAGAGACGT
                           ************************************************** 1050

MDM2_TROSSULUS             TTGGAAAAAATGAACTCGAAAGATAAGGCACTTTTATCTTCTTCTGAATG
MDM2_EDULIS                TTGGAAAAAATGAACTCGAAAGATAAGGCACTTTTATCTTCTTCTGAATG
MDM2_CANCER                TTGGAAAAAATGAACTCGAAAGATAAGGCACTATTATCTTCTTCTGAATG
                           ******************************** ***************** 1100

MDM2_TROSSULUS             TAGTGACATATCGCACATCTCTTCAAAAATTCAACGTCGTTCTTCAAGAG
MDM2_EDULIS                TAGTGACATATCGCACATCTCTTCAAAAATTCAACGTCGTTCTTCAAGAG
MDM2_CANCER                TAGTGACATATCGCACATCTCTTCAAAAATTCAACGTCGTTCTTCAAGAG
                           ************************************************** 1150

MDM2_TROSSULUS             ACCGTCTTACGTCGATAGAATTGGAATCGGACAAGGATTTAATAGCTGAG
MDM2_EDULIS                ACCGTCTTACGTCGATAGAATTGGAATCGGACAAGGATTTAATAGCTGAG
MDM2_CANCER                ACCGTCTTACATCGATAGAATTGGAATCGGACAAGGATTTAATAGCTGAG 1200
                           ********** ***************************************

MDM2_TROSSULUS             GAAGAGGACTCAGGAATTAGTACTCTGTCCAGACAGCAATCTAGTCAAGA
MDM2_EDULIS                GAAGAGGACTCAGGAATTAGTACTCTGTCCAGACAGCAATCTAGTCAAGA
MDM2_CANCER                GAAGAGGACTCAGGGATTAGTACTCTGTCCAGACAGCAATCTAGTCAAGA
                           ************** *********************************** 1250

MDM2_TROSSULUS             AAATTTAGCCGAAAATAAACTGAAATCATATTCGTCACTGATAGAAAATT
MDM2_EDULIS                AAATTTAGCCGAAAATAAACTGAAATCATATTCGTCACTGATAGAAAATT
MDM2_CANCER                AAATTTAGCCGAAAATAAACTGAAATCATATTCGTCACTGATAGAAAATT
                           ************************************************** 1300

MDM2_TROSSULUS             CTTCCACGTTAGGATCGTTTAGAACTTTGTTAAACTTTCAAACAACAGCA
MDM2_EDULIS                CTTCCACGTTAGGATCGTTTAGAACTTTGTTAAACTTTCAAACAACAGCA
MDM2_CANCER                CTTCCACGTTAGGATCATTTAGAACTTTGTTAAACTTTCAAACAACAGCA
                           **************** ********************************* 1350

MDM2_TROSSULUS             TCGGCAAAAAATAAAGATGAAATGACAATATCAAATAAAATAGATAAATT
MDM2_EDULIS                TCGGCAAAAAATAAAGATGAAATGACAATATCAAATAAAATAGATAAATT
MDM2_CANCER                TCGGCAAAAAATAATGATGAAAGAATAAAATCAAATAAAAATGATAATTT
                           **********************  * ** ***********  ***** **  1400

MDM2_TROSSULUS             GGACAAACCACAAAAAGATTTGAAGCTTGAGAAAAGTTCAAACATTAAGG
MDM2_EDULIS                GGACAAACCACAAAAAGATTTGAAGCTTGAGAAAAGTTCAAACATTAAGG
MDM2_CANCER                GGACAAACCACAAAAAGATTTGAAGCTTGAGAAAAGTTCAAACATTAAGG
                           ************************************************** 1450

MDM2_TROSSULUS             AATTCGAAAATTCTCAATCACAACTCGGTCAGTGCTGTATCTGTTTCAGC
MDM2_EDULIS                AATTCGAAAATTCTCAATCACAACTCGGTCAGTGCTGTATCTGTTTCAGC
MDM2_CANCER                AATTCGAAAATTCTCAATCACAACTCGGTCAGTGCTGTATCTGTTTCAGC
                           ************************************************** 1500

MDM2_TROSSULUS             CGGCCTAAAACTGCCAGCATTATTCATGGCTGCACTGGACATCAAGTATG
MDM2_EDULIS                CGGCCTAAAACTGCCAGCATTATTCATGGCTGCACTGGACATCAAGTATG
MDM2_CANCER                CGGCCTAAAACTGCCAGCATTATTCATGGCTGCACTGGACATCAAGTATG
                           **************************************************1550

MDM2_TROSSULUS             TTGTTATAGATGTGCCAAGCGACTGAAAAGACTTGCAAAACCATGCCCCC
MDM2_EDULIS                TTGTTATAGATGTGCCAAGCGACTGAAAAGACTTGCAAAACCATGCCCCC
MDM2_CANCER                TTGTTATAGATGTGCCAAGCGACTGAAAAGACTTGCAAAACCATGCCCCC
                           ************************************************** 1600

MDM2_TROSSULUS             TTTGTAGACGACCTATTCAAAAGGTCATTAAAAACTATTCTGCTTAA
MDM2_EDULIS                TTTGTAGACGACCTATTCAAAAGGTCATTAAAAACTATTCTGCCTAA
MDM2_CANCER                TTTGTAGACGACCTATTCAAAAGGTCATTAAAAACTATTCTGCTTAA
                           ******************************************* *** 1647

ALIGNMENTS MYC-GENE_NUCLEOTIDE SEQUENCES

MYC_CANCER                ATGATGCGTAAGTACTCTGAAATAGTCATACCAAGACGTGAAGGGAACAGTTGCCAAGTT
MYC_TROSSULUS             ATGATGCGTAAGTACTCTGAAATAGTCATACCAAGACGTGAAGGGAACAGTTGCCAAGTT
MYC_EDULIS                ATGATGCGTAAGTACTCTGAAATAGTCATACCAAGACGTGAAGGGAACAGTTGCCAAGTT
                          ************************************************************  60

MYC_CANCER                GTCACCGAAATTGGACATACTGGAATAGCTCAACCTCCTGACAATTCAGTGAAGATGGAT
MYC_TROSSULUS             GTCACCGAAATTGGACATACTGGAATAGCTCAACCTCCTGACAATTCAGTGAAGATGGAT
MYC_EDULIS                GTCACCGAAATTGGACATACTGGAATAGCTCAACCTTCTGACAATTCAGTGAAGATGGCT
                          ************************************ ********************* *  120

MYC_CANCER                AAAATGTGCAAACACTGTTACGTGGATAATTCACATGTTCTTCAACATTGCATGTATGAA
MYC_TROSSULUS             AAAATGTGCAAACACTGTTACGTGGATAATTCACATGTTCTTCAACATTGCATGTATGAA
MYC_EDULIS                AAAATGTGCAAACACTGTTACGTGGATAATTCACATGTTCTTCAACATTGCATGTATGAA
                          ************************************************************  180

MYC_CANCER                ACAGAATATTCATCACCAGCATCTCTCCCAAGTGATGATATTTGGAAGAAGTTTGAGTTG
MYC_TROSSULUS             ACAGAATCTTCATCACCAGCATCTCTCCCAAGTGATGATATTTGGAAGAAGTTTGAGTTG
MYC_EDULIS                ACAGAATCATCATCACCAGCATCTCTCCCAAGTGATGATATTTGGAAGAAGTTTGAGTTG
                          *******  ***************************************************  240

MYC_CANCER                ATTCCAACACCTCCTCGCTCACCAGAAAGAGAAGAACCGTTTGAAATGGATTTAAGCTTG
MYC_TROSSULUS             ATTCCAACACCTCCTCGCTCACCAGAAAGAGAAGAACCGTTTGAAATGGATTTAAGCTTG
MYC_EDULIS                ATTCCAACACCTCCTCGGTCACCAGAAAGAGAAGAACCGTTTGAAATGGATTTAAGCTTG
                          ***************** ******************************************  300

MYC_CANCER                GATTTAGAAGATATGAATTTCCCATTTGATGCAAATTTCTTTGAAAGTGAGGATTTTAAT
MYC_TROSSULUS             GATTTAGAAGATATGAATTTCCCATTTGATGCAAATTTCTTTGAAAGTGAGGATTTTAAT
MYC_EDULIS                GATTTAGAAGATATGAATTTCCCATTTGATGCAAATTTCTTTGAAAGTGAGGATTTTAAT
                          ************************************************************  360

MYC_CANCER                CCGAAAGAAACTTTTGCCGAGTCGCCTCTGTCACAATTGTGCTCTAAATTAATTCAGGAT
MYC_TROSSULUS             CCGAAAGAAACTTTTGCCGAGTCGCCTCCGTCACAATTGTGCTCTAAATTAATTCAGGAT
MYC_EDULIS                CCGAAAGAAACTTTTGCCGAGTCGCCTCCGTCACAATTGTGCTCTAAATTAATTCAGGAT
                          **************************** *******************************  420

MYC_CANCER                TGTATGTGGTCTGGAGATACATTTTCTTCCAGTGAGCAGAAGGATAAAACATCTTCATCT
MYC_TROSSULUS             TGTATGTGGTCTGGAGATACATTTTCTTCCAGTGAGCAGAAGGATAAAACATCTTCATCT
MYC_EDULIS                TGTATGTGGTCTGGAGATACATTTTCTTCCAGTGAGCAAAAGGATAAAACATCTTCATCT
                          ************************************** *********************  480

MYC_CANCER                TTTGACATGAAAACTGTTGATCCAATGGCAGTTTTTCCATGTTCAGTAACCAACAATACA
MYC_TROSSULUS             TTTGACATGAAAACTGTTGATCCAATGGCAGTTTTTCCATGTTCAGTAACCAACAATACA
MYC_EDULIS                TTTGACATGAAAACTGTTGATCCAATGGCAGTTTTTCCATGTTCAGTAACCAACAATACA
                          ************************************************************  540

MYC_CANCER                AACAATTCTCACAGCATTATGGGCTACTTGAGTGAAACAAAGCTTCACAGTCTTGGCACA
MYC_TROSSULUS             AACAATTCTCACAGCATTATGGGCTACTTGAGTGAAACAAAGCTTCACAGTCTTGGCACA
MYC_EDULIS                AACAATTCACACAGCATTATGGGCTACTTGAGTGAAACAAAGCTTCACAGTCTTGGCACA  600
                          ******** ***************************************************

MYC_CANCER                GAAACACCTTCAGATTCAGAAGAAGAAATTGATGTTGTCACCGTTGAAAAACTGCAACAG
MYC_TROSSULUS             GAAACACCTTCAGATTCAGAAGAAGAAATTGATGTTGTCACCGTTGAAAAACTGCAACAG
MYC_EDULIS                GAAACACCTTCAGATTCAGAAGAAGAAATTGATGTTGTCACAGTTGAAAAACTGCAACAG
                          ***************************************** ******************  660

MYC_CANCER                AACTCAACAACTGCTGGACCAACTATCAACAAGTCAAATATTCAGACAAGGAATATAAAA
MYC_TROSSULUS             AACTCAACAACTGCTGGACCAACTATCAACAAGTCAAATATTCAGACAAGGAATATAAAA
MYC_EDULIS                AACTCAACAACTGCTGGACCAACTATCAACAAGTCAAATATTCAGACAAGGAATATAAAA
                          ************************************************************  720

MYC_CANCER                ACACAAATTAAAATTGAAAAGAACAATGGAGAGAAAGTGGAAATAATTCCTTTAAAAACA
MYC_TROSSULUS             ACACAAATTAAAATTGAAAAGAACAATGGAGAGAAAGTGGAAATAATTCCTTTAAAAACA
MYC_EDULIS                ACACAAATTAAAATTGAAAAAAACAATGGAGAGAAAGTGGAAATAATTCCTTTAAAAACA
                          ******************** ***************************************  780

MYC_CANCER                ACAACATTAAAAATCCGAGTAGAAACACCAGACTTGCACAATTATAGTTTACCACATTCA
MYC_TROSSULUS             ACAACATTAAAAATCCGAGTAGAAACACCAGACTTGCACAATTATAGTTTACCACATTCA
MYC_EDULIS                ACAACATTAAAAATCCGAGTAGAAACCCCAGACTTGCATAACTATAGTTTACCACATTCA
                          ************************** *********** ** ******************  840

MYC_CANCER                CATCAACCTAAACGTGTGCGGTCTTATCCAGTGTCTCCATCACATTCACCATTACCTCAG
MYC_TROSSULUS             CATCAACCTAAACGTGTGCGGTCTTATCCAGTGTCTCCATCACATTCACCATTACCTCAG
MYC_EDULIS                CATCAACCTAAACGTGTGCGGTCTTATCCAGTGTCTCCATCACATTCACCATTACCTCAG
                          ************************************************************  900

MYC_CANCER                AAAAGGACTAAAAAGGATTTAAGTGTGCCTGACTTCAAACGTGTTTGTCAAAAATTACGT
MYC_TROSSULUS             AAAAGGACTAAAAAGGATTTAAGTGTGCCTGACTTCAAACGTGTTTGTCAAAAATTACGT
MYC_EDULIS                AAAAGGACTAAAAAGGATTTAAGTGTGCCTGACTTCAAACGTGTTTGTCAAAAATTACGT
                          ************************************************************  960

MYC_CANCER                GCAAGCAAAAGTTCTTCGGACTCAGAAGAGTGTATAAGTGAAGGTGGAAAACGGACACAA
MYC_TROSSULUS             GCAAGCAAAAGTTCTTCGGACTCAGAAGAGTGTATAAGTGAAGGTGGAAAACGGACACAA
MYC_EDULIS                GCAAGCAAAAGTTCTTCTGACTCTGAAGAGTGTATAAGTGAAGGTGGAAAACGGACACAA
                          ***************** ***** ************************************ 1020


MYC_CANCER                CACAATGTGCTTGAAAGGAAACGAAGAAATGACTTGAAATACAGTTTCTTTACATTACGT
MYC_TROSSULUS             CACAATGTGCTTGAAAGGAAACGAAGAAATGACTTGAAATACAGTTTCTTTACATTACGT
MYC_EDULIS                CACAATGTGCTTGAAAGGAAACGAAGAAATGACTTGAAATATAGTTTCTTTACATTACGT
                          ***************************************** ****************** 1080

MYC_CANCER                GACAGTGTCCCAGAACTTAGTAACCAGGAACGTGCTCCTAAAGTGCTCATTCTAAAGAAA
MYC_TROSSULUS             GACAGTGTCCCAGAACTTAGTAACCAGGAACGTGCTCCTAAAGTGCTCATTCTAAAGAAA
MYC_EDULIS                GACAGTGTCCCAGAACTTAGTAACCAGGAACGTGCTCCTAAAGTGCTCATTCTAAAGAAA
                          ************************************************************ 1140

MYC_CANCER                GCCTCAGACTATGTACATTCTTTGAATGTAGACAACAAAAGACTAGAAAGTGAAAAGGCT
MYC_TROSSULUS             GCCTCAGACTATGTACATTCTTTGAATGTAGACAACAAAAGACTAGAAAGTGAAAAGGCT
MYC_EDULIS                GCTTCAGACTATGTACATTCTTTGAATGTAGACAACAAAAGACTAGAAAGTGAAAAGGCT
                          ** ********************************************************* 1200

MYC_CANCER                ACTTTGTTAGCTAAACAGCAGAAATTGAAACGGACTTTAGAAATTCTACAGGATAGTGAC
MYC_TROSSULUS             ACTTTGTTAGCTAAACAGCAGAAATTGAAACGGACTTTAGAAATTCTACAGGATAGTGAC
MYC_EDULIS                ACTTTGTTAGCTAAACAGCAGAAATTGAAACGGACTTTAGAAATTTTACAGGATAGTGAC
                          ********************************************* ************** 1260

MYC_CANCER                TTTTTCTGA
MYC_TROSSULUS             TTTTTCTGA
MYC_EDULIS                TTTTTCTGA
                          ********* 1269


ALIGNMENTS FBXW7-GENE_NUCLEOTIDE SEQUENCES

FBXW7_HAPLOTYPE-B            ATGGCGAGCAAGAAAATAGATATCAACAAGGCTAGTGTAGAAGACTTTGA
FBXW7_TROSSULUS              ATGGCGAGCAAGAAAATAGATATCAACAAGGCTAGTGTAGAAGACTTTGA
FBXW7_EDULIS                 ATGGCAAGCAAGAAAATAGATATCAACAAGGCTAGTGTAGAAGACTTTGA
                             ***** ********************************************  50

FBXW7_HAPLOTYPE-B            AAAAGTAAATGGTATTGGTTTAAGAAAAGCTAAAGCTATCATCAAATACA
FBXW7_TROSSULUS              AAAAGTAAATGGTATTGGTTTAAGAAAAGCTAAAGCTATCATCAAATACA
FBXW7_EDULIS                 AAAAGTAAATGGTATTGGTTTAAGAAAAGCAAAAGCTATCATCAAATACA
                             ****************************** *******************  100

FBXW7_HAPLOTYPE-B            GAAAGTCATTTGGAGGTTTCTCAACGGTAAAGGATCTGAGTAGTGTCCCA
FBXW7_TROSSULUS              GAAAGTCATTTGGAGGTTTCTCAACGGTAAAGGATCTGAGTAGTGTCCCA
FBXW7_EDULIS                 GAAAGTCATTTGGAGGTTTCTCAACCGTAAAGGATTTGAGTAGTGTCCCA
                             ************************* ********* **************  150

FBXW7_HAPLOTYPE-B            GGAGTTGGGGATTGTATTGTGGAACAAAGTCGTACTCAGCTGACATGTAG
FBXW7_TROSSULUS              GGAGTTGGGGATTGTATTGTGGAACAAAGTCGTACTCAGCTGACATGTAG
FBXW7_EDULIS                 GGAGTTGGGGATTGTATTGTGGAACAAAGTCGTACTCAGTTGACATGTAG
                             *************************************** **********  200

FBXW7_HAPLOTYPE-B            TAATCGCAAACGTAAAGCTGTCAGGACACCTAAAGTAAAAGATAATGTTT
FBXW7_TROSSULUS              TAATCGCAAACGTAAAGCTGTCAGGACACCTAAAGTAAAAGATAATGTTT
FBXW7_EDULIS                 TAACTGCAAACGCAAAGCTGTCAGGACACCTAAAGTAAAAGATAATGTTT
                             ***  ******* *************************************  250

FBXW7_HAPLOTYPE-B            GTAGGGAACAGCCATCATCTAGTGGGTTATCAGCCAGATCTATCAGACAC
FBXW7_TROSSULUS              GTAGGGAACAGCCATCATCTAGTGGGTTATCAGCCAGATCTATCAGACAC
FBXW7_EDULIS                 GTAAGGAACAGCCATCATCTAGTGGGTTATCAGCCAGATCTATCAGACAC
                             *** **********************************************  300

FBXW7_HAPLOTYPE-B            AATCAACGAGCACTGTTCAAATCAACAGAGGGAAATGTTGATGATTGTAA
FBXW7_TROSSULUS              AATCAACGAGCACTGTTCAAATCAACAGAGGGAAATGTTGATGATTGTAA
FBXW7_EDULIS                 AATCAACGAGCACTGTTCAAATCAACAGAGGAACATGTTGATGATGGTAA
                             ******************************* * *********** ****  350

FBXW7_HAPLOTYPE-B            CAGTATTGTCTGCCAGGAAGGTCACATGTCAGAGGAAGAGTCTGGCGATG
FBXW7_TROSSULUS              CAGTATTGTCTGCCAGGAAGGTCACATGTCAGAGGAAGAGTCTGGCGATG
FBXW7_EDULIS                 CAGTATTGTATCCCACGAAGGTCACATGTCAGAGGAAGAGTCTGGCGATG
                             ********* * *** **********************************  400

FBXW7_HAPLOTYPE-B            AGGTTATGTTCCGTGAAGAAGACAAGTCTAATGCTCAACATAAATCTAGG
FBXW7_TROSSULUS              AGGTTATGTTCCGTGAAGAAGACAAGTCTAATGCTCAACATAAATCTAGG
FBXW7_EDULIS                 AGGTTATGTTCCGTGAAGAAGACAGGTCTAATGCTCAACATAAATCTAGG
                             ************************ *************************  450

FBXW7_HAPLOTYPE-B            ATTCCTATACCAGTCAATGGACTCTTCAGTAGCCATGTGATGAAAAGAAA
FBXW7_TROSSULUS              ATTCCTATACCAGTCAATGGACTCTTCAGTAGCCATGTGATGAAAAGAAA
FBXW7_EDULIS                 ATTCCTATACCAGTCAATGGACTCTTCAGTAGCCATGTGATGAAAAGAAA
                             **************************************************  500

FBXW7_HAPLOTYPE-B            AGGTCATGTGGTAAAGGAAGAGCCTGACTCGTCTCCTGGAAAAAAATTCT
FBXW7_TROSSULUS              AGGTCATGTGGTAAAGGAAGAGCCTGACTCGTCTCCTGGAAAAAAATTCT
FBXW7_EDULIS                 AGGTCATGTGGTAAAGGAAGAGCCTGACTCGTCTCCTGGGAAAAAATTCT
                             *************************************** **********  550

FBXW7_HAPLOTYPE-B            GTCGAGATGGTGATTATCAAAGTACCTTTGTGAACTTACAAAAGTCATCA
FBXW7_TROSSULUS              GTCGAGATGGTGATTATCAAAGTACCTTTGTGAACTTACAAAAGTCATCA
FBXW7_EDULIS                 GTCGAGATGGTGATTATCAAAGTACCTTTGTGAACTTACAAAAGTCATCA
                             **************************************************  600

FBXW7_HAPLOTYPE-B            AGACAGAATTTAATTGCTACACAACGAATGAGGATTCCCTCCAAAGAGCA
FBXW7_TROSSULUS              AGACAGAATTTAATTGCTACACAACGAATGAGGATTCCCTCCAAAGAGCA
FBXW7_EDULIS                 CGACAGAATTTAATTGCTACACAACGAATGAGGATTCCCTCCAAAGAGCA
                              *************************************************  650

FBXW7_HAPLOTYPE-B            TCCGCCAGACAAACTAGCAGAATGGCTGCAAATATTTCAGAATTGGGGTA
FBXW7_TROSSULUS              TCCGCCAGACAAACTAGCAGAATGGCTGCAAATATTTCAGAATTGGGGTA
FBXW7_EDULIS                 TCCTCCAGACAAATTAGCGGAATGGCTGCAAATATTTCAGAATTGGGGTA
                              ** ********* **** *******************************  700

FBXW7_HAPLOTYPE-B            ATGCTGAGAGGTTGATGGCTCTGGATGAGTTGATTACATTCTGTGATCCC
FBXW7_TROSSULUS              ATGCTGAGAGGTTGATGGCTCTGGATGAGTTGATTACATTCTGTGATCCC
FBXW7_EDULIS                 ATGCTGAGAGGTTGATGGCTCTGGATGAGTTGATTACATTCTGTGATCCC
                             **************************************************  750

FBXW7_HAPLOTYPE-B            ACACAGGTTCGTCACATGATGCAGGTCATTGAACCTCAGTTCCAGAGAGA
FBXW7_TROSSULUS              ACACAGGTTCGTCACATGATGCAGGTCATTGAACCTCAGTTCCAGAGAGA
FBXW7_EDULIS                 ACACAGGTTCGTCACATGATGCAGGTCATTGAACCTCAGTTCCAGAGAGA
                             **************************************************  800

FBXW7_HAPLOTYPE-B            TTTCATATCACTTTTGCCAAAAGAGCTTGCCCTTTATGTGCTGTCTTTCC
FBXW7_TROSSULUS              TTTCATATCACTTTTGCCAAAAGAGCTTGCCCTTTATGTGCTGTCTTTCC
FBXW7_EDULIS                 TTTTATATCACTTTTGCCAAAAGAACTTGCCCTTTATGTGCTGTCTTTCC
                             *** ******************** *************************  850

FBXW7_HAPLOTYPE-B            TGGAGCCAAAATCTTTACTGAAAGCTGCCCAGACGTGTCGATACTGGCGA
FBXW7_TROSSULUS              TGGAGCCAAAATCTTTACTGAAAGCTGCCCAGACGTGTCGATACTGGCGA
FBXW7_EDULIS                 TGGAGCCAAAATCTTTACTTAAAGCTGCCCAGACGTGTCGATACTGGCGA
                             ******************* ******************************  900

FBXW7_HAPLOTYPE-B            GTACTGGCAGAGGACAATTTACTCTGGAGAGAGAAATGTAGGGAAGAAAG
FBXW7_TROSSULUS              GTACTGGCAGAGGACAATTTACTCTGGAGAGAGAAATGTAGGGAAGAAAG
FBXW7_EDULIS                 GTACTGGCAGAGGACAATTTACTCTGGAGAGAGAAATGTCGCGAAGAAAG
                             *************************************** * ********  950

FBXW7_HAPLOTYPE-B            TATTGATGATAATTTAGTATATGGGAGTAATCGACTGAGACGACGTCATA
FBXW7_TROSSULUS              TATTGATGATAATTTAGTATATGGGAGTAATCGACTGAGACGACGTCATA
FBXW7_EDULIS                 TATTGATGATAATTTAGTGTATGGGATTAATCGACTCAGACGACGTCATA
                             ****************** ******* ********* ************* 1000

FBXW7_HAPLOTYPE-B            CCTCACGGTGTCCTTGGAAGTCATTGTACATGAGGCAGCATCAAATAGAA
FBXW7_TROSSULUS              CCTCACGGTGTCCTTGGAAGTCATTGTACATGAGGCAGCATCAAATAGAA
FBXW7_EDULIS                 CCTCACGGTGTCCTTGGAAGTCATTGTACATGAGGCAGCATCAAATAGAA
                             ************************************************** 1050

FBXW7_HAPLOTYPE-B            CTTAACTGGAGAACAGGAGAAATACCTGGTCCTAAGTTATTAAAAGGTCA
FBXW7_TROSSULUS              CTTAACTGGAGAACAGGAGAAATACCTGGTCCTAAGTTATTAAAAGGTCA
FBXW7_EDULIS                 CTCAACTGGAGAACAGGAGAAATACCTGGTCCTAAGTTGTTAAAAGGTCA
                             ** *********************************** *********** 1100

FBXW7_HAPLOTYPE-B            TGATGACCATGTGATAACCTGTTTAGAGTTTTGTGGTACTCGTATTGTCA
FBXW7_TROSSULUS              TGATGACCATGTGATAACCTGTTTAGAGTTTTGTGGTACTCGTATTGTCA
FBXW7_EDULIS                 TGATGACCATGTGATAACCTGTTTAGAGTTTTGTGGAACTCGTATTGTCA
                             ************************************ ************* 1150

FBXW7_HAPLOTYPE-B            GTGGGTCAGACGACAACACCCTCAAAGTATGGTCAGCCATCACAGGAAAG
FBXW7_TROSSULUS              GTGGGTCAGACGACAACACCCTCAAAGTATGGTCAGCCATCACAGGAAAG
FBXW7_EDULIS                 GTGGGTCAGATGACAACACCCTCAAAGTATGGTCAGCCATCACAGGAAAG
                             ********** *************************************** 1200


FBXW7_HAPLOTYPE-B            TGCTTAAGGACATTAGTTGGTCACACAGGAGGAGTGTGGTCATCCCAGAT
FBXW7_TROSSULUS              TGCTTAAGGACATTAGTTGGTCACACAGGAGGAGTGTGGTCATCCCAGAT
FBXW7_EDULIS                 TGCTTAAGGACATTAGTTGGTCACACAGGAGGGGTGTGGTCATCCCAGAT
                             ******************************** ***************** 1250

FBXW7_HAPLOTYPE-B            GTCTAACAGTGTGGTCATTAGTGGTTCTACTGACCGCACACTTAAAGTGT
FBXW7_TROSSULUS              GTCTAACAGTGTGGTCATTAGTGGTTCTACTGACCGCACACTTAAAGTGT
FBXW7_EDULIS                 GTCTAACAGTGTGGTCATCAGTGGTTCTACTGACCGCACACTTAAAGTGT
                             ****************** ******************************* 1300

FBXW7_HAPLOTYPE-B            GGAATGCCGACACAGGCCAGTGTATACATACATTGTACGGACATTCTTCT
FBXW7_TROSSULUS              GGAATGCCGACACAGGCCAGTGTATACATACATTGTACGGACATTCTTCT
FBXW7_EDULIS                 GGAATGCAGACACAGGCCAGTGTATACATACATTGTACGGACATTCTTCT
                             ******* ****************************************** 1350

FBXW7_HAPLOTYPE-B            ACAGTAAGATGTATGCATTTACATAAAAATGTGGTGGTGAGTGGCTCACG
FBXW7_TROSSULUS              ACAGTAAGATGTATGCATTTACATAAAAATGTGGTGGTGAGTGGCTCACG
FBXW7_EDULIS                 ACAGTAAGATGTATGCATTTACATAAAAATGTGGTGGTGAGTGGATCACG
                             ******************************************** ***** 1400

FBXW7_HAPLOTYPE-B            AGATGCTACATTACGGGTATGGGATATTGATAGTGGTGCTTGCTTACATG
FBXW7_TROSSULUS              AGATGCTACATTACGGGTATGGGATATTGATAGTGGTGCTTGTTTACATG
FBXW7_EDULIS                 AGATGCTACATTACGGGTATGGGATATTGATAGTGGTGCTTGTTTACATG
                             ****************************************** ******* 1450

FBXW7_HAPLOTYPE-B            TATTGATGGGACATGTGGCAGCAGTGAGATGTGTTCAGTATGACGGACGG
FBXW7_TROSSULUS              TATTGATGGGACATGTGGCAGCAGTGAGATGTGTTCAGTATGACGGACGG
FBXW7_EDULIS                 TATTGATGGGACATGTGGCAGCAGTGAGATGTGTTCAGTACGATGGAAGG
                             **************************************** ** *** ** 1500

FBXW7_HAPLOTYPE-B            AGGGTGGTCAGTGGAGCCTATGATTATATGGTGAAAGTCTGGGATCCAGA
FBXW7_TROSSULUS              AGGGTGGTCAGTGGAGCCTATGATTATATGGTGAAAGTCTGGGATCCAGA
FBXW7_EDULIS                 AGGGTGGTCAGTGGAGCCTATGATTACATGGTGAAAGTCTGGGATCCAGA
                             ************************** *********************** 1550

FBXW7_HAPLOTYPE-B            AACAGAGACATGTTTACATACACTTCAGGGACATACAAATAGGGTTTACT
FBXW7_TROSSULUS              AACAGAGACATGTTTACATACACTTCAGGGACATACAAATAGGGTTTACT
FBXW7_EDULIS                 AACAGAAACATGTTTACATACACTTCAGGGACATACAAATAGGGTTTACT
                             ****** ******************************************* 1600

FBXW7_HAPLOTYPE-B            CGCTACAGTTTGATGGGATTCACATTGTGAGTGGATCTCTGGACACATCT
FBXW7_TROSSULUS              CGCTACAGTTTGATGGGATTCACATTGTGAGTGGATCTCTGGACACATCT
FBXW7_EDULIS                 CGCTACAGTTTGATGGGATTCACATTGTGAGTGGATCTCTGGACACATCT
                             ************************************************** 1650

FBXW7_HAPLOTYPE-B            ATCCGGGTTTGGGATGTAGAAGCTGGCAACTGTTTACATACATTAATTGG
FBXW7_TROSSULUS              ATCCGGGTTTGGGATGTAGAAGCTGGCAACTGTTTACATACATTAATTGG
FBXW7_EDULIS                 ATCCGGGTTTGGGATGTAGAAGCTGGCAACTGTTTACATACATTAATTGG
                             ************************************************** 1700

FBXW7_HAPLOTYPE-B            TCATCAGTCCTTGACCAGTGGTTTGGAACTGAAGGACAATATCCTGGTTT
FBXW7_TROSSULUS              TCATCAGTCCTTGACCAGTGGTTTGGAACTGAAGGACAATATCCTGGTTT
FBXW7_EDULIS                 TCATCAGTCCTTGACCAGTGGTCTGGAACTGAAGGACAATATCCTGGTTT
                             ********************** *************************** 1750

FBXW7_HAPLOTYPE-B            CTGGTAACGCAGATTCTACTGTAAAAGTATGGGATATAACTACAGGCCAG
FBXW7_TROSSULUS              CTGGTAACGCAGATTCTACTGTAAAAGTATGGGATATAACTACAGGCCAG
FBXW7_EDULIS                 CTGGTAATGCAGATTCTACTGTCAAAGTATGGGATATAACTACAGGCCAG
                             ******* ************** *************************** 1800

FBXW7_HAPLOTYPE-B            TGTCTTCAAACCTTACAAGGACCCAATAAACATTCTAGTGCTGTAACCTG
FBXW7_TROSSULUS              TGTCTTCAAACCTTACAAGGACCCAATAAACATTCTAGTGCTGTAACCTG
FBXW7_EDULIS                 TGTCTTCAAACCTTACAAGGACCCAATAAACATTCTAGTGCTGTGACCTG
                             ******************************************** ***** 1850

FBXW7_HAPLOTYPE-B            TTTACAGTTCAACAAAAAGTTTGTGATCACATCCTCAGACGATGGAACAG
FBXW7_TROSSULUS              TTTACAGTTCAACAAAAAGTTTGTGATCACATCCTCAGACGATGGAACAG
FBXW7_EDULIS                 TTTACAGTTCAACAAAAAGTTTGTGATCACATCCTCAGATGATGGAACAG
                             *************************************** ********** 1900

FBXW7_HAPLOTYPE-B            TCAAAATATGGGACTTAAAAACTGGTGATTTCATCCGTAATCTTGTGGCA
FBXW7_TROSSULUS              TCAAAATATGGGACTTAAAAACTGGTGATTTCATCCGTAATCTTGTGGCA
FBXW7_EDULIS                 TCAAAATATGGGACTTAAAAACTGGTGACTTCATCCGGAATCTTGTGGCA
                             **************************** ******** ************ 1950


FBXW7_HAPLOTYPE-B            TTAGACAGTGGGGGTAGTGGAGGGGTCGTGTGGAGAGTTAGATGTAGTAA
FBXW7_TROSSULUS              TTAGACAGTGGGGGTAGTGGAGGGGTCGTGTGGAGAGTTAGATGTAGTAA
FBXW7_EDULIS                 TTAGACAGTGGGGGTAGTGGAGGGGTCGTGTGGAGAGTTCGATGTAGTAA
                             *************************************** ********** 2000

FBXW7_HAPLOTYPE-B            CACTAAACTTGTCTGTGCTGTTGGTAGCAGAAATGGCACTGAAGAAACAA
FBXW7_TROSSULUS              CACTAAACTTGTCTGTGCTGTTGGTAGCAGAAATGGCACTGAAGAAACAA
FBXW7_EDULIS                 CACTAAACTTGTCTGTGCTGTTGGTAGTAGAAATGGCACTGAAGAAACAA
                             *************************** ********************** 2050

FBXW7_HAPLOTYPE-B            AACTTCATGTACTCGACTTTGATGTGGATGAAAAGAAATGTTTATACTGA
FBXW7_TROSSULUS              AACTTCATGTACTCGACTTTGATGTGGATGAAAAGAAATGTTTATACTGA
FBXW7_EDULIS                 AACTTCATGTACTCGACTTTGATGTGGATGAAAAGAAATGTTTATACTGA
                             ************************************************** 2100
```
