## Supplementary File S2 for "Transcriptomics of mussel transmissible cancer MtrBTN2 suggests accumulation of multiple cancerous traits and oncogenic pathways shared among bilaterians"

```
ALIGNMENTS PIK3CA-GENE_AMINO ACID SEQUENCES


PIK3CA_EDULIS                       MPPSSGELWGHHLMPSTIQVDCLLPTGIIIQVAVSRDETLERIKADLWVK
PIK3CA_TROSSULUS                    MPPSSGELWGHHLMPSTIQVDCLLPTGIIIQVAVSRDETLERIKADLWVK
PIK3CA_ CANCER                      MPPSSGELWGHHLMPSTIQVDCLLPTGIIIQVAVSRDETLERIKADLWVK
                                    **************************************************

PIK3CA_EDULIS                       AKLYPLYERLLEPASYIFVSITQDARKEEFYDETRRFCDLRLFQPILKIV
PIK3CA_TROSSULUS                    AKLYPLYERLLEPASYIFVSITQDARKEEFYDETRRFCDLRLFQPILKIV
PIK3CA_ CANCER                      AKLYPLYERLLEPASYIFVSITQDARKEEFYDETRRFCDLRLFQPILKIV
                                    **************************************************

PIK3CA_EDULIS                       EPVGNREEKMLNFEIGVTIGVSINEFNEMKDLEVMTFRRNILDICKNAID
PIK3CA_TROSSULUS                    EPVGNREEKMLNFEIGVTIGVSINEFNEMKDLEVMTFRRNILDICKNAID
PIK3CA_ CANCER                      EPVGNREEKMLNFEIGVTIGVSINEFNEMKDLEVMTFRRNILDICKNAID
                                    **************************************************

PIK3CA_EDULIS                       ERQRRGKASEALYAYPPDIESSPILPSHLQTKIKEMKNDVVACVWVVSED
PIK3CA_TROSSULUS                    ERQRRGKASEALYAYPPDIESSPILPSHLQTKIKEMKNDVVACVWVVSED
PIK3CA_ CANCER                      ERQRRGKASEALYAYPPDIESSPILPSHLQTKIKEMKNDVVACVWVVSED
                                    **************************************************

PIK3CA_EDULIS                       NSRSKFSVKVSHDAYPIDVIAGTIRRRSRMMGISKEHAERCIEEYSDTYA
PIK3CA_TROSSULUS                    NSRSKFSVKVSHDAYPIDVIAGTIRRRSRMMGISKEHAERCIEEYSDTYA
PIK3CA_ CANCER                      NSRSKFSVKVSHDAYPIDVIAGTIRRRSRMMGISKEHAERCIEEYSDTYA
                                    **************************************************

PIK3CA_EDULIS                       LKVCGCDQFLLEEHPLSQYKYVRECIARDNIPQFMLLTKESLYAAISPNI
PIK3CA_TROSSULUS                    LKVCGCDQFLLEEHPLSQYKYVRECIARDNIPQFMLLTKESLYAAISPNI
PIK3CA_ CANCER                      LKVCGCDQFLLEEHPLSQYKYVRECIARDNIPQFMLLTKESLYAAISPNI
                                    **************************************************

PIK3CA_EDULIS                       FITPSYTQKGMQLLTEINQQKTLSLWEIHAKLRIKILCATYVNVKELGKI
PIK3CA_TROSSULUS                    FITPSYTQKGMQLLTEINQQKTLSLWEIHAKLRIKILCATYVNVKELGKI
PIK3CA_ CANCER                      FITPSYTQKGMQLLTEINQQKTLSLWEIHAKLRIKILCATYVNVKELGKI
                                    **************************************************

PIK3CA_EDULIS                       YVKAGIYHGTEALCEFQDTKMVDSNNPQWHEWLEFLYIQDLPRSAKLCLS
PIK3CA_TROSSULUS                    YVKAGIYHGTEALCEFQDTKMVDSNNPQWHEWLEFLYIQDLPRSAKLCLS
PIK3CA_ CANCER                      YVKAGIYHGTEALCEFQDTKMVDSNNPQWHEWLEFLYIQDLPRSAKLCLS
                                    **************************************************


PIK3CA_EDULIS                       ICYTSKKKREPLSLGWANLQLFDFNDRLMNEKVSLNLWPLPQGMDDLLNY
PIK3CA_TROSSULUS                    ICYTSKKKREPLSLGWANLQLFDFNDRLMNEKVSLNLWPLPQGMDDLLNY
PIK3CA_ CANCER                      ICYTSKKKREPLSLGWANLQLFDFNDRLMNEKVSLNLWPLPQGMDDLLNY
                                    **************************************************

PIK3CA_EDULIS                       VGLPGSNPDQETPCLEIEFDRLSHPVSYPPEKQIEDLAQYAISKEQPLIY
PIK3CA_TROSSULUS                    VGLPGSNPDQETPCLEIEFDRLSHPVSYPPEKQIEDLAQYAISKESPLIY
PIK3CA_ CANCER                      VGLPGSNPDQETPCLEIEFDRLSHPVSYPPEKQIEDLAQYAISKESPLIY
                                    *********************************************.****

PIK3CA_EDULIS                       LDTPQIQAKEQAMVDDITSRDPLSEISEQEKDILWKLREYCIQVPQSLPK
PIK3CA_TROSSULUS                    LDTPQIQAKEQAMVDDITSRDPLSEISEQEKDILWKLREYCIRVPQSLPK
PIK3CA_ CANCER                      LDTPQIQAKEQAMVDDITSRDPLSEISEQEKDILWKLREYCIRVPQSLPK
                                    ******************************************:*******

PIK3CA_EDULIS                       LLQSVKWNEREYVAQLYMLLRRWPRLIPEFAMELLDCSYPDLCVRQYAVT
PIK3CA_TROSSULUS                    LLQSVKWNEREYVAQLYMLLRRWPRLMPEFAMELLDCSYPDLCVRQYAVT
PIK3CA_ CANCER                      LLQSVKWNEREYVAQLYMLLRRWPRLMPEFAMELLDCSYPDLCVRQYAVT
                                    **************************:***********************

PIK3CA_EDULIS                       CLDHGFSDDKLQQYMLQLVQALKFEPYLDSPITRFLLKRALQNQKTGQLF
PIK3CA_TROSSULUS                    CLDHGFSDDKLQQYMLQLVQALKFEPYLDSPITRFLLKRALQNQKTGQLF
PIK3CA_ CANCER                      CLDHGFSDDKLQQYMLQLVQALKFEPYLDSPITRFLLKRALQNQKTGQLF
                                    **************************************************

PIK3CA_EDULIS                       FWHLKSEIHNTSIQLRFGLVLEAFCRGCGSNLKMLLRQVEALDKLTKLTN
PIK3CA_TROSSULUS                    FWHLKSEIHNTSIQLRFGLVLEAFCRGCGSNLKMLLRQVEALDKLTKLTN
PIK3CA_ CANCER                      FWHLKSEIHNTSIQLRFGLVLEAFCRGCGSNLKMLLRQVEALDKLTKLTN
                                    **************************************************


PIK3CA_EDULIS                       MIKNEVKDDINEIMKFLANQLQQPDYQDGLKNFLSPLDNSHILGDLDISH
PIK3CA_TROSSULUS                    MIKNEVKDDINEIMKFLANQLQQPDYQDGLKNFLSPLDNSHILGDLDISH
PIK3CA_ CANCER                      MIKNEVKDDINEIMKFLANQLQQPDYQDGLKNFLSPLDNSHILGDLDISH
                                    **************************************************

PIK3CA_EDULIS                       CSVMTSKKKPLWLVFSNPDVMADIWFTDYKLIFKNGDDLRQDMLTVQLFK
PIK3CA_TROSSULUS                    CSVMTSKKKPLWLVFSNPDVMADIWFTDYKLIFKNGDDLRQDMLTVQLFK
PIK3CA_ CANCER                      CSVMTSKKKPLWLVFSNPDVMADIWFTDYKLIFKNGDDLRQDMLTVQLFK
                                    **************************************************

PIK3CA_EDULIS                       IMDTLWKNEGLDLRLIPYSVVSTGKDVGVIEIVRDSSTIMSIQQKNGIRA
PIK3CA_TROSSULUS                    IMDTLWKNEGLDLRLIPYSVVSTGKDVGVIEIVRDSSTIMSIQQKNGIRA
PIK3CA_ CANCER                      IMDTLWKNEGLDLRLIPYSVVSTGKDVGVIEIVRDSSTIMSIQQKNGIRA
                                    **************************************************

PIK3CA_EDULIS                       AVQMDSLGLYNWIMYHNKDREEQAISNFTRSCAGYCVATFILGIKDRHSG
PIK3CA_TROSSULUS                    AVQMDSLGLYNWIMYHNKDREEQAISNFTRSCAGYCVATFILGIKDRHSG
PIK3CA_ CANCER                      AVQMDSLGLYNWIMYHNKDREEQAISNFTRSCAGYCVATFILGIKDRHSG
                                    **************************************************

PIK3CA_EDULIS                       NIMVRKNGQVFHIDFGHFLDHRKKKFGITRERVPFVLTMDFIRVIARGSD
PIK3CA_TROSSULUS                    NIMVRKNGQVFHIDFGHFLDHRKKKFGITRERVPFVLTMDFIRVIARGSD
PIK3CA_ CANCER                      NIMVRKNGQVFHIDFGHFLDHRKKKFGITRERVPFVLTMDFIRVIARGSD
                                    **************************************************

PIK3CA_EDULIS                       QPLKHKEFKKFQQLCCEAYLIIRKNAYLFINLMTMMLSCGIPELQSLDDI
PIK3CA_TROSSULUS                    QPLKHKEFKKFQQLCCEAYLIIRKNAYLFINLMTMMLSCGIPELQSLDDI
PIK3CA_ CANCER                      QPLKHKEFKKFQQLCCEAYLIIRKNAYLFINLMTMMLSCGIPELQSLDDI
                                    **************************************************

PIK3CA_EDULIS                       SYLRKTLAVEEKDDEKALKYFIAKFNSAYSDAWTVKTDWLFHYMKNR
PIK3CA_TROSSULUS                    SYLRKTLAVEEKDDEKALKYFIAKFNSAYSDAWTVKTDWLFHYMKNR
PIK3CA_ CANCER                      SYLRKTLAVEEKDDEKALKYFIAKFNSAYSDAWTVKTDWLFHYMKNR
                                    ***********************************************


ALIGNMENTS PIK3CB-GENE_AMINO ACID SEQUENCES

PIK3CB_TROSSULUS      MPPVIVSPDLDVTSFQNDLGQLDIDFLLPNGICVPLQVDIDRPLDQIKQELWKQAANYPL
PIK3CB_CANCER         MPPVIVSPDLDVTSFQNDLGQLDIDFLLPNGICVPLQVDIDRPLDQIKQELWKQAANYPL
PIK3CB_EDULIS         MPPVIVSPDLDVTSFQNDLGQLDIDFLLPNGICVPLQVDIDRPLDQIKQELWKQAANYPL
                      ************************************************************

PIK3CB_TROSSULUS      FQQLATFDKYGFLYISGEGQKEDVMDESVGIAELRLAASFLKVVEKPADDKRKTIDRQIN
PIK3CB_CANCER         FQQLATFDKYGFLYISGEGQKEDVMDESVGIAELRLAASFLKVVEKPADDKRKTIDRQIN
PIK3CB_EDULIS         FQQLATFDKYGFLYISGEGQKEDVMDESVGISELRLAASFLKVVEKPADDKRKTIDRQIN
                      *******************************:****************************

PIK3CB_TROSSULUS      SLIGMKGSNYIQAQHSVSKAEISEFRQKMTTYCNNDVLKQANQDPTGLVYFEYKHPCRLR
PIK3CB_CANCER         SLIGMKGSNYIQAQHSVSKAEISEFRQKMTTYCNNDVLKQANQDPTGLVYFEYKHPCRLR
PIK3CB_EDULIS         SLIGMKGSNYIQAQHSVSKAEISEFRQKMTTYCNNDVLKQANQDPTGLVYFEYKHPCRLR
                      ************************************************************

PIK3CB_TROSSULUS      STSELPPDVENMVIDNIVLTVIVQETGTSYKLRVKKVSTPDELVKMTVDKWSSKSGQKTI
PIK3CB_CANCER         STSELPPDVENMVIDNIVLTVIVQETGTSYKLRVKKVSTPDELVKMTVDKWSSKSGQKTI
PIK3CB_EDULIS         STSELPPDVENMVIDYIVLTVIVQETGTSYKLRVKKLSTPDELVKMTVDKWSSKSGQKTI
                      *************** ********************:***********************
PIK3CB_TROSSULUS      NCNDYVLQVLGMNDFLYGDNPLVMFKYVYSCIVKNTAPEFWLKNKSSVLEKQEVSQPHKS
PIK3CB_CANCER         NCNDYVLQVLGMNDFLYGDNPLVMFKYVYSCIVKNTAPEFWLKNKSSVLEKQEVSQPHKS
PIK3CB_EDULIS         NCNDYVLQVLGMNDFLYGDNPLVMFKYVYSCIVKNTAPEFWLKNKSSVLEKQEVSQPHKS
                      ************************************************************

PIK3CB_TROSSULUS      SRTSFQEERRNTVRSEKRSGHCVWDIHNNFYITLHTACNVQVPENCRNLVRLCVGVFHGP
PIK3CB_CANCER         SRTSFQEERRNTVRSEKRSGHCVWDIHNNFYITLHTACNVQVPENCRNLVRLCVGVFHGP
PIK3CB_EDULIS         SRTSFQEERRNTVRSEKRSGHCVWDIHQNFYITLHTACNVQVPENCRNLVRLCVGVFHGP
                      ***************************:********************************

PIK3CB_TROSSULUS      DPLCSIQETKEVFVSPDGLCSFNDCINFDIKVQDLPQMSRLCFGLHCKKGKEPLLLAWAN
PIK3CB_CANCER         DPLCSIQETKEVFVSPDGLCSFNDCINFDIKVQDLPQMSRLCFGLHCKKGKEPLLLAWAN
PIK3CB_EDULIS         DPLCSIQETKEVFVSQDGLCSFNDCINFDIKVQDLPQMSRLCFGLHCKKGKEPLLLAWAN
                      *************** ********************************************

PIK3CB_TROSSULUS      IPIFDYKSNLQKGKLKLPLWPRSELIQQEESNCFPVGTVATNPQGDCSMIEFSVPDFTKK
PIK3CB_CANCER         IPIFDYKSNLQKGKLKLPLWPRSELIQQEESNCFPVGTVATNPQGDCSMIEFSVPDFTKK
PIK3CB_EDULIS         IPIFDYKSNLQKGKLKLPLWPRSELIQQEESYCFPVGTVATNPQGDCSMIEFSVPDFTKK
                      ******************************* ****************************

PIK3CB_TROSSULUS      GSIYYPPIEKVLQCASDHMEEPGKCGSPTWRPNKNIVQQLDTILRANQYTVLQTLDEQQK
PIK3CB_CANCER         GSIYYPPIEKVLQCASDHMEEPGKCGSPTWRPNKNIVQQLDTILRANQYTVLQTLDEQQK
PIK3CB_EDULIS         GSIYYPPIEKVLQCASDHMEEPGKCGSPTWRPNKNIVQQLDSILRANQYTILQTLDEQQK
                      *****************************************:********:*********

PIK3CB_TROSSULUS      QLIWWMRYDIRDKCSEFPHALLYVLYSVSWDNHIEVSKMQALLQTWPTLEADQALTLLDF
PIK3CB_CANCER         QLIWWMRYDIRDKCSEFPHALLYVLYSVSWDNHIEVSKMQALLQTWPTLEADQALTLLDF
PIK3CB_EDULIS         QLIWWMRYDIRDKCSEFPHALLYVLYSVSWDNHIEVSKMQALLQTWPTLEADQALTLLDF
                      ************************************************************

PIK3CB_TROSSULUS      TFPDKYVRKKATEWLDELPDEELAQYLLQLVQALKYENYLNCDLVKFLLRRALQNRNIGH
PIK3CB_CANCER         TFPDKYVRKKATEWLDELPDEELAQYLLQLVQALKYENYLNCDLVKFLLRRALQNRNIGH
PIK3CB_EDULIS         TFPDKYVRKKATEWLDELPDEELAQYLLQLVQALKYENYLNCDLVKFLLRRALQNRNIGH
                      ************************************************************

PIK3CB_TROSSULUS      KLFWLLKSDMHEPSVTVQYGLIIEIYLKANPSHMSILDRQQIILQKLKSITDIHTKAANL
PIK3CB_CANCER         KLFWLLKSDMHEPSVTVQYGLIIEIYLKANPSHMSILDRQQIILQKLKSITDIHTKAANL
PIK3CB_EDULIS         KLFWLLKSDMHEPSVTVQYGLIIEIYLKANPSHMSILDRQQIILQKLKSITDIHTKAANL
                      ************************************************************

PIK3CB_TROSSULUS      KKKAKEKDDSPLQKVMQQSAYKEAFCQIYSPITLMYKLDKIVEKSCKVMDSKKKPFWLEW
PIK3CB_CANCER         KKKAKEKDDSPLQKVMQQSAYKEAFCQIYSPITLMYKLDKIVEKSCKVMDSKKKPFWLEW
PIK3CB_EDULIS         KKKAKEKDDSPLQKVMQQSAYKEAFCKIYSPITLMYKLDKIVEKSCKVMDSKKKPFWLEW
                      **************************:*********************************

PIK3CB_TROSSULUS      TNDDEKGQNIQLIYKCGDDLRQDMLTLQILEVMDTIWQSQGYDLRLNPYGCVATGCEEGM
PIK3CB_CANCER         TNDDEKGQNIQLIYKCGDDLRQDMLTLQILEVMDTIWQSQGYDLRLNPYGCVATGCEEGM
PIK3CB_EDULIS         TNDDEKGQNIQLIYKCGDDLRQDMLTLQILEVMDTIWQSEGYDLRLNPYGCVATGCEEGM
                      ***************************************:********************

PIK3CB_TROSSULUS      IEVVQKSLTLAGIQKWRKLGLDKRSLYDWLKHKNPTEHSLQRAVEEFKLSCAGYAVATYI
PIK3CB_CANCER         IEVVQKSLTLAGIQKWRKLGLDKRSLYDWLKHKNPTEHSLQRAVEEFKLSCAGYAVATYI
PIK3CB_EDULIS         IEVVQKSLTLAGIQKWRKLGLDKRSLYDWLKHKNPTEHSLQRAVEEFKLSCAGYAVATYI
                      ************************************************************

PIK3CB_TROSSULUS      LGVGDRHNDNIMMKESGQLFHIDFGHFLGNKKTKFNINRERVPFILTSHFEYIIKDGDQK
PIK3CB_CANCER         LGVGDRHNDNIMMKESGQLFHIDFGHFLGNKKTKFNINRERVPFILTSHFEYIIKDGDQK
PIK3CB_EDULIS         LGVGDRHNDNIMMKETGQLFHIDFGHFLGNKKTKFNINRERVPFILTSHFEYIIKDGEKK
                      ***************:*****************************************::*

PIK3CB_TROSSULUS      PQNFTDFKDICERAYLIIRSKAHLLIQLFTMMLSSGIPQLNNVSDIDYIKDVLALNSTEE
PIK3CB_CANCER         PQNFTDFKDICERAYLIIRSKAHLLIQLFTMMLSSGIPQLNNVSDIDYIKDVLALNSTEE
PIK3CB_EDULIS         PQNFTHFKEICERAYLIIRSRAHLLIQLFMMMLSSGIPQLNNVSDIDYIKEVLALNSTQE
                      *****.**:***********:******** ********************:*******:*

PIK3CB_TROSSULUS      VALDKFRKKFKEAQDSSWSTTVNWWFHMRVH
PIK3CB_CANCER         VALDKFRKKFKEAQDSSWSTTVNWWFHMRVH
PIK3CB_EDULIS         EALEKFRKKFKEAQESSWSTTVNWWFHMRVH
                       **:**********:****************


ALIGNMENTS PTEN-GENE_AMINO ACID SEQUENCES

PTEN_TROSSULUS      MDFLKGLVSKNKRRHKVDGFDLDLTYIYPNIIAMGFPAEKLEGVYRNHIDDVIEFLDTKH
PTEN_CANCER         MDFLKGLVSKNKRRHKVDGFDLDLTYIYPNIIAMGFPAEKLEGVYRNHIDDVIEFLDTKH
PTEN_EDULIS         MDFLKGLVSKNKRRHKVDGFDLNLTYIYPNIIAMGFPAEKLEGVYRNHIDDVIEFLDTKH
                    **********************:*************************************

PTEN_TROSSULUS      KDHYKVYNLCTERSYDPDRFHGRVVAYPFEDHNPPRLELIKPFCEDLDEWLKKSNENIAA
PTEN_CANCER         KDHYKVYNLCTERSYDPDRFHGRVVAYPFEDHNPPRLELIKPFCEDLDEWLKKSNENIAA
PTEN_EDULIS         KDHYKVYNLCTERSYDPDRFHGRVVAYPFEDHNPPRLELIKPFCEDLDEWLKKSDENIAA
                    ******************************************************:*****

PTEN_TROSSULUS      IHCKAGKGRTGVMICAYMLHRNKFDNSKEALRFYGQARTQDEKGVTIPSQRRYVEYYEYL
PTEN_CANCER         IHCKAGKGRTGVMICAYMLHRNKFDNSKEALRFYGQARTQDEKGVTIPSQRRYVEYYEYL
PTEN_EDULIS         IHCKAGKGRTGVMICAYMLHRNKFDNTKEALRFYGQARTQDEKGVTIPSQRRYVEYYEYL
                    **************************:*********************************

PTEN_TROSSULUS      IRNKLNYKPVALLLKGIEFITVPMHNGSGCSPFFEVYQLKVRVYTSKVYEDIKKGQTSFY
PTEN_CANCER         IRNKLNYKPVALLLKGIEFITVPMHNGSGCSPFFEVYQLKVRVYTSKVYEDIKKGQTSFY
PTEN_EDULIS         IRNKLNYKPVALLLKGIEFITVPMHNGSGCSPFFEVYQLKVRVYTSKVYEDIKKGQTSFY
                    ************************************************************


PTEN_TROSSULUS      MPIEQSVPLCGDIKVVFYNKPRMKKKDKMFQFWLNTFFVEADDKPRENGRKSWAGEVPGS
PTEN_CANCER         MPIEQSVPLCGDIKVVFYNKPRMKKKDKMFQFWLNTFFVEADDKPRENGRKSWAGEVPGS
PTEN_EDULIS         MPIEQSVPLCGDIKVVFYNKPRMKKKDKMFQFWLNTFFVEADDKPKENGRKSWAGEVPGS
                    *********************************************:**************

PTEN_TROSSULUS      SNYYTVTIPKCELDKANKDKAHKLFSPNFQVKLHFTNPENCGLYNSSLKVPDQQGLRHRS
PTEN_CANCER         SNYYTVTIPKCELDKANKDKAHKLFSPNFQVKLHFTNPENCGLYNSSLKVPDQQGLRHRS
PTEN_EDULIS         SNYYTVTIPKCELDKANKDKAHKLFSPNFQVKLHFTNPENCGLYNSSLKVPDQQGLRHRS
                    ************************************************************

PTEN_TROSSULUS      KTLEQICDKNSTTNHSDHSVDGMPHSITALTLNSRRENQSPVITRTRSPSHMTHERPKFN
PTEN_CANCER         KTLEQICDKNSTTNHSDHSVDGMPHSITALTLNSRRENQSPVITRTRSPSHMTHERPKFN
PTEN_EDULIS         KTLEQICDKNSTTNHSDHSVDGMPHSITALTLNSRRENQSPVITRTRSPSHMTHERPKFM
                    ***********************************************************

PTEN_TROSSULUS      LPSNGEGSSDQLSSEANSDTDFPEEDLSDTDDEEEWNDLETTAV
PTEN_CANCER         LPSNGEGSSDQLSSEANSDTDFPEEDLSDTDDEEEWNDLETTAV
PTEN_EDULIS         LPSNGEGSSDQLSSEANSDTDFPEEDLSDTDDEEEWNDLETTAV
                    ********************************************


ALIGNMENTS AKT-GENE_AMINO ACID SEQUENCES (red box: pleckstrin homology domain, green box: protein-kinase domain, blue box: C-terminal domain)
```

```
AKT_EDULIS         MNPSGSSIPVVKEGYLMKRGEHIKNWRKRYFILRKDGSFLGFRAKPEQGFNDPLNDFTVR
AKT_TROSSULUS      MNPSGSSIPVVKEGYLMKRGEHIKNWRKRYFILRKDGSFLGFRAKPEQGFNDPLNDFTVR
AKT_CANCER         MNPSGSSIPVVKEGYLMKRGEHIKNWRKRYFILRKDGSFLGFRAKPEQGFNDPLNDFTVR
                    ************************************************************
```

```
AKT_EDULIS         DCQLMRQERPKANTFIIRGLQWTTVVERMFCVESQEEREEWIAAINSISDELKNSDEGAE
AKT_TROSSULUS      DCQLMRQERPKANTFIIRGLQWTTVVERMFCVESQEEREEWIAAINSISDELKNSDEGAE
AKT_CANCER         DCQLMRQERPKANTFIIRGLQWTTVVERMFCVESQEEREEWIAAINSISDELKNSDEGAE
                    ************************************************************
```

```
AKT_EDULIS         GPSLFDKSKKKVSLDDFEFLKVLGKGTFGKVILCKEKVTGHLLAIKILKKQVIIQKDEVA
AKT_TROSSULUS      GPSLFDKSKKKVSLDDFEFLKVLGKGTFGKVILCKEKVTGHLLAIKILKKQVIIQKDEVA
```

AKT_CANCER	GPSLFDKSKKKVSLDDFEFLKVLGKGTFGKVILCKEKLTGHLLAIKILKKQVIIQKDEVA

*************************************:**********************

```
AKT_EDULIS         HTLTENRVLQTTKHPFLTQLKYSFQTTDRLCFVMEYVNGGELFFHLSRERVFSEERTKFY
AKT_TROSSULUS      HTLTENRVLQTTKHPFLTQLKYSFQTTDRLCFVMEYVNGGELFFHLSRERVFSEERTKFY
AKT_CANCER         HTLTENRVLQTTKHPFLTQLKYSFQTTDRLCFVMEYVNGGELFFHLSRERVFSEERTKFY
                    ************************************************************
```

```
AKT_EDULIS         GAEIISALGYLHENNIVYRDLKLENLLLDKDGHIKIADFGLCKEEMFYGASTKTFCGTPE
AKT_TROSSULUS      GAEIISALGYLHENNIVYRDLKLENLLLDKDGHIKIADFGLCKEEMFYGASTKTFCGTPE
AKT_CANCER         GAEIISALGYLHENNIVYRDLKLENLLLDKDGHIKIADFGLCKEEMFYGASTKTFCGTPE
                    ************************************************************
```

```
AKT_EDULIS         YLAPEVLEDNDYGRAVDWWGTGVVMYEMMCGRLPFYNRDHDVLFELILLQSVRFPRTLSE
AKT_TROSSULUS      YLAPEVLEDNDYGRAVDWWGTGVVMYEMMCGRLPFYNRDHDVLFELILLQSVRFPRTLSE
AKT_CANCER         YLAPEVLEDNDYGRAVDWWGTGVVMYEMMCGRLPFYNRDHDVLFELILLQSVRFPRTLSE
                    ************************************************************
```

```
AKT_EDULIS         DAKSLLEGFLKKNPHERLGGSEQDVKEIIAHPFFKTVNWQDLIEKKITPPWKPDVKNDYD
AKT_TROSSULUS      DAKSLLEGFLKKNPHERLGGSEQDVKEIIAHPFFKTVNWQDLIEKKITPPWKPDVKNDYD
AKT_CANCER         DAKSLLEGFLKKNPHERLGGSEQDVKEIIAHPFFKTVNWQDLIEKKITPPWKPDVKNDYD
                    ************************************************************
```

```
AKT_EDULIS         TKYIPDEFAQVPLSLTPNSKESHVELSSIAEESELPYFEQFSYHGSRAGMGESYLQSHVN
AKT_TROSSULUS      TKYIPDEFAQVPLSLTPNSKESHVELSSIAEESELPYFEQFSYHGSRAGMGESYLQSHVN
AKT_CANCER         TKYIPDEFAQVPLSLTPNSKESHVELSSIAEESELPYFEQFSYHGSRAGMGESYLQSHVN
                    ************************************************************

AKT_EDULIS         DGEIF
AKT_TROSSULUS      DGEIF
AKT_CANCER         DGEIF
                    *****


ALIGNMENTS P53-GENE_AMINO ACID SEQUENCES(yellow box, transactivation domain; black box: proline-rich domain; blue box,DNA-binding region; red box,tetramerisation region; green box, SAM domain)
```

```
p53_TROSSULUS      MSQASVSTTCTPSGPPMSQETFEYLWNTLGEVTQEGGYTNITSKESIDYAFSEAEDETSI
p53_CANCER         MSQASVSTTCTPSGPPMSQETFEYLWNTLGEVTQEGGYTNITSKESIDYAFSEAEDETSI
p53_EDULIS         MSQASVSTTCTPSGPPMGQETFEYLWNTLGEVTQEGGYTNITSKESIDYAFSEAEDETSI
                   *****************.******************************************
```

```
p53_TROSSULUS      SVEKYRITSNDSISDLLNPIIGQTTSASSMSPDSQTNIIGSSASSPYNDTITSPPPYSPH
p53_CANCER         SVEKYRITSNDSISDLLNPIIGQTTSASSMSPDSQTNIIGSSASSPYNDTITSPPPYSPH
p53_EDULIS         SVEKYRITSNDSISDLLNPIIGQTTSASSMSPDSQTNIIGSSASSPYNDTITSPPPYSPH
                   ************************************************************
```

```
p53_TROSSULUS      TSMQSPIPSVPSNTDYPGDYGFTISFSQPSKETKSTTWTYSESLKKLYVRMATTCPIRFK
p53_CANCER         TSMQSPIPSVPSNTDYPGDYGFTISFSQPSKETKSTTWTYSESLKKLYVRMATTCPIRFK
p53_EDULIS         TSMQSPIPSVPSNTDYPGDYGFTISFSQPSKETKSTTWTYSESLKKLYVRMATTCPIRFK
```

************************************************************

```
p53_TROSSULUS      CLRQPPQGCVIRAMPIFMKPEHVQEPVKRCPNHATSKEHNENHPAPTHLCRCEHKLAKFV
p53_CANCER         CLRQPPQGCVIRAMPIFMKPEHVQEPVKRCPNHATSKEHNENHPAPTHLCRCEHKLAKFV
p53_EDULIS         CLRQPPQGCVIRAMPIFMKPEHVQEPVKRCPNHATSKEHNENHPAPTHLCRCEHKLAKFV
                   ************************************************************
```

```
p53_TROSSULUS      EDPYTSRQSVLIPHEIPQAGSEWVTNLFQFMCLGSCVGGPNRRPIQIVLTLEKDNQVLGR
p53_CANCER         EDPYTSRQSVLIPHEIPQAGSEWVTNLFQFMCLGSCVGGPNRRPIQIVLTLEKDNQVLGR
p53_EDULIS         EDPYTSRQSVLIPHEIPQAGSEWVTNLFQFMCLGSCVGGPNRRPIQIVLTLEKDNQVLGR
                   ************************************************************
```

```
p53_TROSSULUS      RAVEVRICACPGRDRKADEKAALPPCKQSPKKGQKVNIINEITTVTPGGKKRKAEDEPFT
p53_CANCER         RAVEVRICACPGRDRKADEKAALPPCKQSPKKGQKVNIINEITTVTPGGKKRKAEDEPFT
p53_EDULIS         RAVEVRICACPGRDRKADEKAALPPCKQSPKKGQKVNIINEITTVTPGGKKRKAEDEPFT
                   ************************************************************
```

```
p53_TROSSULUS      LSVRGRENYEILCRLRDSLELSSMVPQNQIDVYKQKQLDTNRQSSPTT--ARVVTLPTHD
p53_CANCER         LSVRGRENYEILCRLRDSLELSSMVPQNQIDVYKQKQLDTNRQSSPTT--ARVVTLPTHD
p53_EDULIS         LSVRGRENYEILCRLRDSLELSSMVPQNQIDVYKQKQLDTNRQSSPTTSTARVVTLPTHD
                   ************************************************  **********
```

```
p53_TROSSULUS      NTPITIQGEGRQTTLPFTADLNGQVTSSQNGVVENHGNIKEELMANGDHSISNWLTTLGL
p53_CANCER         NSPITIQGEGRQTTLPFTADLNGQVTSSQNGVVENHGNIKEELMANGDHSISNWLTTLGL
p53_EDULIS         NTPITIQGEGRQTTLPFTADLNGQVTSSQNGVVENHGNIKEELMANGDHSISNWLTTLGL
                   *:**********************************************************
```

```
p53_TROSSULUS      SAYIDNFHQQNLFTMEQLDDFTVEDLQKMRIGTSHRNKIWKALVEFHSESITISDSQSLQ
p53_CANCER         SAYIDNFHQQNLFTMEQLDDFTVEDLQKMRIGTSHRNKIWKALVEFHSESITISDSQSLQ
p53_EDULIS         SAYIDNFHQQNLFTMEQLDDFTVEDLQKMRIGTSHRNKIWKALVEFHSESITISDSQSLQ
                   ************************************************************

p53_TROSSULUS      RDVSTASTISMTSQASISQNSQNSTYCPGYYEVTRYTFKHTVSYTKVDERSPKRAKVD
p53_CANCER         RDVSTASTISMTSQASISQNSQNSTYCPGYYEVTRYTFKHTVSYTKVDERSPKRAKVD
p53_EDULIS         RDVSTASTISMTSQASISQNSQNSTYCPGYYEVTRYTFKHTVSYTKVDERSPKRAKVD
                   **********************************************************


ALIGNMENTS MDM2-GENE_AMINO ACID SEQUENCES (yellow box, SWIB/MDM2 domain; red box, RanBP2 region; green box, RING region with NoLS highlighted in green)
```

```
MDM2_TROSSULUS             MCNNPNLIVRQPPDTYSTPAGSNVSTLKQLTESHYITTEECEVNYPMVRP
MDM2_EDULIS                MCNNPNLIVRQPPDTYSTPAGSNVSTLKQLTESHYITTEECEVNYPMVRP
MDM2_CANCER                MCNNPNLIVRQPPDTYSSAAGSNVSTLEQLTESHYITTEECVVNYPMVRP
                           *****************:.********:************* ********
```

```
MDM2_TROSSULUS             RREFLDLLKQVGASWEVLSRKEVCSYLKDYIGSRQLYNPNDPSIVFCKKD
MDM2_EDULIS                RREFLDLLKQVGASWEVLSRKEVCSYLKDYIGSRQLYNPNDPSIVFCKKD
MDM2_CANCER                RREFLDLLKQVGASWEVLSRKEVCSYLKDYIGSRQLYNPNDPSIVFCKKD
                           **************************************************
```

```
MDM2_TROSSULUS             PLGKVFGVDKFTFTEATKLIDANCYQLPDSCMVKKEQLVAIPKKRSNSSN
MDM2_EDULIS                PLGKVFGVDKFTFTEATKLIDANCYQLPDSCMVKKEQLVAIPKKRSNSSN
MDM2_CANCER                PLGKVFGVDKFTFTEATKLIDANCYQLPDSCMVKKEQLVAIPKKRNNSSN
                           *********************************************.****

MDM2_TROSSULUS             QVNQIVTVSCPKLIDILVHSPSTQTGEQQKKQNSFEGQTSDTSSNRAQNL
MDM2_EDULIS                QVNQIVTVSCPKLIDILVHSPSTQTGEQQKKQNSLEGQTSDTSSNRAQNL
MDM2_CANCER                QVKQIVTVSCPKLIDILVHSPSTQTGEQQKKQNSFEGQTSDTSSNRAQNL
                           ********************************** ***************

MDM2_TROSSULUS             SAASTGRRRHRWVSERGDSEPDIIAIELDADNDDDAIGVEVGTNGTEDVM
MDM2_EDULIS                SAASTGRRRHRWVSERGDSEPDIIAIELDADNDDDAIGVEVGTNGTDDVM
MDM2_CANCER                SAASTGRRRHRWVSERGDSEPDIIAIELDAYNDDDAIGVEVETNGTEDVM
                           ****************************** ********** ****:***

MDM2_TROSSULUS             SVEVDADRISFEYEVCSDEDDIISSTASVSGISDTNVVIICGDSDMEFWG
MDM2_EDULIS                SVEVDADRISFEYEVCSDEDDIISSTASVSGISDTNVVIICGDSDMEFWG
MDM2_CANCER                SVEVDADRISFEYEVCSDEDDIISSTASVSGISDTNVVIICGDSDMEFWG
                           **************************************************
```

```
MDM2_TROSSULUS             DSDSSDSELSDGDKWTCTECDTKNSPLNGHCGHCKKVRPNWLPETTRKRR
MDM2_EDULIS                DSDSSDSELSDGDKWTCTECDTKNSPLNGHCGHCKKVRPNWLPETTRKRR
MDM2_CANCER                DSDSSDSELSDGDKWTCTECDTKNSPLNGHCGHCKKVRPNWLPETTRKRR
                           **************************************************

MDM2_TROSSULUS             LEKMNSKDKALLSSSECSDISHISSKIQRRSSRDRLTSIELESDKDLIAE
MDM2_EDULIS                LEKMNSKDKALLSSSECSDISHISSKIQRRSSRDRLTSIELESDKDLIAE
MDM2_CANCER                LEKMNSKDKALLSSSECSDISHISSKIQRRSSRDRLTSIELESDKDLIAE
                           **************************************************

MDM2_TROSSULUS             EEDSGISTLSRQQSSQENLAENKLKSYSSLIENSSTLGSFRTLLNFQTTA
MDM2_EDULIS                EEDSGISTLSRQQSSQENLAENKLKSYSSLIENSSTLGSFRTLLNFQTTA
MDM2_CANCER                EEDSGISTLSRQQSSQENLAENKLKSYSSLIENSSTLGSFRTLLNFQTTA
                           **************************************************
```

```
MDM2_TROSSULUS             SAKNKDEMTISNKIDKLDKPQKDLKLEKSSNIKEFENSQSQLGQCCICFS
MDM2_EDULIS                SAKNKDEMTISNKIDKLDKPQKDLKLEKSSNIKEFENSQSQLGQCCICFS
MDM2_CANCER                SAKNKDEGIKSNKNDKLDKPQKDLKLEKSSNIKEFENSQSQLGQCCICFS
```

******* **** ************************************

```
MDM2_TROSSULUS             RPKTASIIHGCTGHQVCCYRCAKRLKRLAKPCPLCRRPIQKVIKNYSA
MDM2_EDULIS                RPKTASIIHGCTGHQVCCYRCAKRLKRLAKPCPLCRRPIQKVIKNYSA
MDM2_CANCER                RPKTASIIHGCTGHQVCCYRCAKRLKRLAKPCPLCRRPIQKVIKNYSA
                           ************************************************


ALIGNMENTS MYC-GENE_AMINO ACID SEQUENCES (blue box, transcriptional activation domain; red box, DNA-binding region)
```

```
MYC_TROSSULUS          MMRKYSEIVIPRREGNSCQVVTEIGHTGIAQPPDNSVKMDKMCKHCYVDNSHVLQHCMYE
MYC_CANCER             MMRKYSEIVIPRREGNSCQVVTEIGHTGIAQPPDNSVKMDKMCKHCYVDNSHVLQHCMYE
MYC_EDULIS             MMRKYSEIVIPRREGNSCQVVTEIGHTGIAQPSDNSVKMAKMCKHCYVDNSHVLQHCMYE
                       ********************************.****** ********************
```

```
MYC_TROSSULUS          TESSSPASLPSDDIWKKFELIPTPPRSPEREEPFEMDLSLDLEDMNFPFDANFFESEDFN
MYC_CANCER             TEYSSPASLPSDDIWKKFELIPTPPRSPEREEPFEMDLSLDLEDMNFPFDANFFESEDFN
MYC_EDULIS             TESSSPASLPSDDIWKKFELIPTPPRSPEREEPFEMDLSLDLEDMNFPFDANFFESEDFN
                       ** *********************************************************
```

```
MYC_TROSSULUS          PKETFAESPPSQLCSKLIQDCMWSGDTFSSSEQKDKTSSSFDMKTVDPMAVFPCSVTNNT
MYC_CANCER             PKETFAESPLSQLCSKLIQDCMWSGDTFSSSEQKDKTSSSFDMKTVDPMAVFPCSVTNNT
MYC_EDULIS             PKETFAESPPSQLCSKLIQDCMWSGDTFSSSEQKDKTSSSFDMKTVDPMAVFPCSVTNNT
                       ********* **************************************************

MYC_TROSSULUS          NNSHSIMGYLSETKLHSLGTETPSDSEEEIDVVTVEKLQQNSTTAGPTINKSNIQTRNIK
MYC_CANCER             NNSHSIMGYLSETKLHSLGTETPSDSEEEIDVVTVEKLQQNSTTAGPTINKSNIQTRNIK
MYC_EDULIS             NNSHSIMGYLSETKLHSLGTETPSDSEEEIDVVTVEKLQQNSTTAGPTINKSNIQTRNIK
                       ************************************************************

MYC_TROSSULUS          TQIKIEKNNGEKVEIIPLKTTTLKIRVETPDLHNYSLPHSHQPKRVRSYPVSPSHSPLPQ
MYC_CANCER             TQIKIEKNNGEKVEIIPLKTTTLKIRVETPDLHNYSLPHSHQPKRVRSYPVSPSHSPLPQ
MYC_EDULIS             TQIKIEKNNGEKVEIIPLKTTTLKIRVETPDLHNYSLPHSHQPKRVRSYPVSPSHSPLPQ
                       ************************************************************
```

MYC_TROSSULUS	KRTKKDLSVPDFKRVCQKLRASKSSSDSEECISEGGKRTQHNVLERKRRNDLKYSFFTLR

```
MYC_CANCER             KRTKKDLSVPDFKRVCQKLRASKSSSDSEECISEGGKRTQHNVLERKRRNDLKYSFFTLR
MYC_EDULIS             KRTKKDLSVPDFKRVCQKLRASKSSSDSEECISEGGKRTQHNVLERKRRNDLKYSFFTLR
                       ************************************************************
```

MYC_TROSSULUS	DSVPELSNQERAPKVLILKKASDYVHSLNVDNKRLESEKATLLAKQQKLKRTLEILQDSD

```
MYC_CANCER             DSVPELSNQERAPKVLILKKASDYVHSLNVDNKRLESEKATLLAKQQKLKRTLEILQDSD
MYC_EDULIS             DSVPELSNQERAPKVLILKKASDYVHSLNVDNKRLESEKATLLAKQQKLKRTLEILQDSD
                       ************************************************************

MYC_TROSSULUS          FF
MYC_CANCER             FF
MYC_EDULIS             FF
                       **

ALIGNMENTS FBXW7-GENE_AMINO ACID SEQUENCES

FBXW7_CANCER                 MASKKIDINKASVEDFEKVNGIGLRKAKAIIKYRKSFGGFSTVKDLSSVP
FBXW7_TROSSULUS              MASKKIDINKASVEDFEKVNGIGLRKAKAIIKYRKSFGGFSTVKDLSSVP
FBXW7_EDULIS                 MASKKIDINKASVEDFEKVNGIGLRKAKAIIKYRKSFGGFSTVKDLSSVP
                             **************************************************

FBXW7_CANCER                 GVGDCIVEQSRTQLTCSNRKRKAVRTPKVKDNVCREQPSSSGLSARSIRH
FBXW7_TROSSULUS              GVGDCIVEQSRTQLTCSNRKRKAVRTPKVKDNVCREQPSSSGLSARSIRH
FBXW7_EDULIS                 GVGDCIVEQSRTQLTCSNCKRKAVRTPKVKDNVCKEQPSSSGLSARSIRH
                             ****************** ***************:***************

FBXW7_CANCER                 NQRALFKSTEGNVDDCNSIVCQEGHMSEEESGDEVMFREEDKSNAQHKSR
FBXW7_TROSSULUS              NQRALFKSTEGNVDDCNSIVCQEGHMSEEESGDEVMFREEDKSNAQHKSR
FBXW7_EDULIS                 NQRALFKSTEEHVDDGNSIVSHEGHMSEEESGDEVMFREEDRSNAQHKSR
                             ********** :*** ****.:*******************:********

FBXW7_CANCER                 IPIPVNGLFSSHVMKRKGHVVKEEPDSSPGKKFCRDGDYQSTFVNLQKSS
FBXW7_TROSSULUS              IPIPVNGLFSSHVMKRKGHVVKEEPDSSPGKKFCRDGDYQSTFVNLQKSS
FBXW7_EDULIS                 IPIPVNGLFSSHVMKRKGHVVKEEPDSSPGKKFCRDGDYQSTFVNLQKSS
                             **************************************************

FBXW7_CANCER                 RQNLIATQRMRIPSKEHPPDKLAEWLQIFQNWGNAERLMALDELITFCDP
FBXW7_TROSSULUS              RQNLIATQRMRIPSKEHPPDKLAEWLQIFQNWGNAERLMALDELITFCDP
FBXW7_EDULIS                 RQNLIATQRMRIPSKEHPPDKLAEWLQIFQNWGNAERLMALDELITFCDP
                             **************************************************
```

```
FBXW7_CANCER                 TQVRHMMQVIEPQFQRDFISLLPKELALYVLSFLEPKSLLKAAQTCRYWR
FBXW7_TROSSULUS              TQVRHMMQVIEPQFQRDFISLLPKELALYVLSFLEPKSLLKAAQTCRYWR
FBXW7_EDULIS                 TQVRHMMQVIEPQFQRDFISLLPKELALYVLSFLEPKSLLKAAQTCRYWR
                             **************************************************
```

```
FBXW7_CANCER                 VLAEDNLLWREKCREESIDDNLVYGSNRLRRRHTSRCPWKSLYMRQHQIE
FBXW7_TROSSULUS              VLAEDNLLWREKCREESIDDNLVYGSNRLRRRHTSRCPWKSLYMRQHQIE
FBXW7_EDULIS                 VLAEDNLLWREKCREESIDDNLVYGINRLRRRHTSRCPWKSLYMRQHQIE
                             ************************* ************************
```

```
FBXW7_CANCER                 LNWRTGEIPGPKLLKGHDDHVITCLEFCGTRIVSGSDDNTLKVWSAITGK
FBXW7_TROSSULUS              LNWRTGEIPGPKLLKGHDDHVITCLEFCGTRIVSGSDDNTLKVWSAITGK
FBXW7_EDULIS                 LNWRTGEIPGPKLLKGHDDHVITCLEFCGTRIVSGSDDNTLKVWSAITGK
```

**************************************************

```
FBXW7_CANCER                 CLRTLVGHTGGVWSSQMSNSVVISGSTDRTLKVWNADTGQCIHTLYGHSS
FBXW7_TROSSULUS              CLRTLVGHTGGVWSSQMSNSVVISGSTDRTLKVWNADTGQCIHTLYGHSS
FBXW7_EDULIS                 CLRTLVGHTGGVWSSQMSNSVVISGSTDRTLKVWNADTGQCIHTLYGHSS
                             **************************************************

FBXW7_CANCER                 TVRCMHLHKNVVVSGSRDATLRVWDIDSGACLHVLMGHVAAVRCVQYDGR
FBXW7_TROSSULUS              TVRCMHLHKNVVVSGSRDATLRVWDIDSGACLHVLMGHVAAVRCVQYDGR
FBXW7_EDULIS                 TVRCMHLHKNVVVSGSRDATLRVWDIDSGACLHVLMGHVAAVRCVQYDGR
                             **************************************************

FBXW7_CANCER                 RVVSGAYDYMVKVWDPETETCLHTLQGHTNRVYSLQFDGIHIVSGSLDTS
FBXW7_TROSSULUS              RVVSGAYDYMVKVWDPETETCLHTLQGHTNRVYSLQFDGIHIVSGSLDTS
FBXW7_EDULIS                 RVVSGAYDYMVKVWDPETETCLHTLQGHTNRVYSLQFDGIHIVSGSLDTS
                             **************************************************

FBXW7_CANCER                 IRVWDVEAGNCLHTLIGHQSLTSGLELKDNILVSGNADSTVKVWDITTGQ
FBXW7_TROSSULUS              IRVWDVEAGNCLHTLIGHQSLTSGLELKDNILVSGNADSTVKVWDITTGQ
FBXW7_EDULIS                 IRVWDVEAGNCLHTLIGHQSLTSGLELKDNILVSGNADSTVKVWDITTGQ
                             **************************************************

FBXW7_CANCER                 CLQTLQGPNKHSSAVTCLQFNKKFVITSSDDGTVKIWDLKTGDFIRNLVA
FBXW7_TROSSULUS              CLQTLQGPNKHSSAVTCLQFNKKFVITSSDDGTVKIWDLKTGDFIRNLVA
FBXW7_EDULIS                 CLQTLQGPNKHSSAVTCLQFNKKFVITSSDDGTVKIWDLKTGDFIRNLVA
                             **************************************************

FBXW7_CANCER                 LDSGGSGGVVWRVRCSNTKLVCAVGSRNGTEETKLHVLDFDVDEKKCLY
FBXW7_TROSSULUS              LDSGGSGGVVWRVRCSNTKLVCAVGSRNGTEETKLHVLDFDVDEKKCLY
FBXW7_EDULIS                 LDSGGSGGVVWRVRCSNTKLVCAVGSRNGTEETKLHVLDFDVDEKKCLY
                             *************************************************
```
